# Intercontinental genomic parallelism in multiple adaptive radiations

**DOI:** 10.1101/856344

**Authors:** Isabel S. Magalhaes, James R. Whiting, Daniele D’Agostino, Paul A. Hohenlohe, Muayad Mahmud, Michael A. Bell, Skúli Skúlason, Andrew D.C. MacColl

## Abstract

Parallelism, the evolution of similar traits in populations diversifying in similar conditions, provides good evidence of adaptation by natural selection. Many studies of parallelism have focused on comparisons of strongly different ecotypes or sharply contrasting environments, defined *a priori*, which could upwardly bias the apparent prevalence of parallelism. Here, we estimated genomic parallelism associated with individual components of environmental and phenotypic variation at an intercontinental scale across four adaptive radiations of the three-spined stickleback (*Gasterosteus aculeatus*), by associating genome-wide allele frequencies with continuous distributions of environmental and phenotypic variation. We found that genomic parallelism was well predicted by parallelism of phenotype-environment associations, suggesting that a quantitative characterization of phenotypes and environments can provide a good prediction of expected genomic parallelism. Further, we examined the explanatory power of genetic, phenotypic, and environmental similarity in predicting parallelism. We found that parallelism tended to be greater for geographically proximate, genetically similar radiations, highlighting the significant contingency of standing variation in the early stages of adaptive radiations, before new mutations accumulate. However, we also demonstrate that distance within multivariate environmental space predicts parallelism, after correction for genetic distance. This study thus demonstrates the relative influences of environment, phenotype and genetic contingency on repeatable signatures of adaptation in the genome.

## Introduction

Adaptive radiations are well known as rapid branchings on the tree of life (^1^).They may be the source of most biodiversity, and their study has revealed a great deal about the evolution of phenotypic diversity (^1,2^). However, patterns in adaptive radiations also highlight some of the unknowns about how biodiversity evolves. For example, although adaptive radiations are typified by abundant phenotypic diversity, not all trait combinations evolve in every radiation, while on the other hand organisms in different places sometimes arrive at very similar endpoints (^3,4^). This suggests that Stephen Jay Gould’s famous contention that evolution is contingent and unrepeatable (^5^) cannot be completely true. A good deal of work has focussed on the role that genetic correlations between traits might play in creating constraints on diversity, but so far the answers provided by this approach have not been entirely satisfactory (^3,6,7^). Alongside these processes, it is probable evolution can be shaped in a predictable way by common environments and shared selective regimes, and that the emergence of repeatable patterns of evolution, a process predominantly known as parallelism (which we distinguish from convergence here by the inclusion of shared evolutionary ‘start’, as well as ‘end’, points, although see (^8,9^), is the direct result of environmental similarities within and between radiations. Striking examples of phenotypic parallelism (^1,10^) in the natural world support this hypothesis, and the persistent appearance of familiar forms in similar ecological niches demonstrates the significance of selection in this process.

However, a significant problem with studies that have focused on phenotypic parallelism is that they have concentrated largely on the comparison of pairs of strongly different ecotypes or widely different environments (^11–13^). Such studies could upwardly bias the apparent prevalence of parallelism because the chosen comparisons were known *a priori* to occur repeatedly in different locations, effectively constraining the evolutionary end point. The approach also conceals the role of individual components of environmental variation in driving parallelism, as similar environments are often assumed based on comparable phenotypes. This gap surely needs addressing for a complete understanding of adaptation (^14^). In addition, traits that are measurable in sufficient numbers from (usually) wild organisms are generally limited to morphological and life history traits (^13^). This seriously compromises our ability to understand adaptation, a great deal of which is likely to be physiological in the broad sense. Such drawbacks highlight the importance of combining measures of phenotype and environment alongside genomics in studies of parallelism. The detection of consistent genomic signatures across multiple, independent natural populations has proven to be a valuable tool for studies of evolutionary patterns and the discovery of genes involved in adaptation (^15–17^). However, yet again our comprehension of the relationship between genomic parallelism and continuous phenotypic or environmental variation is surprisingly poor.

Until recently, a major barrier to combining genotype, environment and phenotype has been the high costs of sequencing, prohibiting large-scale genomic sampling that could be aligned with large-scale ecological sampling. However with dramatic drops in DNA sequencing costs, such genomic data can nowadays be used alongside ecological data to associate allele frequencies (^18^), and determine genomic regions associated with individual components of environment and phenotypes. Such an approach is essential if studies of parallelism are to shift from description to hypothesis testing, but has rarely been applied (but see ^12,19,20^), and it remains to be shown whether signals of parallelism obtained from continuous measures are comparable to those from ecotypes and from previous studies. Here we make use of such methods to test for environmentally and phenotypically associated genomic parallelism across radiations of three-spined stickleback fish (*Gasterosteus aculeatus*, hereafter ‘stickleback’).

Stickleback provide a powerful natural experiment to test parallelism. They are primitively marine but have undergone replicated adaptive radiations across the northern hemisphere following colonisation of freshwater in widely separated geographical locations. This allows us to compare multiple populations derived from the marine ancestor, and provides a model for exploring both phenotypic parallelism and its genetic basis (^21,22^) in response to environmental variation. Phenotypic parallelism in populations that have evolved independently in similar habitats is well established (^23,24^), and whilst often considered in dichotomous pairings of marine – freshwater, benthic – limnetic, lake -stream ecotypes, there is a huge amount of continuous phenotypic variation among freshwater populations that has hitherto rarely been explored (^25^) in this context. In addition, genome scans have identified loci that have come repeatedly under selection across the contrasting ecotype pairs (^17,26,27^), but the combination of the phenotypic and genomic parallelism in one study is rare (although see ^28^) and has not previously been done at the scale of replicated adaptive radiations across continents.

For this study we sampled 73 freshwater lake populations from four adaptive radiations in Alaska, British Columbia (‘BC’), Iceland and the island of North Uist (‘Scotland’) (Fig. 1, Supplementary Table 1), measured a set of six biotic and abiotic environmental variables and a set of 12 phenotypic traits (measures of body shape, armour traits and gill rakers) (Supplementary Table 2), and performed a genome-wide scan of a total of 1,304 individual stickleback using restriction-site associated DNA (RAD) sequencing. We first assessed environmental and phenotypic parallelism among radiations, and their phylogenetic relationship, which provided a reference for how much genomic parallelism to expect. We then scanned the genome for associations with continuous environmental and phenotypic axes of variation within radiations, and identified genomic parallelism by looking at the presence of allele frequency associations in the same genomic regions across radiations. As marine-freshwater parallelism is well-documented (^17,27,29^), we compared our results for parallelism across freshwater radiations with well-studied marine-freshwater parallelism in this species, and used the results as a positive control for the methods used. Finally, we examined how the prevalence of parallel genomic regions is associated with the phylogenetic histories of our adaptive radiations and how well it can be explained by multivariate quantification of environmental and phenotypic similarity between radiations. Rewards to be gained by connecting the evolution of parallelism more explicitly to the environmental and phenotypic variation include a better grasp of why some traits evolve in concert and a predictive understanding of parallelism and repeatability (^4^). This new understanding is essential if we are to reach a consensus on how biodiversity is altered by adaptation.

**Fig. 1.**
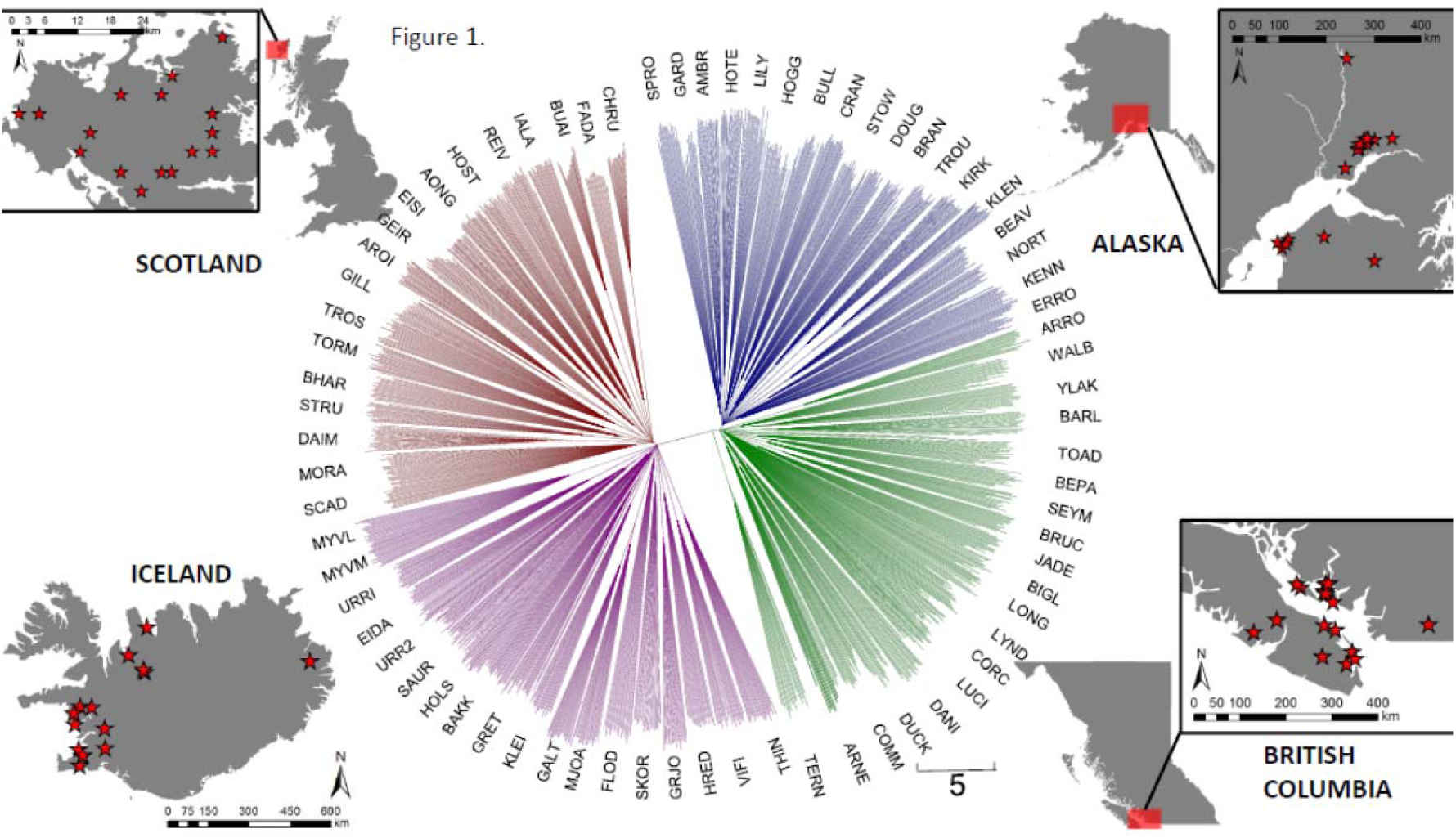
Sampling sites and bootstrapped NJ tree for stickleback from 73 freshwater populations from four countries on two continents, based on 8,395 genetic markers for 1,304 individuals. All nodes shown have bootstrap support of at least 80 (other nodes were collapsed). Branches are coloured by the radiation to which they belong. Tips represent individual fish, which were generally tightly clustered by population (small labels). Stars represent lakes sampled.

### Environmental and phenotypic similarity across radiations

Interpreting inter-radiation patterns of genomic parallelism associated with phenotypic traits and environmental variables requires that we first assess the environmental and phenotypic similarity across all four radiations analysed here, as well as the relationship between the two types of variables. Environmental similarity is predicted to influence both phenotypic and genomic parallelism, as similar selection regimes are imposed on organisms (^13,30,31^). These analyses provide insights into some major factors that are likely to influence the emergence of a common pattern of genomic divergence at such a large geographical scale, and indicate how much genomic parallelism associated with environments and phenotypes to expect.

#### I) Environment

Adaptation of stickleback to freshwater in these radiations has happened rapidly, over the course of the past few thousand years, and has most likely been driven by strong selection favouring pre-existing alleles present at low frequencies in the ancestral marine population (^32^). Therefore, we might expect to observe shared features of diversification when multiple groups of organisms experience similar environmental circumstances, as similar selection regimes are imposed on organisms (^12,33^). A Principal Component Analysis (PCA) on six measures of water chemistry (pH, calcium (Ca), sodium (Na) and zinc (Zn), and parasitism (prevalence of the parasites *Gyrodactylus spp*. (Gyro) and *Schistochephalus solidus* (Schisto)) across all lakes revealed that the first axis of environmental variance (Env_PC1_) separated lakes along a gradient of pH and calcium concentration (Fig. 2; Supplementary Table 3). This axis did not separate radiations but emphasised the variation from alkaline to acid present in all of them, with the widest range in Alaska. The second axis of environmental variance (Env_PC2_) distinguished mostly between lakes with high and low zinc and largely separated the European from the North American lakes. The two groups of European lakes overlapped very little environmentally with each other along the second axis or with the North American lakes, but the variation in BC lakes was completely subsumed within that of the more environmentally variable Alaskan lakes.

**Fig. 2.**
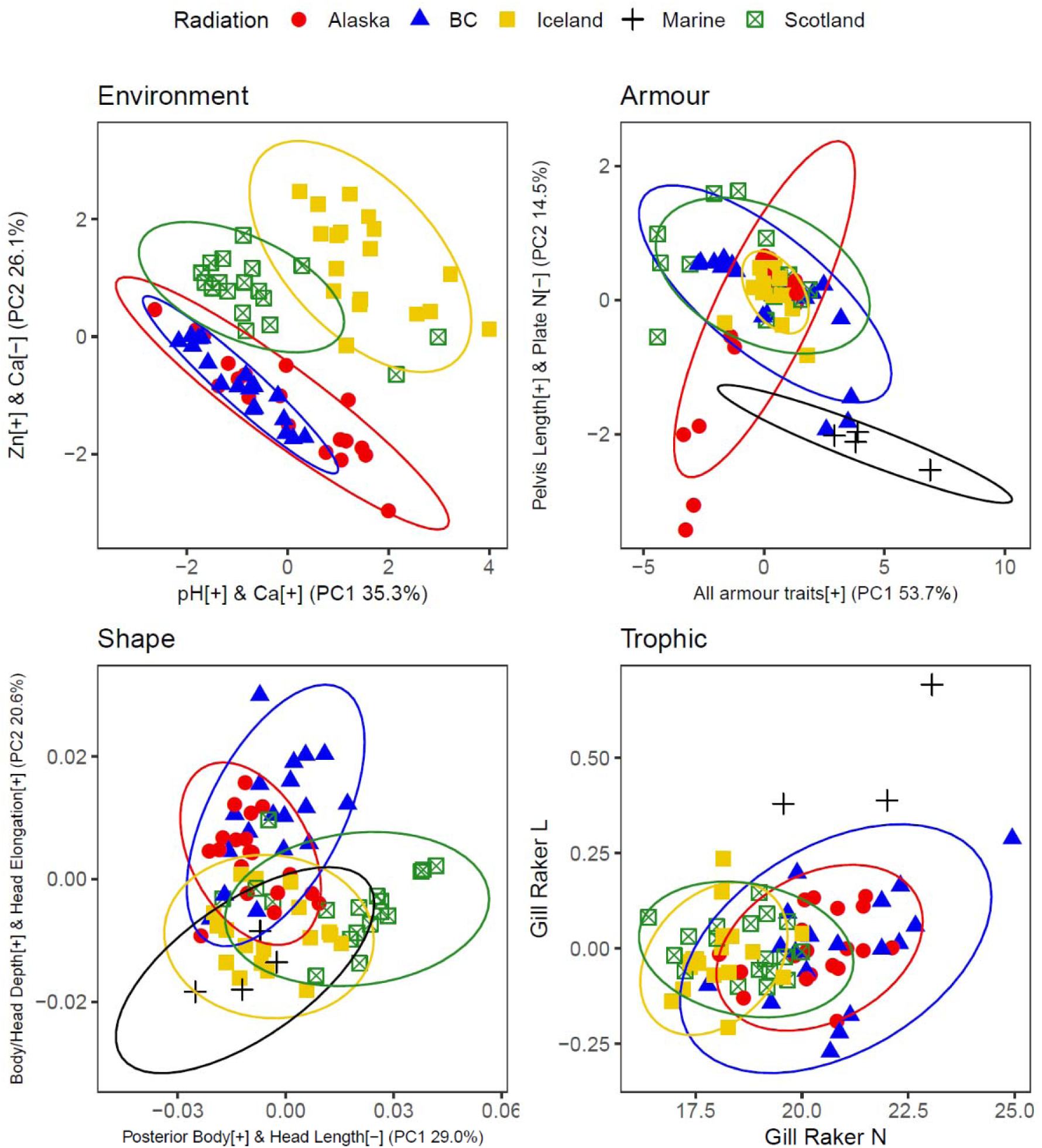
Principal Component Analyses of environmental variables (Environment); regression residuals of Procrustes coordinates against log centroid of body shape (Shape); armour traits (Armour: length of dorsal spines 1 (DS1) and 2 (DS2), length of pelvic spine (PS), length and height of pelvis (LP and HP), length of biggest armour plate (BAP) and number of armour plates (Plate_N); and gill raker numbers (Gill_Raker_N) vs gill raker length (Gill_Raker_L) (Trophic). Each point represents a population and ellipses are 95% confidence ellipses. Names of variables with the highest positive (+) or negative (-) loadings on each axes are on legends of each axes. All loadings of variables on the first 3 PCs are in Supplementary Table 3. Marine populations (+) are projected where data was available using PC loadings calculated with freshwater populations only.

An analysis of the similarity of the direction of the major PC vectors (θ) of environmental variation (PCs that explained at least 10% of the variation) and of the magnitude of their variation revealed that Alaska and BC exhibited the strongest environmental similarity in terms of direction (θ = 8.71°, *p* = 0.077) (Supplementary Table 4), as lower values of θ are indicative of more common environmental covariance. However, they also showed the greatest difference in magnitudes of variation, as BC had the lowest amount of environmental variation of all radiations while Alaska had the highest. Iceland on the other hand was the most unique radiation in terms of environmental covariance, a pattern likely driven by Iceland’s uniquely volcanic nature.

#### II) Phenotypes – Armour

The major axis of armour variation (Armour_PC1_) represented a correlated axis of all traits, particularly dorsal spine length and pelvic characteristics (Fig. 2; Supplementary Table 3). Although there were significant differences in Armour_PC1_ among both radiations and lakes (Supplementary Table 5), the direction of this vector was highly conserved across all radiations (4.13° ≤ θ ≤ 8.37°; 0.004 ≤ *p* ≤ 0.053) (Supplementary Table 4). Nonetheless, we found populations with extreme armour, such as absence or extreme reduction of dorsal spines, pelvic spines, pelvis size, and generally low Armour_PC1_, that occur in Scotland and Alaska but not elsewhere. However, while in Scotland pelvic armour reductions were accompanied by complete or almost complete loss of armour plates (high Armour_PC2_), Alaskan populations retained those even when other features of armour were highly reduced. These deviations produce the somewhat anomalous relationship between Armour_PC1_ and Armour_PC2_ in Alaska (Fig. 2). Iceland exhibited particularly low variation in armour traits compared to other radiations, while BC had the highest. These results indicate that whilst the direction of armour variation occurred predominantly on a shared axis across adaptive radiations, the amount of armour variation along that axis was variable. When comparing all freshwater populations with four marine ones (one from each country), we found that, aside from a few populations from BC, there was no overlap in armour traits between marine and freshwater populations. This is because marine populations have a higher number of lateral plates and more exaggerated armour traits in general. Importantly however, projection of marine armour phenotypes suggests they fall on the same axis but beyond freshwater space (Fig. 2).

#### III) Phenotypes -Body shape

Despite some overlap in body shape across all radiations suggesting similar morphologies have evolved repeatedly across continents (Fig. 2), there were significant differences in body shape morphologies among both radiations and lakes (Supplementary Table 5). The most extreme body shapes were found in Scotland, where some populations have very elongated and slender bodies with small heads (Shape_PC1_ axis), and in BC where some populations had the deepest bodies and heads and the longest heads (Shape_PC2_ axis). Scotland had the largest amount of variation in body shape, which is a surprising result given this variation is found within 1000 km^2^ of North Uist, a much smaller area than the other three radiations. The orientation of the major PC vectors (θ) revealed that North American radiations were the most similar (θ = 3.49°, p=0.19), followed by the European radiations (θ = 8.48°, p=0.63) (Supplementary Table 4), which highlights a potential effect of geography/shared ancestry on the major axis of shape variation. Marine populations overlapped in body shape with some of the freshwater populations that had lower ShapePC1 and ShapePC2 scores, most of them found in Iceland (Fig. 2).

#### IV) Phenotypes - Trophic morphology

The largest variation in gill raker numbers and length was found in Alaska, while BC populations contained a subset of that variation (Fig. 2). Across the two dimensions of gill raker number and length, variation within radiations was generally constrained to a major axis in which populations with more gill rakers also tended to have longer gill rakers. All freshwater populations generally had shorter gill rakers for their size relative to marine populations.

#### V) Relationship between environmental and phenotypic similarity

We used GLMs to test for parallel associations between the first two environmental PCs (Env_PC1_ and Env_PC2_) and armour, shape, and gill raker morphology across radiations. We found that Armour_PC1_ and Armour_PC2_ were also significantly associated with Env_PC1_ (Armour_PC1_: F=6.13, p=0.016; ArmourPC2: F=4.11, p=0.047), but not with Env_PC2_ (Armour_PC1_: F= 1.12, p=0.293; Armour_PC2_: F=1.10, p=0.298) (Supplementary Table 6). Armour_PC1_ correlations with Env_PC1_ were similar across radiations but significant slope variation was observed between radiations for Armour_PC2_ and Env_PC1_ associations (F= 2.76, p=0.050). The result highlights parallelism in the way general skeletal traits (Armour_PC1_) are reduced with lower pH and calcium (Env_PC1_), but non-parallel associations between these measures and the relationship between pelvic size and plate number (Armour_PC2_). The latter is consistent with previous findings of a lack of convergence in plate number in response to Ca between North American and European radiations (^34^), while the former suggests that common pH and calcium environments, experienced in all radiations as a shared acid-alkali axis, promotes phenotypic parallelism.

Shape_PC1_ and Shape_PC2_ were significantly associated with both Env_PC1_ (Shape_PC1_: F= 4.85, p=0.031; Shape_PC2_: F=28.20, p<0.001) and Env_PC2_ (Shape_PC1_: F= 7.56, p=0.008; Shape_PC2_: F=48.01, p<0.001), suggesting body shape variation is strongly linked to specific measures of water chemistry. Intercepts across radiations varied significantly but slopes did not, indicating parallelism across radiations in the way shape and environment are associated (Supplementary Table 6). The result highlights parallelism in the way bodies become more elongated and slender and heads are reduced with lower pH and calcium and higher Zinc.

As for gill rakers, we expected trophic morphology to evolve in response to zooplankton communities, which in turn have been shown to vary in response to measures of water chemistry (^35,36^), in particular Zn which was the dominant component of Env_PC2_. Consistent with this, we found gill raker number to be strongly significantly associated with both Env_PC2_ (F=43.74, p<0.001) and Env_PC1_ (F=18.17, p<0.001). Slopes varied between radiations for the association of gill raker number and Env_PC1_, demonstrating non-parallelism in trophic response to pH and calcium. In contrast, common slopes across radiations highlight parallelism in trophic association between gill raker number and Zn variation (Env_PC2_).

Corroborating predictions that selection imposed by similar environments can lead to phenotypic parallelism (^1^), we found that similar variation from alkaline to acid environments (Env_PC1_) present in all radiations was significantly associated with parallel variation in body shape and armour. If phenotypic adaptation happens via natural selection on genetic variation, we would then expect these associations to translate into genomic parallelism associated with both environmental variables that show a similar range across radiations (pH and Ca) and phenotypic traits associated with them.

### Phylogenetic relationship among radiations

The probability of genomic parallelism has been linked to time since lineages split, highlighting the probable influence of genetic similarity and shared ancestral variants on parallelism (^16^), it is therefore important that we know the genetic relationship across all populations before interpreting patterns of genomic parallelism. Genomic parallelism across populations within any one radiation is likely to evolve from shared genetic variation, but the probability of that should decline in geographically distant radiations as a function of common ancestry (^16,37^). Indeed, recent studies have highlighted the probable limitation of parallelism in this system across continental scales (^20,38,39^). A Neighbour Joining (NJ) tree based on 8,395 unlinked SNPs showed that, with the exception of a likely recently formed population in Alaska (TERN), which is basal to both the Alaskan and British Columbian radiations, the four geographic locations form four well-resolved radiations (Fig. 1). A PCA on the same dataset confirmed that radiations form independent clusters, and also revealed that the dominant axis of variation (PC1 = 36.0%) separates North American and European radiations pairs from one another (Supplementary Fig. 1). North American radiations then separated on PC2 (7.0%) and European radiations on PC3 (5.8%). In addition to being close together in the NJ tree, we found that the geographically adjacent radiations were also the most genetically similar (Supplementary Table 7): Alaska and BC (mean pairwise F_ST_ = 0.198), and Scotland and Iceland (F_ST_ = 0.194), suggesting that although these form independent clusters, the lineage split between them is relatively recent, or that any gene flow between radiations is occurring within the Atlantic and Pacific groups. Genetic divergence between inter-continental pairings was found to be stronger and deeper (0.314 ⍰ F_ST_ ⍰ 0.338) than within continents, which is consistent with previous studies that have estimated the time of divergence between stickleback from Europe and North America to be approximately 200,000 years (^40,41^ but see ^42^). Further, between-continent structuring accounts for the largest proportion of molecular variance in our data (AMOVA: σ = 889.7, 34.7%). Within continents, populations within radiations (σ = 366.5, 14.3%) were more genetically variable than radiations (σ = 142.3, 5.6%) (Supplementary Table 8). This highlights that molecular variance isn’t structuring according to geographic scale (Continent > Radiation > Populations), but rather gene flow may be occurring between intra-continental radiations.

Following the idea that the probability of parallelism at the genetic level is linked to time since lineages split (i.e. genetic similarity), we would expect groups with closer evolutionary histories, or indeed present gene flow, to show the strongest genomic parallelism if adaptation occurs through shared standing genetic variation or the exchange of beneficial freshwater alleles between intra-continental radiations via marine populations (^29,32^).

### Phenotypically and environmentally associated SNPs and genomic regions within radiations

We scanned the genome to identify regions associated with individual components of phenotypic and environmental variation within each of the four adaptive radiations. We focused on repeated changes within the same genomic regions, rather than on reuse of the same mutations. This is because the causal mutations are unknown in most cases and may not be sequenced by reduced representation sequencing methods such as RAD-sequencing. For each radiation separately (N=18 or 19 populations) we used Bayesian linear models implemented in Bayenv2 (^43^) to identify associations between population SNP allele frequencies (N=10 to 21 individual fish, mean= 17.8), the set of six biotic and abiotic environmental variables and the set of 12 phenotypic traits (Supplementary Table 2). As a positive control for the methods used we compared our results for parallelism across freshwater radiations with well-studied marine-freshwater parallelism in this species (^17,27,29^), and then examined genomic differentiation between all freshwater populations pooled within a radiation and four marine populations pooled together (one from each country, Supplementary Table 1).

Our analyses identified population allele frequencies of several thousand SNPs for each radiation as being highly correlated (high Bayesfactor [>log_10_(1.5)] and top 5% of Spearman’s ρ, see methods) with the abiotic and biotic environmental variables (‘environmentally associated SNPs’) or with phenotypic traits (‘phenotypically associated SNPs’ (Supplementary Table 9). SNPs were then mapped onto non-overlapping sliding windows of 50kb, 75kb, 100kb or 200kb, which allowed us to test the robustness of our results across different extents of linkage. Further, we repeated our analysis across windows of equivalent genetic distance (0.1 cM windows), which confirmed that our results were not influenced by variable linkage across the genome (Supporting Information). Here we report only results for 50kb windows, given that this is consistent with approximate linkage disequilibrium within the stickleback genome (^44,45^). Results for window sizes of 75kb, 100kb, 200kb can be found in Supplementary Dataset 1. Our 50kb dataset was composed of 4,868 windows with SNPs in all locations, covering approximately 55% of the 447 Mb genome, with a further 1,940 windows sequenced in 2 or more locations providing information on an additional 21.7% of the genome.

Windows were classified as ‘environmentally associated’ and ‘phenotypically associated’ if they contained more associated SNPs than expected under a 99% binomial expectation (^46^). We found 1791 unique 50kb windows associated with an environmental variable or a phenotypic trait (Supplementary Dataset 1), ranging from 146 windows associated with pelvic spine length in BC, to 21 associated with Na variation, also in BC. It is striking how many windows show strong signals of association with phenotypic and environmental variables, even when their variation is modest, clearly supporting the adaptive nature of these radiations.

Out of all the unique windows, 431 were associated with both an environmental and phenotypic variable in the same radiation, suggesting several regions associated with phenotypic traits might also be responsible for local adaptation to environments (Supplementary Dataset 1). It also suggests that directly measuring important aspects of the environment may provide profitable ways of discerning the genomic basis of adaptation and of identifying agents of selection, while bypassing the often-difficult measurement of phenotypes.

### Genomic parallelism associated with environmental and phenotypic variation across radiations

To quantify genomic parallelism we identified environmentally- and phenotypically-associated windows that were shared across two or more of our radiations, i.e. parallel windows (Supplementary Fig. 2). We then compared the overall observed numbers of parallel windows to a null distribution of randomly associated windows permuted over 10,000 iterations.

We quantified genomic parallelism at three levels: 1) gross, general levels of parallelism associated across all radiations for groups of phenotypes or environment; 2) genomic parallelism associated with individual variables across all comparisons to understand contributions of individual variables; 3) parallelism for individual variables in specific pairings, to identify pairs of radiations having the highest levels of parallelism, and for which variables. The latter is important as patterns of parallelism may be lost when we pool more than two radiations or variables together, if for example parallelism is very strong in one radiation-pairing but not others.

When quantifying the overall level of genomic parallelism associated with groups of environmental or phenotypic variables we found no environmentally- or phenotypically-associated 50kb windows parallel in all four radiations for individual variables (Randomised permutations N_Expected-Environmental_= 0.0002, N_Exp-Pheno_ = 0.0001), but one window was parallel in a group of three radiations (chrIV: 14400000-14450000 associated with length of pelvis in BC, Iceland and Scotland) (N_Exp-Env_ = 0.05, N_Exp-Pheno_ = 0.161, *p* = 0.149). Many windows however exhibited parallelism between pairs of two radiations. A total of 39 environmentally associated windows (pooled across all 6 environmental variables) (N_Exp_ = 11.9, 95% Upper limit (UL) = 18, *p* < 0.001) and 65 phenotypically associated windows (pooled across all 12 phenotypic variables) (N_Exp_ = 30.9, 95% UL = 40, *p* < 0.001) were parallel between two radiations (Supplementary Fig. 2). Parallelism was disproportionately greater for armour and gill raker traits (number mostly) than for shape, with the 65 phenotypically associated windows split into 46, 12 and seven associated windows for armour, gill raker and shape variables respectively (χ^2^= 6.506, P = 0.04). This is consistent with the fact that skeletal traits, several of which are known to have simple genetic architectures (^26,47,48^), are particularly likely to show evidence of phenotypic parallelism. Interestingly, parallel associated windows (mean SNP N = 7.13) had on average more SNPs per window than non-parallel windows (mean SNP N = 6.27) (GLM, LRT_1,3867_ = 22.1, *p* < 0.001) and exhibited slightly stronger signals of association with variables (mean residual SNPs above expected = 1.82 parallel; 1.61 non-parallel; GLM, LRT_1,3867_ = 5.05, *p* = 0.025).

Random permutations indicated that there were statistically significant levels of parallelism for the number of windows (‘significantly parallel windows’) associated with two environmental variables (Ca and pH) (Fig. 3; Supplementary Table 10). Consistent with our expectations from patterns of environmental parallelism, these were the same variables that share an axis of variation (Env_PC1_) across freshwater environments in all radiations. In addition, we did not detect significant genomic parallelism associated with variables that exhibit variation between radiations, such as salinity, zinc and S. solidus prevalence. These results together confirm that common environmental axes, such as the one experienced in all radiations as a shared acid-alkali axis, likely promote signals of parallelism in the genome.

**Fig. 3.**
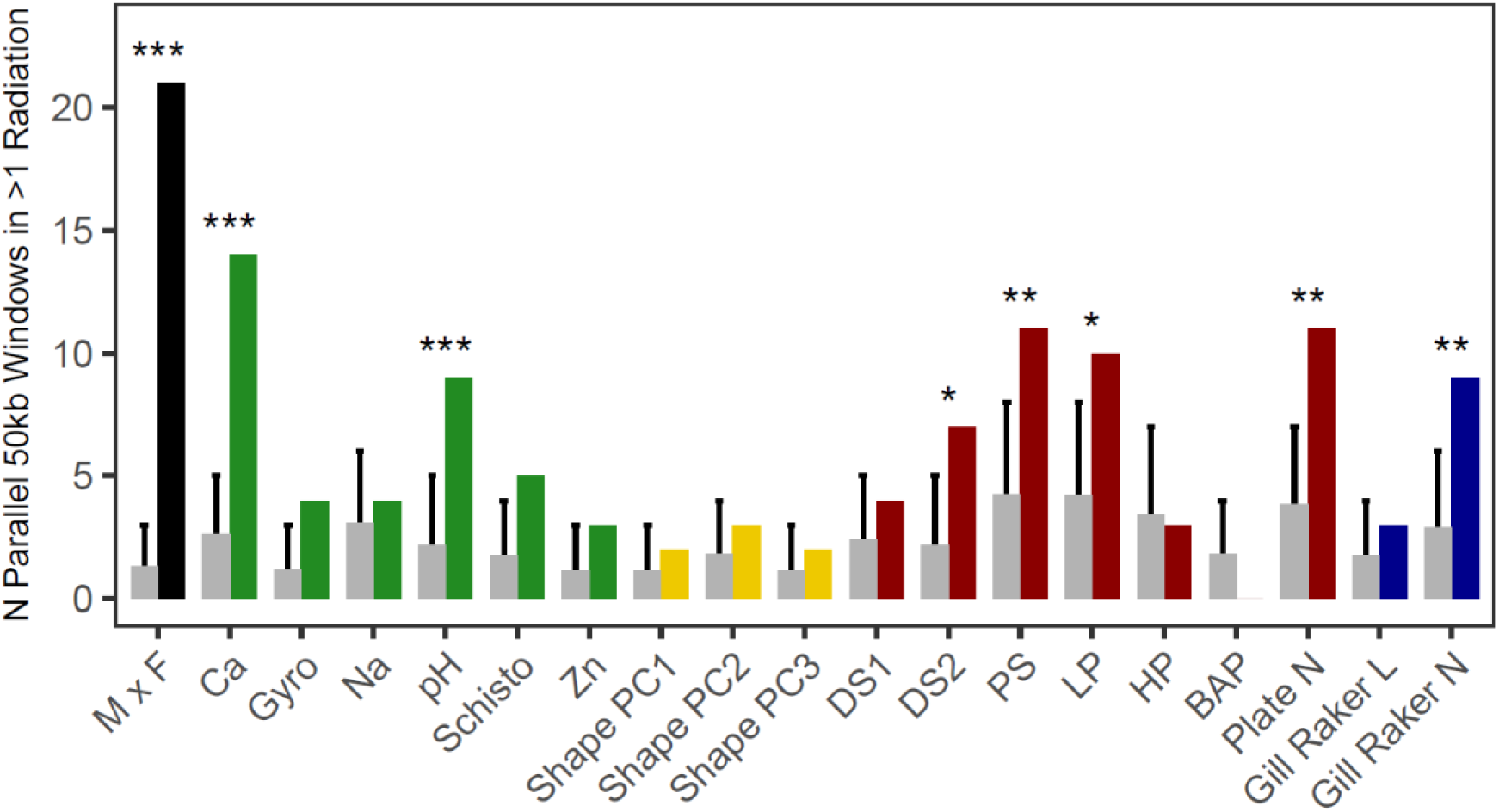
Expected and observed counts of 50kb windows containing an above 99% binomial expectation number of SNPs associated with marine x freshwater (MxF), environmental variables and phenotypic traits in at least 2 radiations. Expected bars (grey) represent mean counts across 10,000 simulated outcomes with 95% confidence intervals per a one-tailed hypothesis. Asterisks denote significance of FDR-corrected one-tailed tests between the observed counts and the 100,000 simulated counts at the <0.05 (*), <0.01 (**) and <0.001 (***) levels.

We also found more genomic parallelism than expected by chance associated with a total of five phenotypic variables (Fig. 3; Supplementary Table 10): four armour traits (2^nd^ dorsal spine, pelvic spine length, length of pelvis and armour plate number) and gill raker number. These results are consistent with the high heritability of skeletal traits (^26,47,48^), and suggest that variation in armour is the result of different genotypes being selected in different environments. Further, parallel QTLs have been described for gill raker number (^49^), but not length, which exhibits more plasticity (^50^). Body shape traits were not associated with any significant genomic parallelism, despite parallelism across radiations in the way shape is associated with the environment. It is probable the partly plastic nature of body shape (^51,52^) leads to an association between environment and body shape via the reaction norm rather than genomic re-use.

Our analyses also showed much stronger parallelism across marine-freshwater (MxF) comparisons than across freshwater variables (Fig. 3), and 44/158 of MxF associated windows overlapped with previously identified genomic regions contributing to marine-freshwater divergence (^17,44^) (Supplementary Table 11). Several regions found parallel for MxF analyses were also parallel for Ca, pH, Na, armour traits and gill raker number (Supplementary Table 12). These results suggest our methods and sequencing coverage are robust enough to recover known parallel regions, and interestingly that genomic parallelism associated with freshwater variables is more modest than marine-freshwater parallelism. The latter likely reflects subtler variation between habitats within radiations in comparison to stark marine-freshwater contrasts. Further, these results highlight that binary ecotype pairings, which likely include variation in many environmental and phenotypic traits, lump together parallelism of many components of fitness without being able to discern which are parallel and which are non-parallel. Overall, our results suggest that across these four freshwater adaptive radiations, evolution of these phenotypes and environmentally associated traits are disproportionately linked to the same genomic regions.

Within specific pairings, we found the greatest number of significantly parallel windows in the comparison between Alaska and BC (two environmental variables: Ca and *Gyrodactylus spp*., and three phenotypic traits: pelvic spine length, plate number and gill raker number), followed by Iceland and Scotland (two environmental variables: Ca and pH, and one phenotypic trait: dorsal spine length) (Supplementary Fig. 2; Supplementary Table 10). Two environmental variables and one phenotypic trait were also associated with significantly more parallel windows in comparisons between Iceland and the North American radiations: Ca and length of the pelvis (Alaska and Iceland) and *Schistocephalus solidus* (BC and Iceland). Pelvic spine length was the only variable of any category found to be parallel between Scotland and BC, and no significant parallelism was detected between Scotland and Alaska (suggesting observed overlap may be the result of chance). Phylogenetic patterns and the segregation of molecular variance strongly support the notion that radiations within continents share similar genetic variation, making parallelism through shared standing variation the most parsimonious explanation for our intra-continental parallelism biases. Experimental studies with stickleback have demonstrated rapid morphological adaptation from standing genetic variation, even recovering diverse morphologies from variation found within phenotypically-derived freshwater populations (^53^). Coancestry patterns, centred at the focal, causative loci, can discern between whether parallel evolution occurs on *de novo* mutations, standing variation, or introgressed alleles, however we lack the sequencing resolution in our current data to make these comparisons.

### Linkage and the genomic location of parallel regions

As with any reduced-representation genomic approach, the power of RAD sequencing to detect loci associated with phenotypic or environmental variables depends on linkage disequilibrium (LD) between markers and functional loci(^44,54,55^). The scale of LD varies widely across organisms and within genomes, but has been relatively well-characterized in stickleback (^56,57^), and RAD sequencing has been used successfully in this species specifically to test for genomic parallelism (^30,44,53,56,57^). One explanation for variable linkage across the genome is the negative relationship it shares with local recombination rate. To examine its influence on our results, we estimated recombination rate using a previously published genetic map (^31^) for our 50kb windows, and marked windows that were associated with any variable and associated windows that were also parallel across radiations. Recombination was significantly reduced in associated windows and parallel windows compared with non-associated windows (Supplementary Fig. 3; Kruskal-Wallis, *χ*^2^ = 121.43, *p* < 0.001), but did not differ significantly between outlier and parallel windows (*p* = 0.55). Reduced recombination can be an important process in adaptation through maintaining adaptive alleles, as has been demonstrated in this species (^58^) and others (eg.^59^). However, we cannot rule out that these patterns are produced by an increased ability to detect selection in low-recombination windows as a product of increased linkage with causative SNPs. The latter suggests that our estimates of association, and by extension parallelism, may be conservative if false-negatives are pervasive in high recombination regions. Importantly however, our signatures of parallelism cannot be explained by variable recombination.

Windows based on genetic distance (0.1 cM) corroborated 50kb results, returning strong signals of parallelism for calcium, pH, pelvic spine length, pelvis length, plate number and gill raker number. Interestingly, we also recovered weakly significant parallelism for several other environmental and armour variables (Supplementary Fig. 4, Supplementary Information), suggesting potentially stronger parallelism than we report at 50kb windows.

**Fig. 4.**
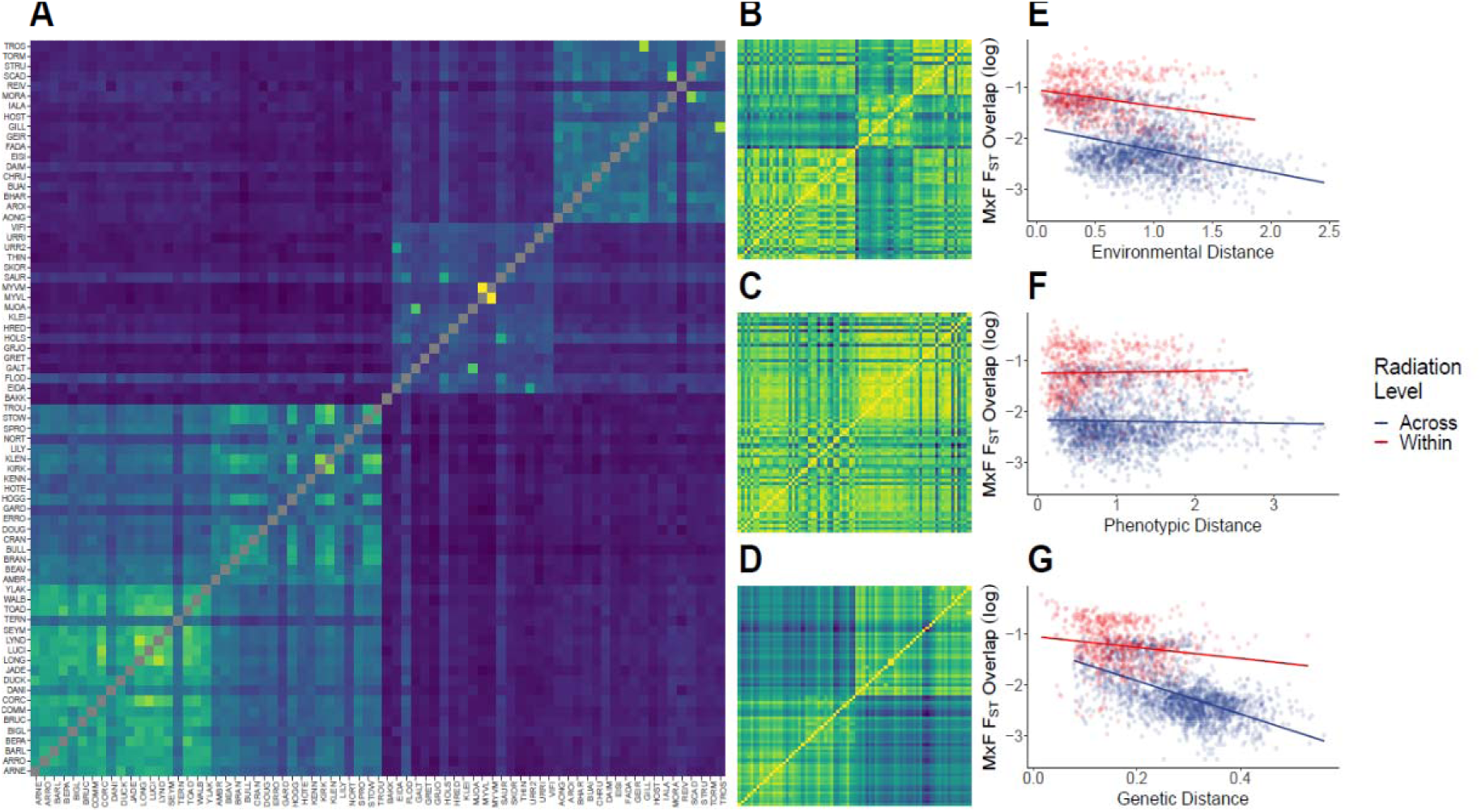
Associations between genome-wide marine - freshwater F_ST_ and environmental, phenotypic and genetic distance across all pairwise comparisons of 73 freshwater populations. A) Proportion of MxF F_ST_ 50kb outlier windows that overlap among freshwater replicates. Freshwater populations are ordered as Alaska, British Columbia, Iceland and Scotland, with these location distinguishable as four clear clusters. B) Environmental distance between freshwater populations, recorded as euclidean distance in PCA space for all 6 environmental variables. C) Phenotypic distance between freshwater populations, recorded as for environmental distance for the 12 phenotypic variables. D) Genetic distance between freshwater populations, recorded as genome-wide pairwise F_ST_ based on 8,395 unlinked SNPs. E-G) Associations between environmental (E), phenotypic (F) and genetic (G) distance and MxF F_ST_ overlap (log-transformed). Points are coloured according to whether pairwise comparison is being made within a radiation or across radiations.

To assess wider, linked parallel regions, we plotted 50kb windows across the genome to examine clustering of all associated windows (Supplementary Fig. 5). Adjacent windows (two or more, Supplementary Table 13) were pooled together to inspect putative causative genes (Supplementary Dataset 2). Using these two methods we identified a number of wider genomic regions that exhibited parallelism across multiple radiations. An example, and good positive control for our method, involves the pooling of windows associated with plate number in three radiations (Alaska, BC and Iceland) on chromosome IV around the well-known *Eda* gene, which has a well-established role in producing variation in the number of armour plates (^17,26,27^). This pattern emerges despite the limited variation in plate number across freshwater populations.

Pooling adjacent windows also identified a large cluster (250 kb) on chromosome I containing genes *igfbp2a, stk11ip* and *atp1a1*, and strongly associated with calcium, sodium and pH in several radiations (Supplementary Data 2). The clustering of windows in this region is perhaps unsurprising given it contains a known inversion (^60^), which as discussed should be beneficial for adaptive haplotypes by reducing local recombination (^61^). Within this region specifically, it is likely that the *atpa1a1* gene causes large effects on fitness, given its previously detected association with the major ecological transition from marine to freshwater (^62^) and functional role in metal ion management (^17,60^). Its apparent role in adaptation to much smaller cation variation between freshwater environments in this study is interesting in light of Fisher’s geometric model of adaptation, since this predicts that only alleles of smaller effect should fix as a fitness optimum is approached (^63^). Thus, it may be that different mutations in this gene have a spectrum of effect sizes, or that changes cause subtler differences in expression, rather than larger coding differences.

Our results support the existence of genomic regions of physically linked genes that are hitch-hiking in separate radiation pairs, and may contain genes that are parallel across all radiations but undetected by our genomic methods. Extensive linkage disequilibrium in freshwater populations is consistent with what is expected under strong directional selection after colonization from marine populations and has been reported for stickleback populations from Alaska (^44^), but it had not previously been observed for the same regions across several independent adaptive radiations. These results are also consistent with previous findings of large numbers of SNPs highly divergent between marine and freshwater stickleback aggregating in just 19 short genomic regions, including three known inversions (^60^), one of which we also detected and highlighted above.

### Relationships between genomic parallelism and phylogenetic, phenotypic and environmental similarity

Freshwater populations have radiated from marine common ancestors (^1,16^), thus the parallelism patterns described in this study are putatively the result of multiple marine - freshwater transitions. Based on this assumption, and using our SNP data from a marine population within each radiation, we performed genome-wide F_ST_ outlier analyses comparing each freshwater population with the marine population within that radiation. We then took the top 5% of 50kb windows according to F_ST_ in each marine – freshwater comparison and looked for overlapping outlier windows across all comparisons (N = 2,628). This quantified the extent of repeated genome-wide differentiation for MxF transitions within and across all radiations (Fig. 4a).

We then used Mantel tests to test statistically the effects on MxF genomic parallelism of relative genetic divergence and environmental/phenotypic similarity alongside one another. We compared the matrix of overlapping marine - freshwater outlier F_ST_ windows to equivalent matrices for environmental dissimilarity (Euclidean distances), phenotypic dissimilarity (Euclidean distances), and genetic dissimilarity (F_ST_ values) (Fig. 4). Across all population comparisons the number of parallel windows was strongly negatively correlated with genetic dissimilarity (*r* = -0.61, *p* < 0.001), and also negatively correlated with environmental dissimilarity (*r* = -0.42, *p* < 0.001) and phenotypic dissimilarity (*r* = -0.11, *p* = 0.022). These results highlight that genomic parallelism increases in populations that are more genetically, environmentally and phenotypically similar, though to varying extents.

Parallel windows were more common in intra-rather than inter-continental comparisons. This highlights the significance of our European and North American pairings as the major contributors towards pairwise signals of genomic parallelism, as we discussed previously, and strongly suggests that genomic parallelism at large geographic scales is contingent on shared genetic variation. Further, at an intra-continental scale, there exists the possibility of haplotype sharing between radiations by gene flow through marine populations, which may be facilitated in North America, despite the greater geographic distance, by a shared coastline connecting Alaska and BC (^64^). Recent research has also highlighted a probable genetic bottleneck in the founding of Atlantic marine populations that restricts shared freshwater alleles between Atlantic and Pacific freshwater populations (^39^).

We also conducted partial mantel tests for the effects of environmental and phenotypic similarity whilst controlling for genetic similarity, given this is likely to correlate with environment and phenotype in some cases due to geographic proximity. Effects of environmental dissimilarity were marginally reduced when controlling for genetic similarity, but were still strongly negative (*r* = -0.35, *p* < 0.001), suggesting that similar environments promote genomic parallelism irrespective of genetic similarity. Phenotypic dissimilarity was no longer associated with genomic parallelism after controlling for genetic similarity (*r* = - 0.10, *p* = 0.097). This latter result suggests that environmental similarity is a better predictor of genomic parallelism than phenotypic similarity (at least in terms of observable morphometric phenotypes) in this system.

In conclusion, our study is the largest to date in this system addressing the relative effects of environment, phenotype and genetics in predicting parallel evolution. Genetic similarity is the best predictor of genomic parallelism here, in line with recent results and expectations regarding sharing of ancestral variants and introgression between populations (^29,39^). However, even whilst controlling for this, environmental variation (most likely Ca and pH variation in particular) is a good predictor of genome-wide parallelism among freshwater populations on a wide geographic scale. In particular, the higher environmental similarity among North American radiations, compared to European populations, provides a good explanation for the strong phenotypic and genomic parallelism among those freshwater populations, alongside the probable genetic bottleneck in the founding of Atlantic marine populations (^39^). The fact that, despite the larger geographic distance between the two North American radiations, they have more similar phenotypes and exhibit stronger genomic parallelism demonstrates that environment, and by extension selection, can counteract the effects of distance to some extent. Explicit measurements of environment are thus important in predicting parallel evolution even across large geographic scales, and distance alone can be misleading. Phenotypic parallelism was observed for several traits across continents, and populations with similar phenotypes also exhibit stronger genomic parallelism. However, this relationship is weaker than for commonality of environment, and is likely confounded by variable heritability and genetic architecture among phenotypes. Most importantly, our results highlight that quantitative analyses of phenotypes and environments and of their relationship can provide a good prediction of expected genomic parallelism, and provide a much clearer picture of major factors that are likely to influence the emergence of a common pattern of genomic divergence than analyses of dichotomous phenotypes and environments.

## Methods

### Sampling and environmental data collection

We sampled 18 lakes in North Uist, Scotland between April and June 2013, 18 lakes from Iceland between May and June 2014, 18 lakes from British Columbia (BC) between April and May 2015 and 19 lakes from the Cook Inlet basin, Alaska in June 2015. Lake names, geographic coordinates and numbers of samples used in the study are shown in Supplementary Table 1. We measured the pH, concentrations of metallic cation concentrations sodium (“Na”), calcium (“Ca”) and zinc (“Zn”) of each lake, and calculated population prevalence of *Gyrodactylus spp*. and *Schistochephalus solidus*. Concentrations of cations, pH and parasite prevalence per lake are shown in Supplementary Table 2. Details of the fish collection and quantification of abiotic and abiotic variables can be found in Supplementary Information.

### DNA extractions, RAD library preparation and sequencing

Genomic DNA was purified from 10 to 21 individuals from each of the populations, chosen to represent a widely distributed subset of the most environmentally and phenotypically variable lakes (Supplementary Dataset 3). Extracted genomic DNA was normalized to a concentration of 25 ng /μL in 96-well plates.

In 2014 we conducted RAD sequencing on samples from Scotland and from Iceland. Sequencing libraries were prepared and processed into RAD following the modified libraries according to (^65^). In 2016 we conducted RAD sequencing on samples from British Columbia and from Alaska. Sequencing libraries were prepared following the modified single-digest RAD protocol of (^66^). The two RADseq protocols interrogate the same set of loci across the genome, so that the SNP data are compatible across all four radiations. See Supplementary Information for details of the RAD library preparation and sequencing.

### Population genetics statistics and phylogenetic tree

Raw sequence reads were demultiplexed using Stacks – 1.35 (^67^). Number of reads per individual are shown in Supplementary Dataset 3 (see Supplementary Information for details on the alignment of reads and Stacks pipeline used). For Bayenv2 data, autosomal SNPs were called as being in >7 populations, >50% of the individuals within a population, and with a MAF-filter of 0.05. After filtering we retained 26,990, 26,937, 29,111, 26,169 SNPs for Scotland, Iceland, British Columbia and Alaska respectively. For analyses of population structure across all radiations, a subset of unlinked SNPs were generated. Here, autosomal SNPs were called that were present in all radiations and in >50% of individuals within a radiation, with a MAF-filter of 0.05 (within a radiation). Only the first SNP per RAD locus was retained. F_ST_ was bootstrapped and calculated in POPULATIONS. This set of SNPs were then pruned for linkage disequilibrium in plink using indep-pairwise 50 5 0.2. The set of unlinked SNPs were used to construct a neighbour-joining tree for all fish in the R package ‘adegenet’, using a distance matrix computed from the SNP data (^68^). The tree was bootstrapped 100 times and nodes with less than 80% support were collapsed. The tree was plotted using the ‘ape’ package in R (^69^). PCA analysis of population structure was conducted using plink (^70^).

### Phenotypic and environmental variation - body shape, armour, gill rakers and environmental data analyses

All morphological measurements (body shape, body armour and spine traits) were done following (^25^). Details of the quantification of phenotypic traits can be found in Supplementary Information.

We performed three Principal Component Analyses (PCAs): one on the armour traits, another on body shape, and another on the 6 environmental variables. Body shape and armour PCAs were performed on regression residuals of all individuals from all radiations pooled together to extract the common PCs of body shape and armour variation (Shape_PCs and ArmourPCs) and environmental variation and retained axes that explained more than 10% of the total variance. Armour and environmental PCAs were conducted with scaled inputs due to different units of measurement between variables. Shape PCAs were conducted on morphometric residuals, and as such were not scaled. All phenotypic analyses, including ANOVAs and ANCOVAs and plotting were done in R version 3.4.3.

### Genotype-Environment/Phenotype Associations

For each radiation separately (N=18 to 19 populations) we used Bayenv2 (^43^) to identify associations between genomic allele frequencies (N=10 to 21 individual fish, mean= 17. 8), the set of six biotic and abiotic environmental variables (Ca, Na, pH, Zn, prevalence of *Gyrodactylus spp*. and *Schistochephalus solidus*) and the set of 12 phenotypic traits (Shape_PC1, Shape_PC2, Shape_PC3, DS1, DS2, PS, LP, HP, BAP, Plate_N, Gill_Raker_L and Gill_Raker_N) mentioned above. For each radiation, a matrix of genetic covariance was calculated using a subset of SNPs limited to a single SNP per RAD-locus and pruned for linkage disequilibrium (R^2^ < 0.4) in plink (^70^). This cut-off was selected to balance the trade-off between SNPs retained and minimising the effects of linkage. Covariance matrices were therefore calculated using 9619, 7983, 7300 and 5705 SNPs for Alaska, BC, Iceland and Scotland respectively. Covariance matrices were calculated across 100,000 iterations and averaged across 5 independent runs. Bayenv2 was run independently 8 times and final results were averaged across runs.

Environmentally and phenotypically associated SNPs were selected as having a log_10_-BayesFactor > 1.5 and an absolute Spearman’s rank coefficient above the 95th percentile. The combination of BayesFactor and non-parametric measure of correlation helps to avoid selecting SNPs with high BayesFactors due to spurious populations (^18^). SNPs were grouped into 50kb, 75kb, 100kb, 200kb and 0.1 cM windows (Supplementary Table 14) to test the robustness of our results across different extents of linkage. To evaluate whether windows were environmentally or phenotypically associated, we adapted the methodology of (^46^). We calculated the upper 99% binomial expectation for the number of associated SNPs given the total number of SNPs in a specific window, and selected windows that had a greater number of associated SNPs than this expectation. This method controls for variation in SNP density across windows and ensures that significant windows exhibit consistent allele frequency correlations across multiple SNPs. We visualised the genomic locations of associated windows using Manhattan plots (Supplementary Fig. 5) and plotting the residual number of outlier SNPs above the binomial expectation (Supplementary Fig. 6). Linkage groups I-XXI were visualised with the exception of XIX; windows on scaffolds were not visualised. Finally, we compared these associated windows across radiations to examine those that were parallel.

### Parallelism statistics

For all radiation groupings (11 combinations in total: one four radiation grouping, four three radiation groupings, and six two radiation groupings), we calculated the significance of parallel window counts using a permutation method. For each environmental or phenotypic variable, we randomly drew N windows from each radiation’s total pool where N was equivalent to the associated window count for each radiation. We then assessed the overlap of randomly associated windows across radiations and pooled the results over 10,000 iterations. The output from all permutations was used as a null distribution to infer p-values, which were then FDR-corrected using the R package *qvalue* (^71^).

### Grouping of adjacent windows and expanding parallelism regions

Windows of 50kb and above were based on a linkage assumption and to minimise non-independence between windows. There were, however, occasionally adjacent windows associated with the same variable across different groupings. Large regions of relatively strong linkage are plausible if recombination is reduced through processes such as genomic rearrangements. To investigate these, we grouped associated windows that were adjacent as well as those that were direct matches across radiations based on the likelihood of adjacent associated windows resulting independently being low, suggesting non-independence and probable linkage. These windows are available in Supplementary Table 13.

### Multivariate vector comparison of environments and phenotypes

Vectors for environmental, shape and armour PCs (>10% variance) and gill raker data were calculated as the difference between the linearly predicted maximum and minimum values per radiation. Angles (θ) and difference in length (Δ*L*) were calculated for each vector between radiations. Significance of vectors was determined through permutations by simulating random traits from a normal distribution with mean and s.d. equivalent to the observed data and assessing vectors of random traits/variation as above (Supplementary Information).

### Comparing relative influences of environment, phenotype and genetics

F_ST_ was calculated between each freshwater population and its relevant marine population (Alaska = MUD1, BC = LICA, Iceland = NYPS, Scotland = OBSM) in 50kb windows using the R package ‘PopGenome’ (REF). For each MxF comparison, windows above the 95% quantile were classed as outliers. Outlier windows were compared across all pairwise freshwater comparisons (2,628 comparisons among 73 populations), with overlapping outliers representing MxF F_ST_ parallelism. Dissimilarity matrices of environment and phenotype were calculated as Euclidean distance in PCA space for the 6 and 12 environmental and phenotypic variables respectively. The genetic dissimilarity matrix was composed of genome-wide pairwise F_ST_ estimates between freshwater populations. The matrix of MxF parallelism was associated to environmental, phenotypic and genetic dissimilarity matrices using Mantel tests with 9,999 permutations. Partial Mantel tests were performed with genetic distance as the conditional matrix for environmental and phenotypic effects on MxF parallelism, again with 9,999 permutations.

## Supporting information

Supplementary Information Magalhaes Whiting et al

## Acknowledgements

We thank Shaun Robertson, Rebecca Young, Abdul Rahman, Brian Santos, Sara Goodacre, Petur Halldorsson, Bjarni K. Kristjánsson, Dolph Schluter, Kieran Samuk, Diana Rennison and Sara Miller for help with the sampling and sampling permits. We are grateful to Ann Lowe and Laura Dean for help with the DNA extractions, to Cody Wiench, Amanda Stahlke and Sarah Hendricks for help making the RAD-libraries and to John Brookfield for discussion of probability calculations. This work was funded by a NERC grant (NE/J02239X/1 to A.D.C.M), and further support was provided by NIH grant P30GM103324.

## Contributions

I.S.M, A.D.C.M and J.R.W. conceived the project, interpreted the data, and wrote the manuscript. I.S.M, D.D., and A.D.C.M performed field work. I.S.M, M.M. and D.D. generated the phenotypic data. I.S.M. and P.H. generated RAD data and J.R.W., I.S.M. and P.H. analysed it. P.H., M.B. and S.S. helped with the sampling and revised the manuscript.

## Data Accessibility

Bam files of aligned reads for each individual and corresponding sample information have been deposited in the European Nucleotide Archive database under the project PRJEB20851, with the sample accession numbers ERS1831811-ERS1833111, and run accession numbers ERR2055459-ERR2056759. Scripts used for all analyses are available at https://github.com/JimWhiting91/stickleback_adaptive_radiations/

