## Supplementary Information Magalhaes Whiting et al for "Intercontinental genomic parallelism in multiple adaptive radiations"

**This PDF file includes:**

Supporting Information and Supplementary Methods – pages 2 to 6

Supplementary Figs. 1 to 6 – pages 7 to 29

Supplementary Tables 1 to 14 – pages 30 to 58

Captions for Supplementary Datasets 1 to 4 – page 59

References for SI citations – pages 59-30

**Other supplementary materials for this manuscript include the following:**

Supplementary Datasets 1 to 3

**Supporting Information 1 - Marine x Freshwater (MxF) comparisons**

We wanted to compare our freshwater results to well-studied genomic parallelism in marine x freshwater comparisons. This analysis had two aims: 1) to act as a positive control to assess our methods ability to detect regions of known genomic parallelism; 2) to compare the extent of freshwater parallelism to MxF parallelism. To this end, we compiled four additional SNP datasets. Each dataset included the freshwater populations for a specific radiation and additional genomic data from four marine populations sampled from each country (OBSM - Scotland; NYPS - Iceland; LICA - BC, MUD - Alaska). Thus, each dataset contained 23 (Alaska) or 22 (BC, Iceland, Scotland) populations. SNPs were called using stacks Populations program, with a map assigned to each dataset that denoted “marine” and “freshwater” populations. SNPs that were not present in both “marine” and “freshwater” populations were removed; SNPs present in <50% of individuals within either “marine” or “freshwater” were removed; SNPs with a minor allele frequency below 0.05 were removed; all SNPs per locus were retained; and data from the sex chromosome (XIX) were also removed. SNPs were output to VCF format with 24,416, 26,658, 21,089 and 20,605 SNPs retained in Alaska, BC, Iceland, and Scotland respectively.

To make the MxF analyses as comparable as possible to our intra-radiation analyses, we used Bayenv2 in the same way. To generate Bayesfactors and Spearman’s ρ, we calculated allele frequencies within each of the 23 or 22 populations and assigned marine populations “1” and freshwater populations “-1” in the environment input file. Covariance matrices were calculated using a subset of each SNP dataset pruned for linkage and averaged over 10 independent runs. Bayenv2 was run over each SNP for 100,000 MCMC iterations with results averaged over 10 independent runs. Outliers were determined using the methods used for intra-radiation analyses, and all downstream analyses were run in the same way. To recap, outlier SNPs were determined as having log_10_(Bayesfactors) > 1.5 and Spearman’s ρ values above the 0.95 quantile. Outlier windows were determined as those that contained more outlier SNPs than a 99% binomial expectation given the number of SNPs in that window. Outlier windows were overlapped, and significance was determined by comparing observed overlap to a null distribution of random overlap over 10,000 permutations. In total, we detected 91 (Alaska), 34 (BC), 30 (Iceland) and 29 (Scotland) 50kb outlier windows, which represented 1.07%, 0.40%, 0.35% and 0.34% of windows that contained SNPs.

For 50kb windows, one window (groupXX:8950000-9000000; FDR = 0.0003) was found to overlap in all four MxF datasets, and three overlapped in Alaska, Iceland and Scotland (groupI:21600000-21650000; groupI:21750000-21800000; groupIV:12800000-12850000; FDR < 0.0001). In pairwise comparisons, overlap was greatest for Alaska and BC (N_windows_ = 8; FDR < 0.0001), followed by Iceland and Scotland (N_windows_ = 5; FDR < 0.0001), Alaska and Iceland (N_windows_ = 2; FDR = 0.08), Alaska and Scotland (N_windows_ = 1; FDR = 0.409), BC and Iceland (N_windows_ = 1; FDR = 0.204), and no windows overlapped in BC and Scotland.

**Supporting Information 2 - Windows based on genetic distance (0.1 cM) and recombination estimates**

Variable linkage across the genome moderates our ability to detect genomic associations with environmental/phenotypic variation by influencing linkage between sequenced and causative variants. Thus, our ability to detect associations is lower in a low-linkage window than a high-linkage window despite equal physical size of the windows. To assess whether this may influence our results, we estimated windows based on genetic distance, using a previously published genetic map (1). This approach is less-desirable than our physical distance windows because the genetic map itself has limited coverage within chromosomes and does not extent to unplaced scaffolds. However, it does allow us to assess how patterns of genomic parallelism may be driven by variable linkage in a subset of our original data. We chose to use windows of 0.1 cM as this gave us a broadly similar number of windows to our 50kb dataset.

Signals of parallelism for 0.1 cM windows were strongest for variables for which we similarly detected genomic parallelism with 50kb windows: calcium, pH, pelvic spine length, pelvis length, plate N and gill raker N all exhibited parallelism FDR < 0.001. Further, weakly significant genomic parallelism (FDR < 0.05) was recovered for *Gyrodactylus* prevalence, salinity, zinc, dorsal spine 1 & 2 length and pelvis height. This suggests that our physical distance windows may be conservative estimates of parallelism.

0.1 cM windows also recovered several instances of higher order parallelism (parallelism in > 2 radiations) than observed for 50kb windows. Two windows on chromosome 4 (46.3-46.4 cM; 42.6-42.7 cM) were found associated with zinc variation and plate N respectively in Alaska, BC and Iceland. A further window on chromosome 4 (44.2-44.3 cM) was associated with pelvis length in BC, Iceland and Scotland. This latter window likely corresponds to our three-way pelvis length window detected at 50kb resolution.

To estimate recombination for our 50kb windows, we overlapped recombination rate estimates between genetic markers in the map with our 50kb windows and took the weighted mean of overlapping recombination regions. Recombination was weighted according to the proportional coverage of each recombination interval within the 50kb window.

**Supplementary methods:**

### Sampling and environmental data collection.

Fish were collected by setting between 10 and 30 unbaited minnow traps (Gee traps, Dynamic Aqua, Vancouver, Canada) in water approximately 0.3–3m deep, within 5 m of shore along a 100–400m stretch of shoreline. The fish were haphazardly selected with individuals of all sizes, sex and breeding condition collected (see Supplementary Table S17 for details of sex of each fish collected).

We chose environmental variables that were likely to cause natural selection on the fish and could be precisely measured . For abiotic environmental variables we chose pH, calcium (Ca), sodium (Na) and zinc (Zn), which have been associated previously with the evolution of body shape, size and armour in stickleback (2, 3, 4). We measured the pH of each lake using a calibrated pH meter (Multi 340i, WTW, Weilheim, Germany). The concentrations of metallic cation concentrations sodium (“Na”), calcium (“Ca”) and zinc (“Zn”) were obtained by collecting a filtered water sample acidified with 2% nitric acid in the field from each lake. These samples were then analysed at the Division of Agriculture & Environmental Science at the University of Nottingham for metallic cation concentrations by inductively coupled plasma mass spectrometry (ICP-MS) and anions using a Dionex DX500 ion chromatograph with an IonPac AS14A (4 x 250 mm).

Many biotic variables are difficult to quantify precisely so we used the prevalence of two parasites *Gyrodactylus* sp. (ectoparasitic trematodes) and *Schistocephalus solidus* (endoparasitic cestode) that are likely to have a significant impact on the reproduction and life cycle of stickleback (5, 6, 7). The prevalence of *Gyrodactylus spp.* and *Schistochephalus solidus* per fish were counted during dissections and parasite numbers were averaged by lake. Concentrations of cations, pH and parasite prevalence per lake are shown in Supplementary Table S2.

**Phenotypic data collection.**

The 12 phenotypic traits are all known to have undergone major changes as result of the migration of sticklebacks from marine to freshwater habitats (8) and are known to be highly variable between freshwater populations (4).

All morphological measurements and analyses were done using the left side of the fish. To quantify body shape variation across individuals and lakes we digitized 13 homologous landmarks (n=16) using the TPS software package (8, 9, 10). Landmark coordinates for 1298 individuals were exported, and analyzed using MORPHOJ 1.03 (12). Briefly, we first performed a Procrustes superimposition to extract shape coordinates for further analyses (13, 14). For each radiation separately, we then performed a size correction to account for any allometric effects (15, 16) using multivariate regressions of Procrustes coordinates against the logarithm of centroid size and tested its significance using a permutation test against the null hypothesis of independence (10 000 iterations).

Using photographs of the left side of alizarin-stained individual fish we measured the following body armour and spines traits: number of armour plates (“Plate_N”), length of 1^st^ dorsal spine (“DS1”) and 2^nd^ dorsal spine (“DS2”), length of biggest armour plate (“BAP”), pelvic spine length (“PS”), length of the horizontal process of the pelvis (“LP”) and height of the ascending process of the pelvis (“HP”) (30). The first gill arch from at least three fish per population (mean = 13.8) was removed and photographed at 10x magnification under a graticule. The total number of gill rakers (“Gill_Raker_N”) was then counted from the photograph, and the three longest rakers measured to give an average ‘gill raker length’ (“Gill_Raker_L”). As all traits, apart from number of plates, were significantly correlated with SL (results not shown), we size corrected the data by regressing each trait against SL (all individuals from all lakes pooled) and used the residuals in further analyses.

**DNA extractions, RAD library preparation and sequencing.**

In 2014 we conducted RAD sequencing on samples from Scotland and from Iceland. Sequencing libraries were prepared and processed into RAD following the modified libraries according to (17), using the restriction enzyme SbfI-HF (NEB), which digests DNA at an 8bp recognition sequence and is expected to cut roughly 22,000 locations across the stickleback genome. Each sample was individually ligated to adaptors with 6bp in-line barcodes and multiplexed in libraries of 192. We sequenced these RAD libraries in four lanes on an Illumina HiSeq sequencer at the University of Oregon, producing 100-bp single-end reads.

In 2016 we conducted RAD sequencing on samples from British Columbia and from Alaska. Sequencing libraries were prepared following the modified single-digest RAD protocol of (16), using the restriction enzyme SbfI. Each sample was individually labelled using one of 96 unique 8bp barcodes, and each pool of 96 samples received a different 8bp inline index using the NEBNext Ultra DNA Library Prep Kit for Illumina (18). We sequenced these RAD libraries in two lanes of Illumina NextSeq high-output sequencing at the University of Oregon, using paired-end 75bp reads. After filtering for the presence of a correct index, barcode, and SbfI cut site, these two lanes of sequencing produced a total of 256 million and 267 million read pairs, respectively.

**Population genetics statistics.**

The retained reads were aligned to the three-spined stickleback reference genome (version BROADs1, Ensembl release 82) using GSnap (19). Reference mapping with GSnap took sequence quality information into account, allowed for up to five mismatches and up to 2 indels between each read and the reference sequence and ignored reads that mapped against more than a single position in the genome. The STACKS pipeline was used to analyse mapping files and population genetics statistics were calculated using the POPULATIONS program in Stacks. POPULATIONS was run independently for each radiation with the following filters applied: SNPs that were present in less than 8 populations were removed; SNPs present in <50% of individuals within a population were removed; SNPs with a minor allele frequency below 0.05 were removed; all SNPs within a locus were retained; and data from the sex chromosome (XIX) were also removed. These filters were chosen to maximise SNP count whilst still providing enough information for allele frequencies and environmental and phenotypic variables to be correlated. After filtering we retained 26,990, 26,937, 29,111, 26,169 SNPs within a GENEPOP formatted output for Scotland, Iceland, British Columbia and Alaska, respectively. GENEPOP files were converted to BAYENV2 format using PGDSpider2 (20).

**Permutations for genomic parallelism**

We quantified genomic parallelism associated with individual variables by carrying out randomised permutations (akin to χ^2^, but avoiding Poisson assumptions) with the expectation that all of the windows to which our (environmentally and phenotypically associated) SNPs mapped had an equal probability of being associated with environmental or phenotypic variables, assuming independence between windows. Whilst the latter is technically untrue, non-independence should be minimal at window sizes of 50Kb or greater given linkage disequilibrium in the stickleback genome. Results were also consistent at larger window sizes (up to 100 kb) where non-independence is further minimised but traded-off against signal.

**Multivariate vectors**

To quantify variation within a vector, we calculated the major vector through each radiation within multivariate space. Put simply, rather than a vector representing the divergence between two populations in contrasting habitats, here vectors represent the major axis of variation across all populations in a radiation. As such, the direction of the vector is unimportant, and orthogonality (90°) is the most divergent angle possible. We produced four separate vectors representing environmental variation, shape variation, armour variation and gill raker morphologies. For each of these groupings, we used PCA to remove collinearity and retained PCs that explained at least 10% of the total variance. We then scaled PC scores according to their variance explained (under the assumption that equal weighting should not be given to deviations along PC4 if, for example, it only represents a fraction of the absolute variance relative to PC1). For each scaled, independent PC trait, we calculated the degree of change as the difference between the minimum and maximum predicted values of the linear relationship across all population-means within a radiation. We then calculated the angle between vectors (θ) using the R function *angle.calc()* and the difference in vector lengths (Δ*L*). Vectors that share low (θ) suggest radiations exhibit similar directions of variation, and thus convergent trait covariance, whilst low values Δ*L* indicate similar magnitudes of variation. Because PC scalings were dependent on the group of variables being examined, values for Δ*L* are relative and only comparable within variable groupings.

To understand whether values of θ were lower than expected through random chance, we simulated new variable distributions based on normal distributions with mean and standard deviation matching observed data. For example, to simulate a randomised armour vector, we calculated mean and standard deviation within each radiation for each of the 7 armour variables considered, and used this information to simulate 7 new ‘random’ armour variables. We then repeated the process detailed above, removing collinearity through PCA and performing vector calculations. We permuted this analysis 1,000 times for environmental, shape, armour and gill-raker vectors, and compared null distributions of θ against observed values with a one-tailed hypothesis.

**Supplementary Fig. 1**. Three-dimensional plot of the PCA on 8,395 SNPs for 1,304 individuals.

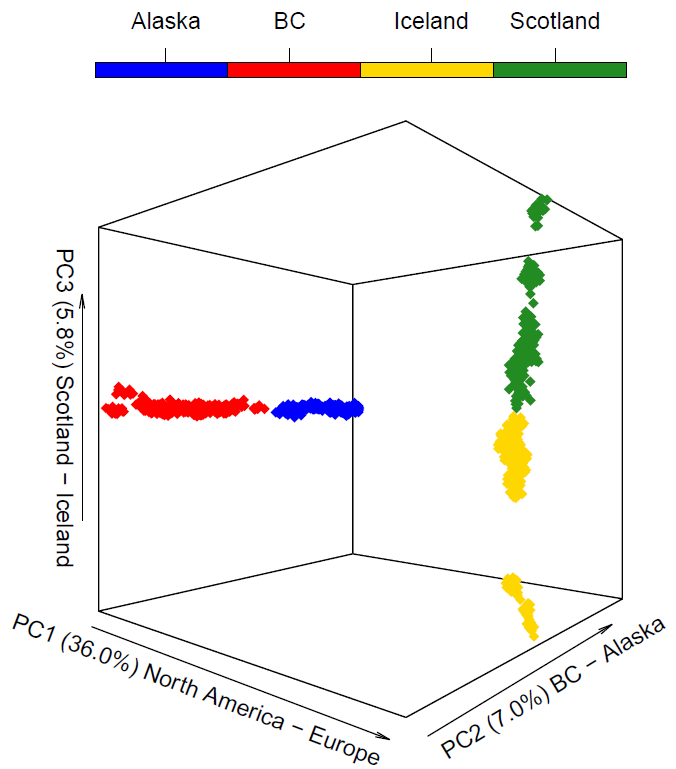

Supplementary Fig. 2. Numbers of 50kb windows with SNPs associated with each of the 6 environmental variables (upper Venn diagram) and 3 classes of phenotypic traits (lower Venn diagram) that are shared and unique among the four adaptive radiations. Numbers presented here are counts of windows associated with each individual variable or group of variables, so if a window is associated with more than one variable it will count towards numbers of parallel windows presented in more than one segment. This is purely for visualisation purposes, as only unique window numbers were used to estimate levels of parallelism between radiations. Phenotypic traits are pooled into the following categories: armour traits (dorsal spines 1 and 2, pelvic spine, height and length of pelvis, length of biggest armour plate and number of armour plates); Body shape PCs (PC1, 2 and 3 of body shape) and gill rakers (gill rakers number and length). For phenotypic traits *N* is the number of traits pooled per category.

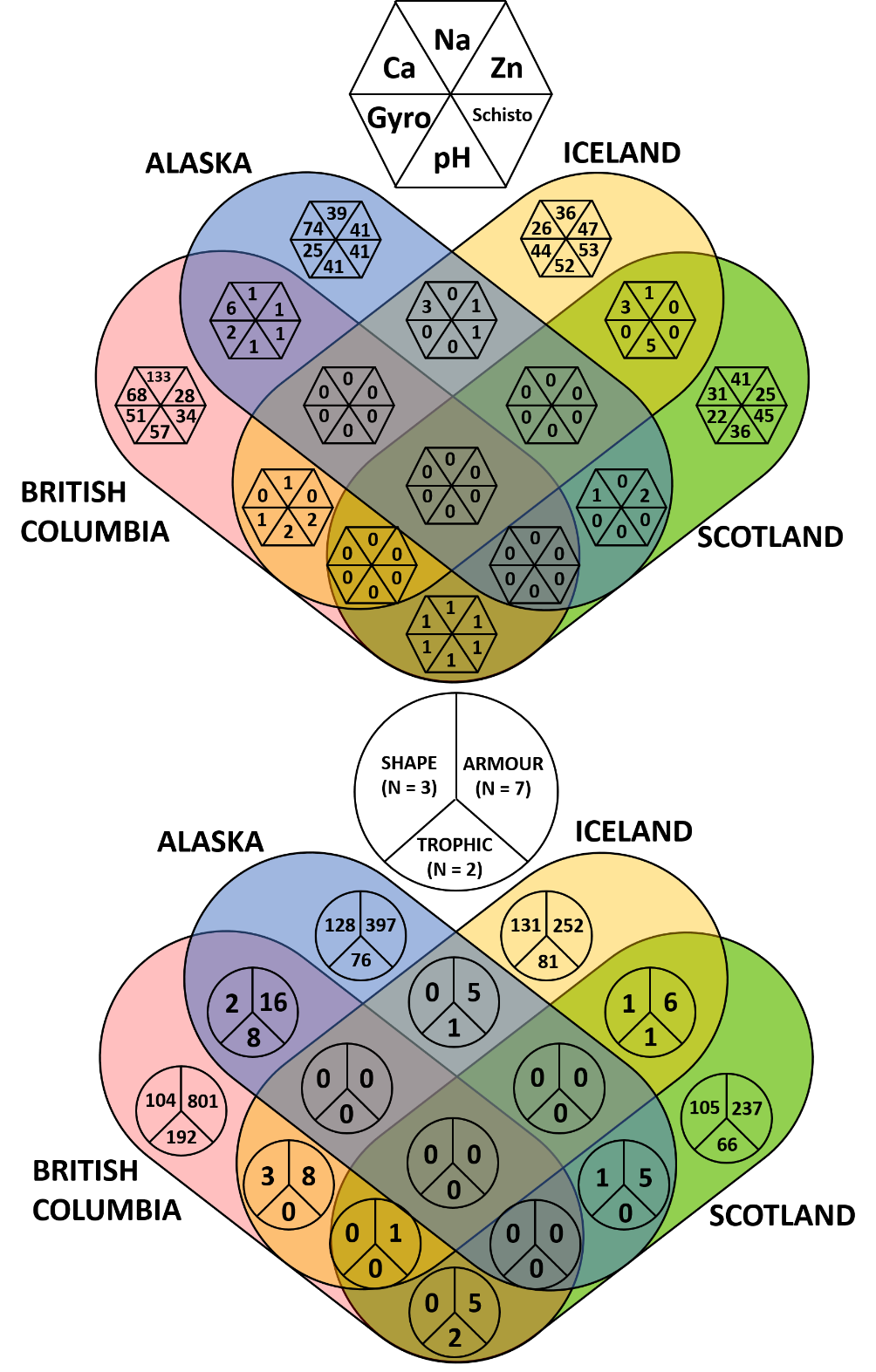

Supplementary Fig. 3. Recombination for all windows, classified as non-associated (‘Neutral’), environmentally or phenotypically associated (‘Outlier’), or parallel across radiations (‘Overlap’). Pairwise wilcoxon tests are shown between all violins (p > 0.05 = NS; p < 0.001 = ***).

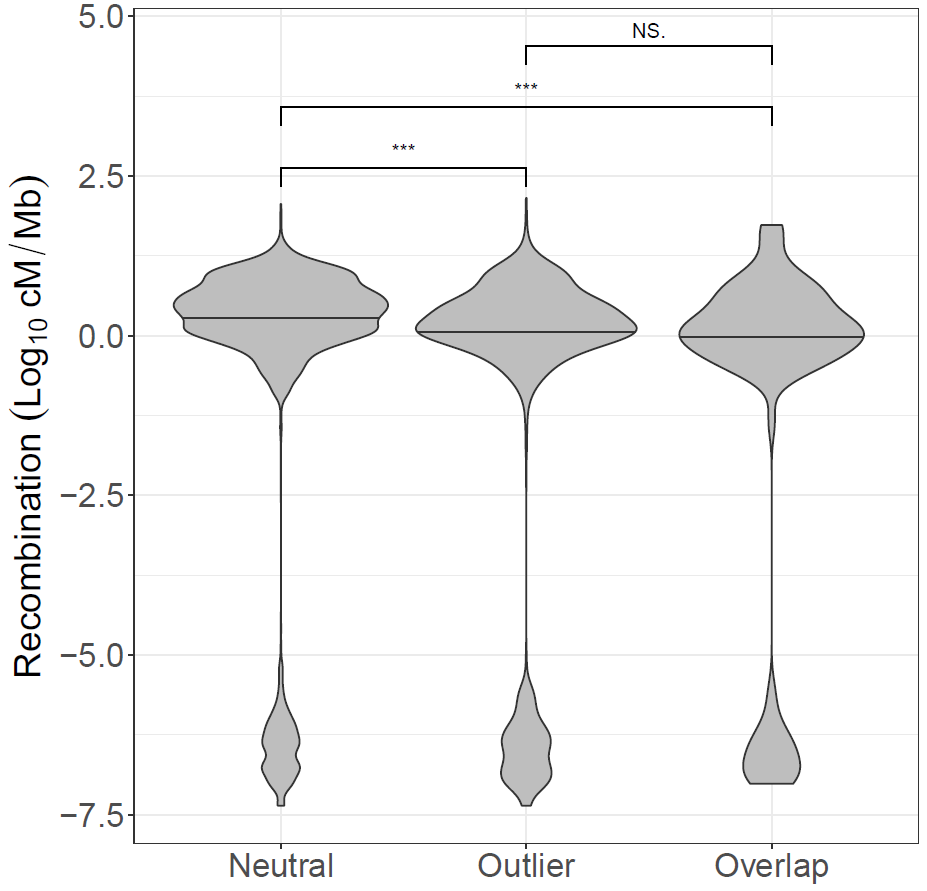

**Supplementary Fig. 4**. Expected and observed counts of 0.1cM windows containing an above 99% binomial expectation number of SNPs associated with environmental variables and phenotypic traits in at least 2 radiations. Expected bars represent mean counts across 10,000 simulated outcomes with 95% confidence intervals per a one-tailed hypothesis. Asterisks denote significance of FDR-corrected one-tailed tests between the observed counts and the 100,000 simulated counts at the <0.05 (*), <0.01 (**) and <0.001 (***) levels.

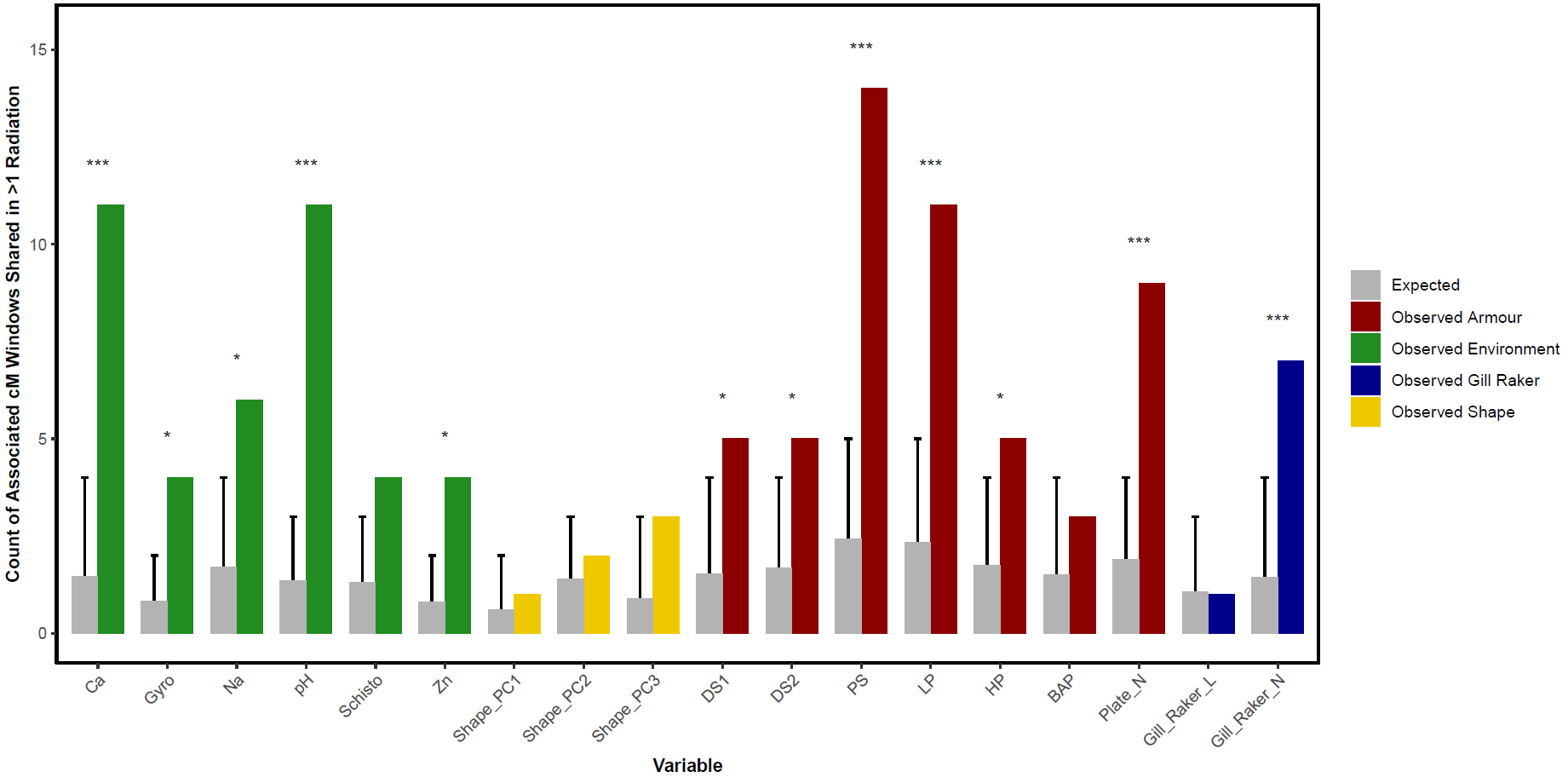

**Supplementary Fig. 5**. Manhattan plots summarizing correlations of allele frequency with all environmental variables and phenotypic traits. Bars show residual associated SNPs above the 99% binomial expectation. Parallel windows are highlighted as circles (associated in 2 radiations) or triangles (associated in 3 radiations).

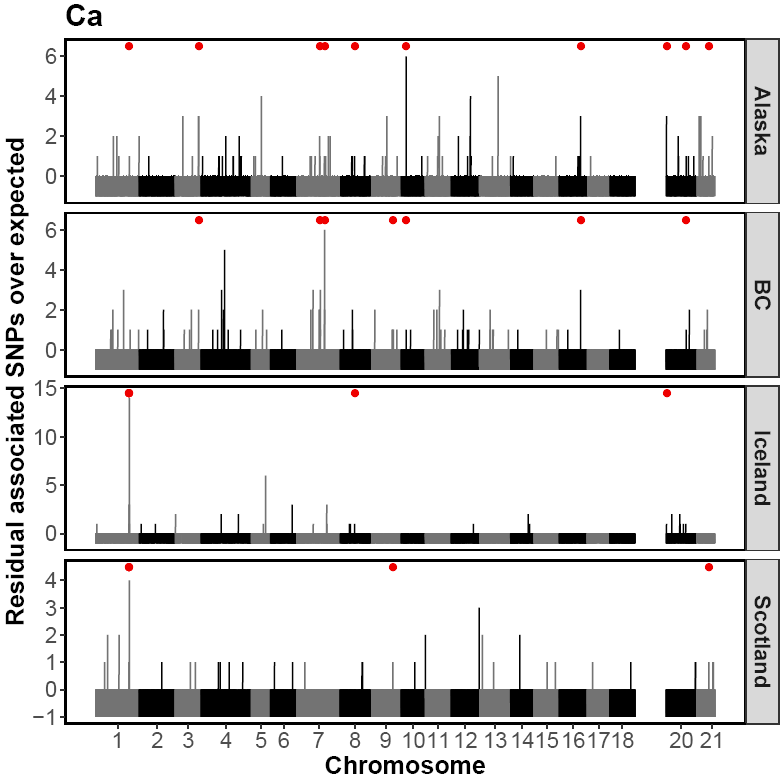

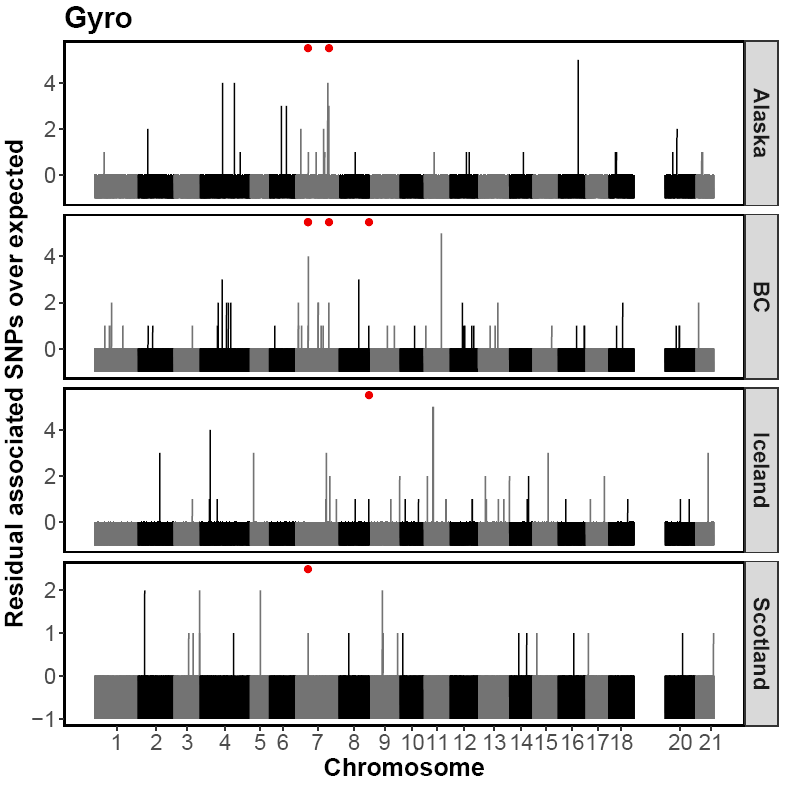

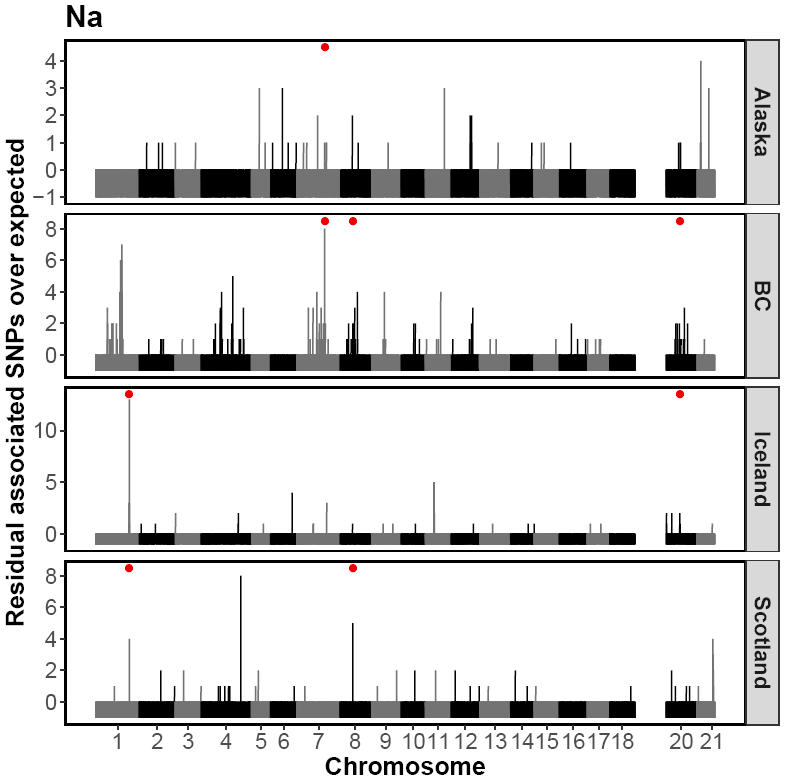

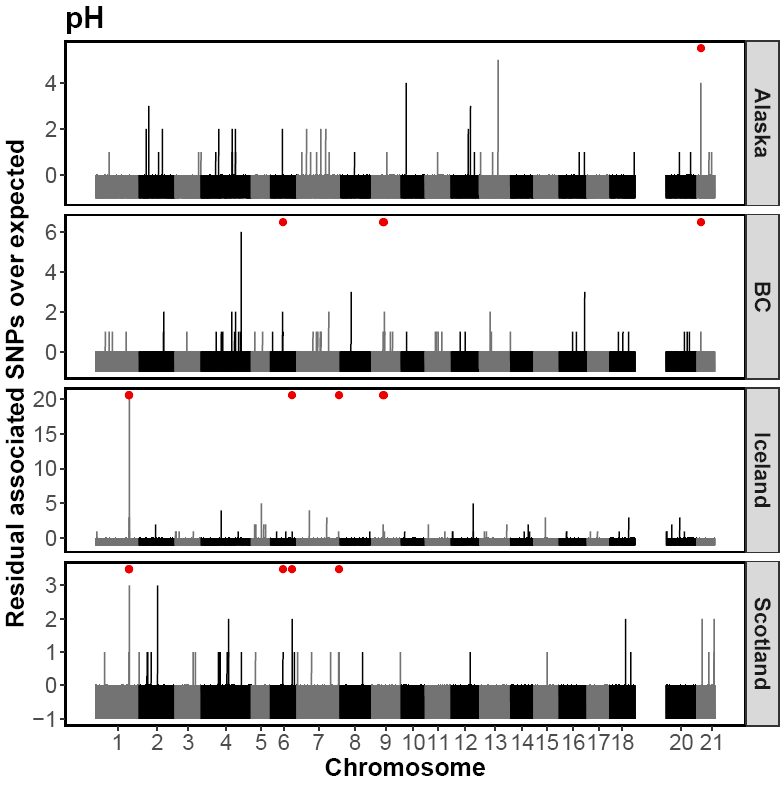

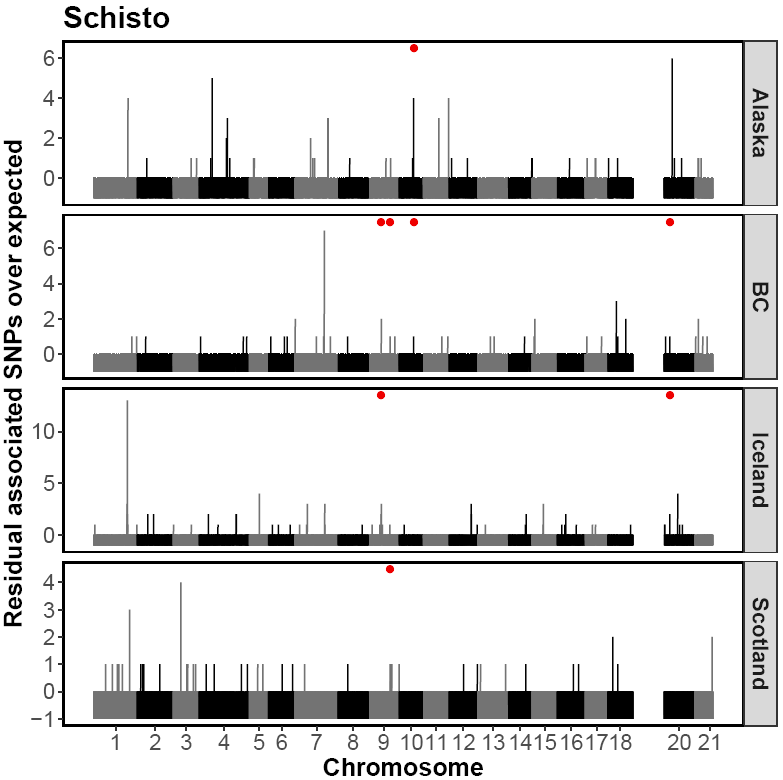

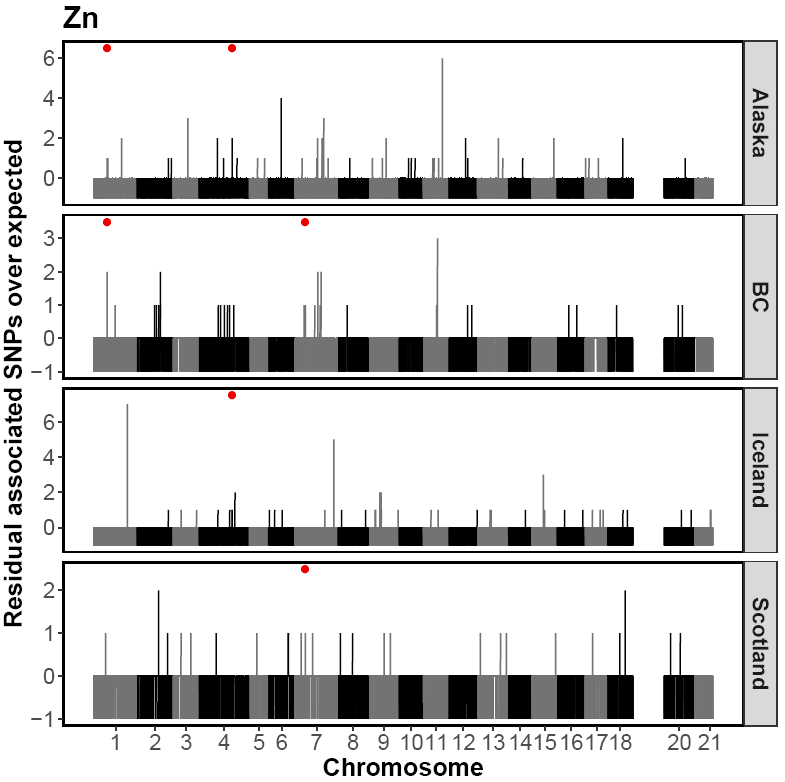

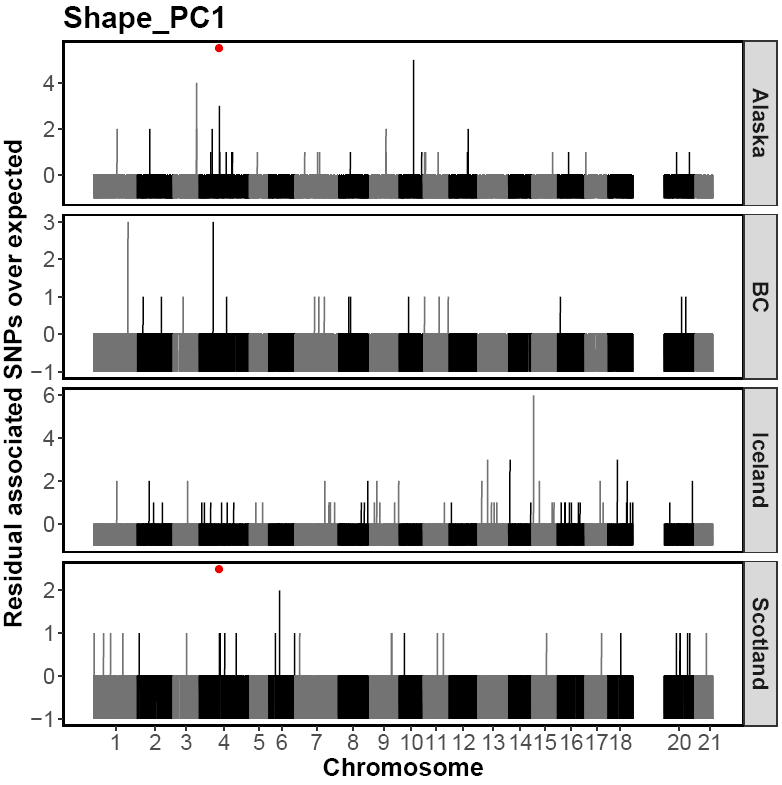

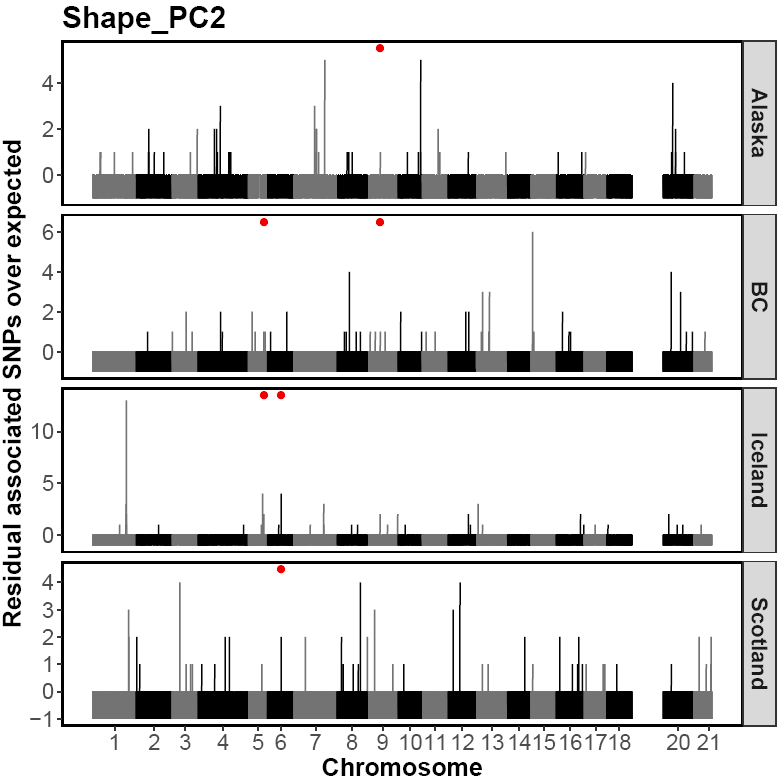

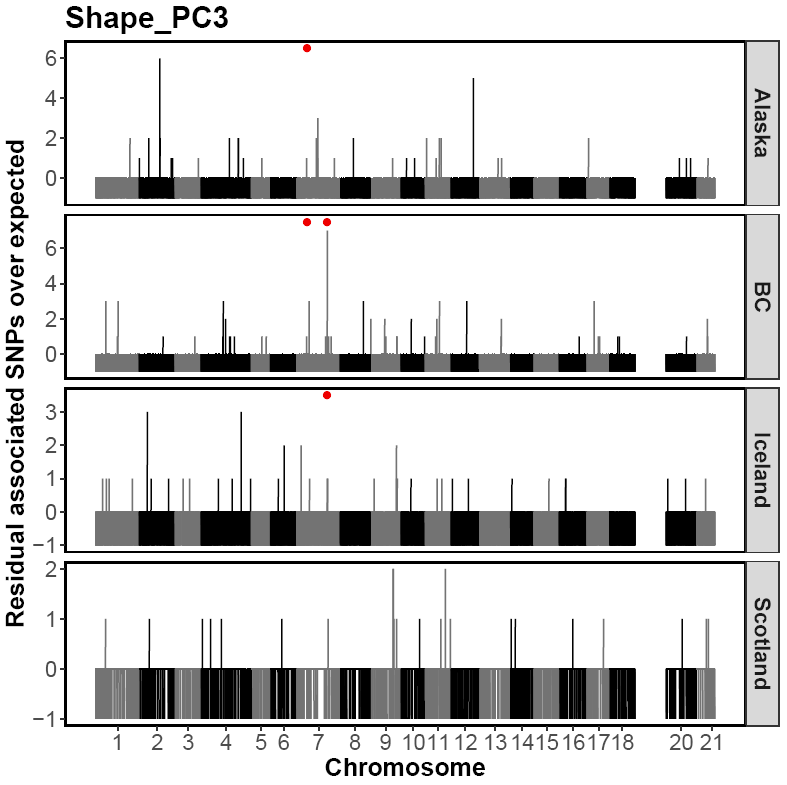

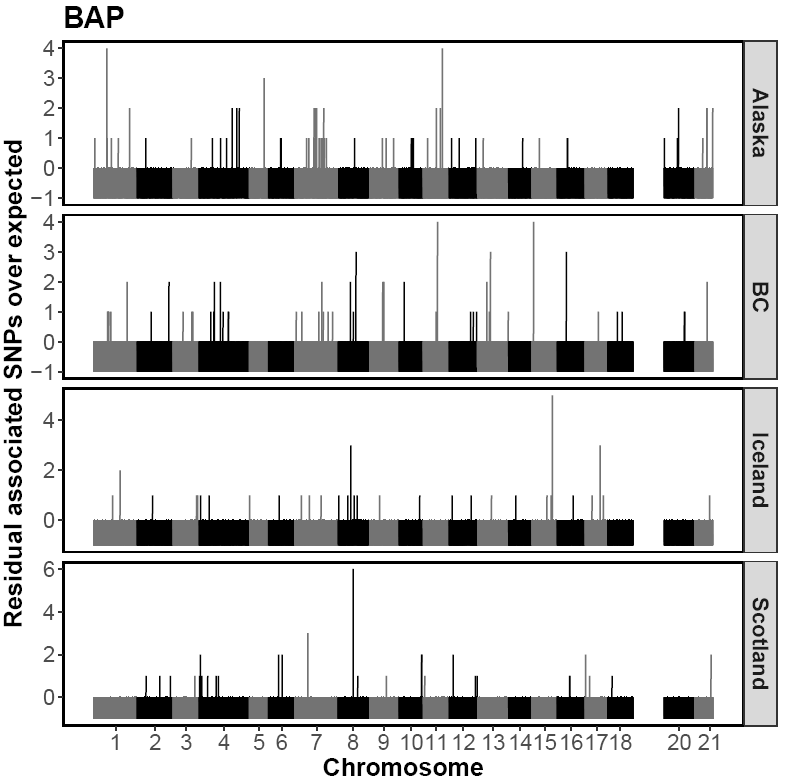

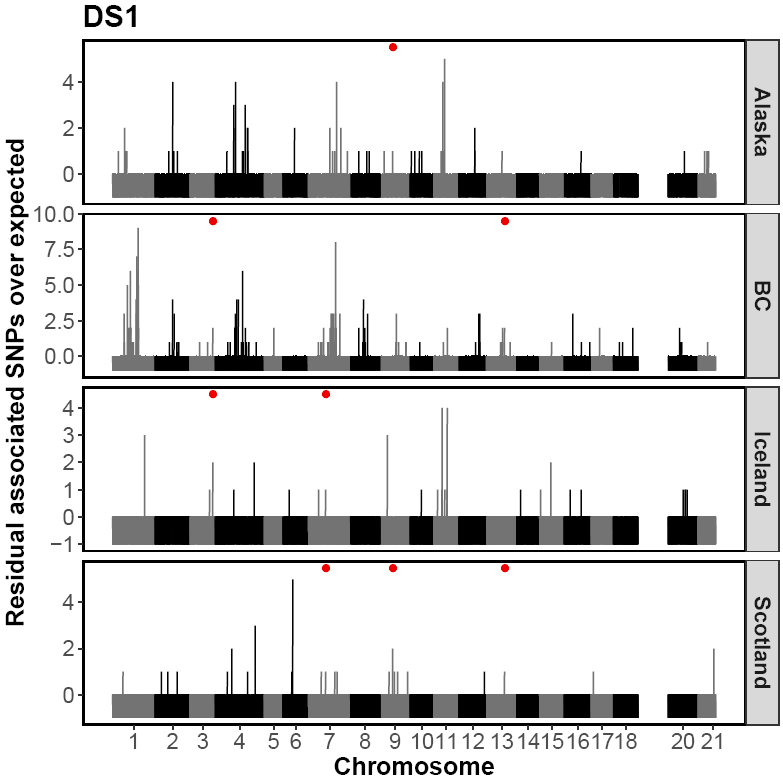

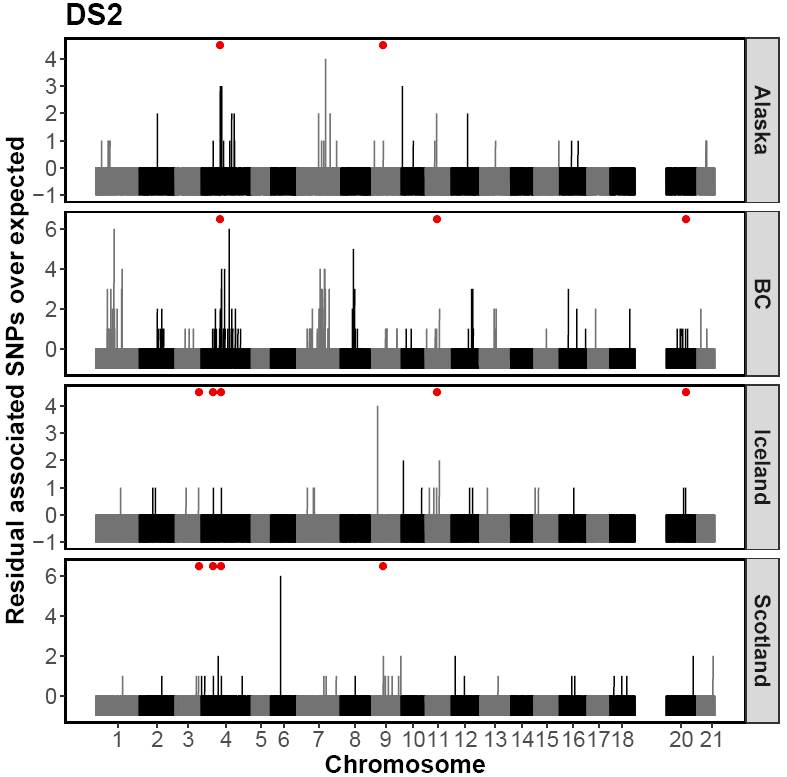

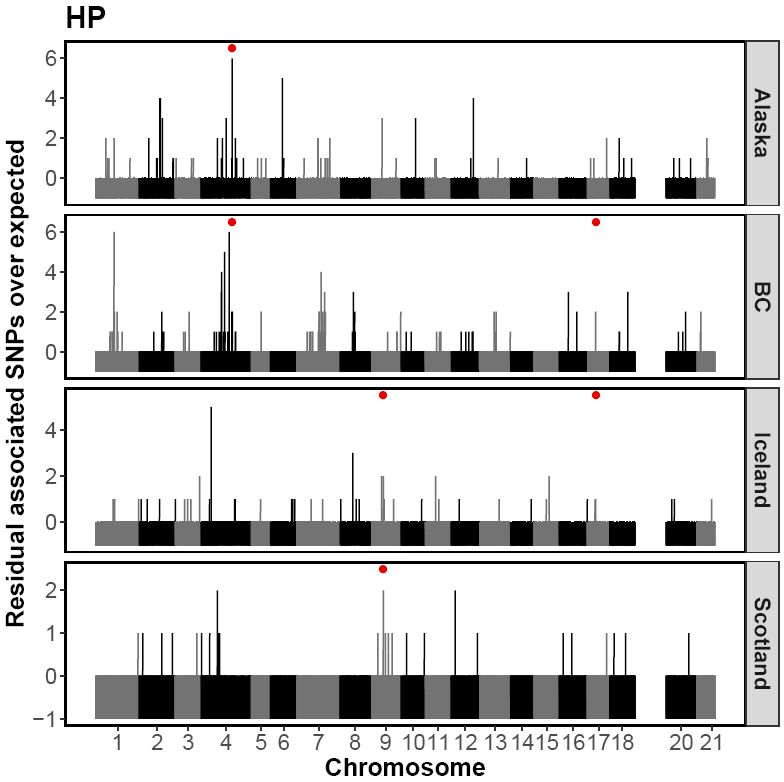

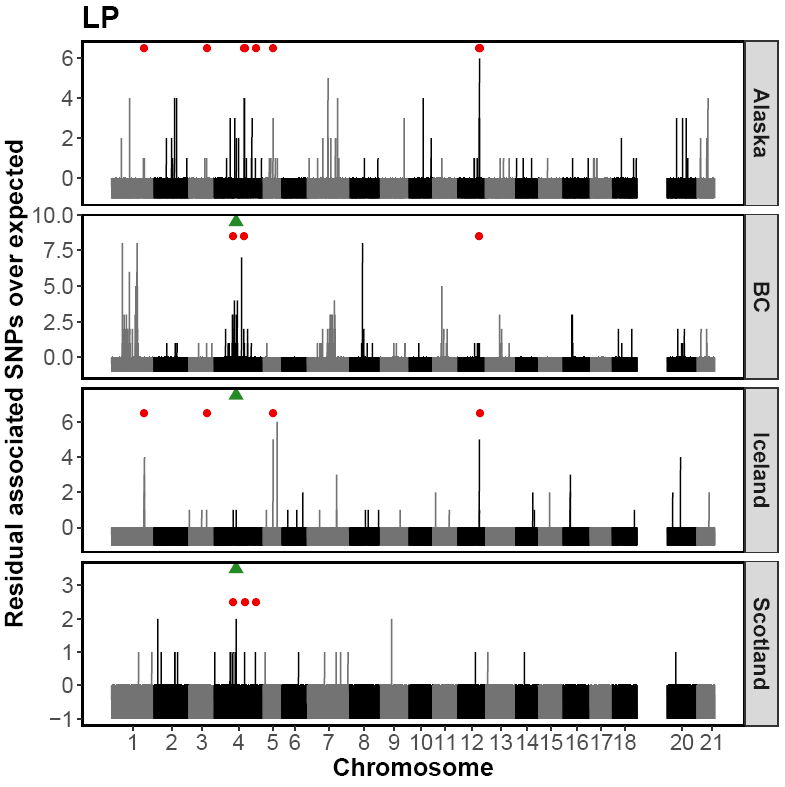

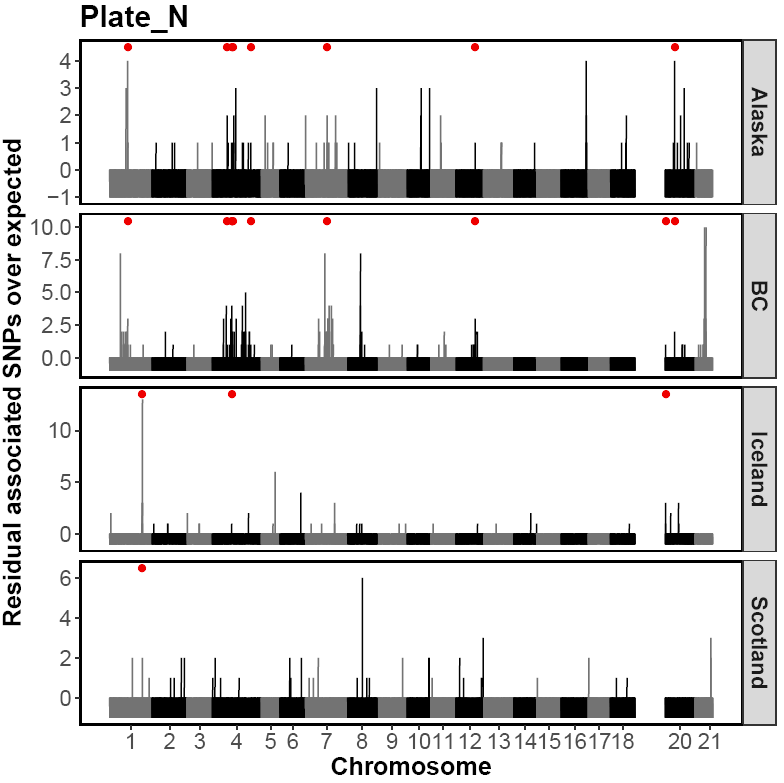

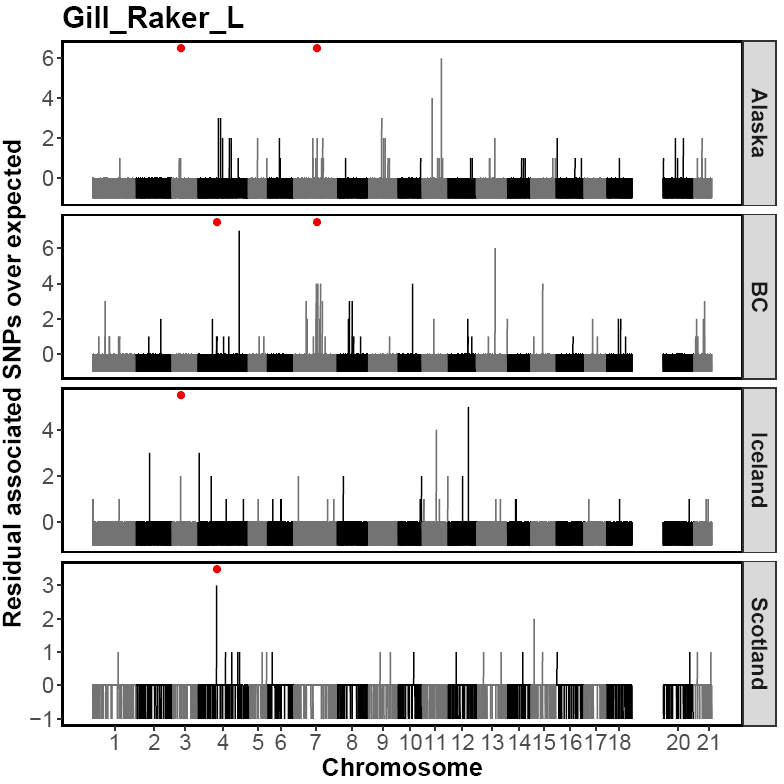

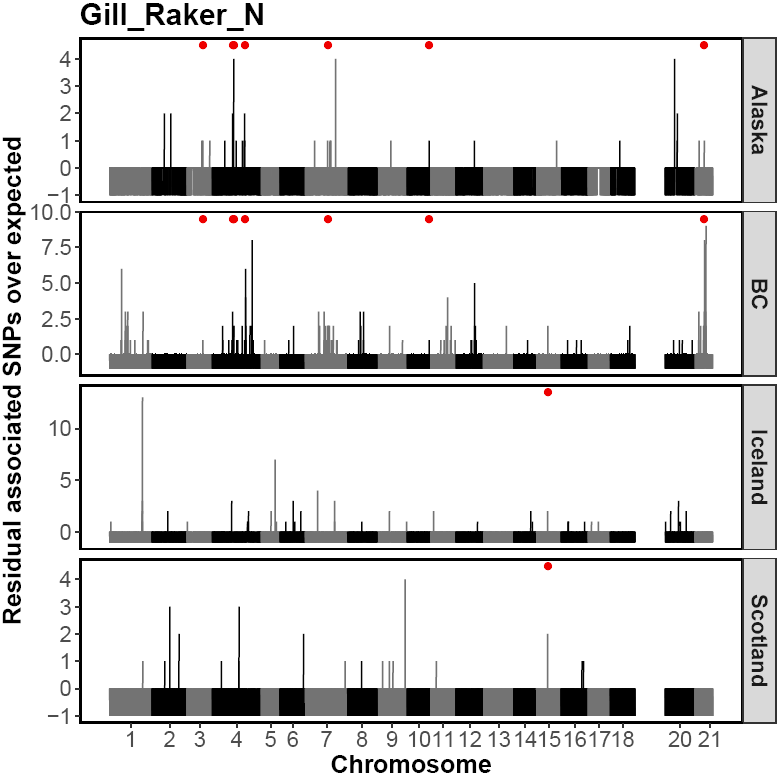

**Supplementary Fig. 6**. Example illustration for Calcium of outlier windows on the basis of SNPs above a 99% binomial expectation (blue line). Red points denote 50kb outlier windows, whilst black points denote 50kb windows with associated SNPs in line with or less than the binomial expectation, given the number of SNPs within each window (SNP count).

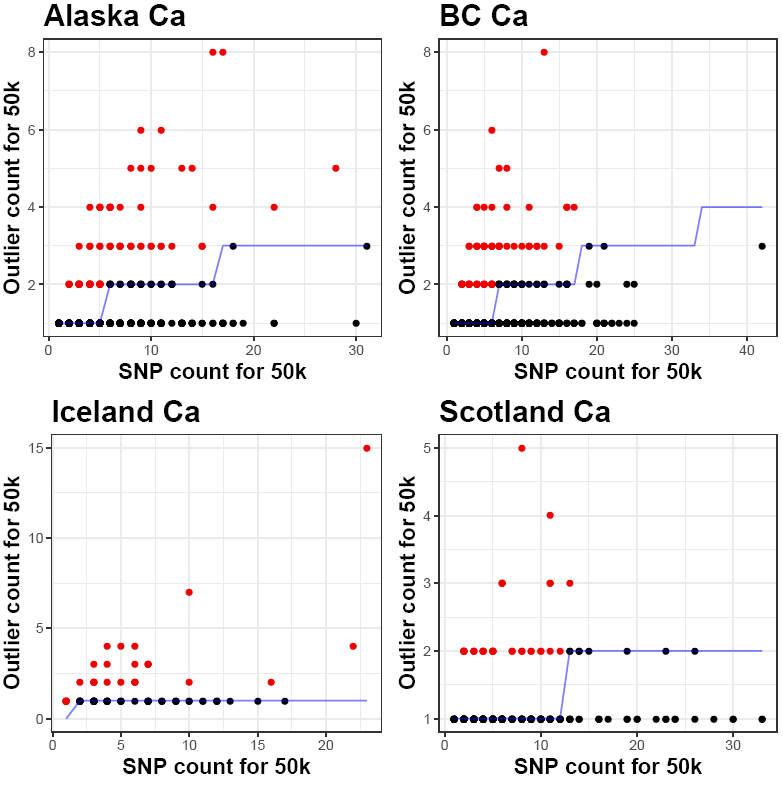

Supplementary Table 1. Names of lakes sampled, abbreviations of their names, geographic region and area within the geographic region where samples came from, date of sampling, geographic coordinates of each lake and numbers of three-spined sticklebacks analysed from each lake.

| Population name | Code | Geographic region | Area | Date | Latitude | Longitude | N |
| --- | --- | --- | --- | --- | --- | --- | --- |
| *Freshwater* |  |  |  |  |  |  |  |
| Aonghais | AONG | Scotland | W | 16.05.2013 | 57°38'39.89"N | 7°16'27.49"W | 18 |
| Mhic a'Roin | AROI | Scotland | S | 09.05.2013 | 57°35'40.8"N | 7°25'52.5"W | 19 |
| a'Bharpa | BHAR | Scotland | S | 14.05.2013 | 57°34'15.1"N | 7°18'07.1"W | 18 |
| na Buaile | BUAI | Scotland | NE | 13.05.2013 | 57°38'48.94"N | 7°11'52.85"W | 17 |
| Chadha Ruaidh | CHRU | Scotland | SE | 29.04.2013 | 57°35'38.63"N | 7°11'51.84"W | 17 |
| an Daimh | DAIM | Scotland | S | 30.04.2013 | 57°35'35.40"N | 7°12'33.27"W | 20 |
| Eisiadar | EISI | Scotland | N | 17.05.2013 | 57°37'54.97"N | 7°21'14.53"W | 17 |
| Fada | FADA | Scotland | E | 17.05.2013 | 57°37'3.69"N | 7°12'33.92"W | 15 |
| nan Geireann | GEIR | Scotland | N | 16.05.2013 | 57°38'24.56"N | 7°17'24.93"W | 17 |
| Mhic Gille-bhride | GILL | Scotland | W | 09.05.2013 | 57°36'7.11"N | 7°24'35.29"W | 17 |
| Hosta | HOST | Scotland | NW | 15.05.2013 | 57°37'37.65"N | 7°29'27.28"W | 19 |
| Iala | IALA | Scotland | E | 16.05.2013 | 57°37'11.28"N | 7°12'20.41"W | 18 |
| na Moracha | MORA | Scotland | S | 07.05.2013 | 57°34'24.75"N | 7°16'33.73"W | 21 |
| na Reival | REIV | Scotland | W | 04.05.2013 | 57°36'40.78"N | 7°30'52.84"W | 19 |
| Scadavay | SCAD | Scotland | S | 06.05.2013 | 57°35'4.09"N | 7°14'9.56"W | 17 |
| nan Strùban | STRU | Scotland | SW | 17.05.2013 | 57°33'29.8"N | 7°21'09.1"W | 16 |
| Tormasad | TORM | Scotland | SW | 01.05.2013 | 57°33'43.7"N | 7°19'00.6"W | 19 |
| Trosavat | TROS | Scotland | SW | 19.05.2013 | 57°35'3.85"N | 7°24'48.40"W | 20 |
| Bakkatjorn | BAKK | Iceland | SW | 02.06.2014 | 64° 9'19.29"N | 22° 1'6.99"W | 19 |
| Eidarvatn | EIDA | Iceland | E | 17.06.2014 | 65°23'17.26"N | 14°21'35.94"W | 19 |
| Flodid | FLOD | Iceland | N | 09.06.2014 | 65°29'27.95"N | 20°21'33.59"W | 20 |
| Galtabol | GALT | Iceland | N | 09.06.2014 | 65°15'40.89"N | 19°44'24.30"W | 19 |
| Grettislaug | GRET | Iceland | N | 13.06.2014 | 65°52'57.3"N | 19°44'13.1"W | 19 |
| Grjotarvatn | GRJO | Iceland | W | 08.07.2014 | 64°46'36.39"N | 22° 0'6.89"W | 18 |
| Holsvatn | HOLS | Iceland | W | 26.05.2014 | 64°30'48.00"N | 22° 8'43.28"W | 18 |
| Hredavatn | HRED | Iceland | W | 26.05.2014 | 64°45'56.66"N | 21°34'21.72"W | 18 |
| Kleifarvatn | KLEI | Iceland | SW | 22.05.2014 | 63°55'27.55"N | 21°59'56.42"W | 16 |
| Mjoavatn | MJOA | Iceland | N | 09.06.2014 | 65°15'41.45"N | 19°48'4.60"W | 20 |
| Myvatn-lava | MYVL | Iceland | NE | 14.06.2014 | 65°37'41.49"N | 16°55'31.57"W | 19 |
| Myvatn-mud | MYVM | Iceland | NE | 14.06.2014 | 65°39'7.67"N | 16°58'8.95"W | 20 |
| Sauravatn | SAUR | Iceland | W | 26.05.2014 | 64°39'47.23"N | 22° 7'32.61"W | 16 |
| Skorradalsvatn | SKOR | Iceland | W | 30.06.2014 | 64°28'57.48"N | 21°19'2.22"W | 19 |
| Thingvallavatn | THIN | Iceland | SW | 23.05.2014 | 64°11'11.6"N | 21°05'18.1"W | 20 |
| Urridavatn | URR2 | Iceland | SE | 17.06.2014 | 65°17'59.45"N | 14°27'21.82"W | 19 |
| Urridakotsvatn | URRI | Iceland | E | 19.05.2014 | 64° 4'5.41"N | 21°54'34.08"W | 19 |
| Vífilsstaðavatn | VIFI | Iceland | SW | 18.05.2014 | 64° 4'45.72"N | 21°52'38.86"W | 20 |
| Ambrose | AMBR | B.C. | Sechelt P. | 20.05.2015 | 49°44'4.70"N | 124° 1'32.52"W | 17 |
| Beaver | BEAV | B.C. | Vancouver I. | 12.05.2015 | 48°48'41.96"N | 124° 4'51.38"W | 18 |
| Brannen | BRAN | B.C. | Vancouver I. | 13.05.2015 | 49°12'51.16"N | 124° 2'59.50"W | 18 |
| Bullocks | BULL | B.C. | Saltspring I. | 10.05.2015 | 48°52'24.4"N | 123°30'22.0"W | 15 |
| Cranby | CRAN | B.C. | Texada I. | 29.04.2015 | 49°41'31.85"N | 124°30'27.00"W | 20 |
| Dougan | DOUG | B.C. | Vancouver I. | 12.05.2015 | 48°42'51.73"N | 123°36'38.44"W | 16 |
| Errock | ERRO | B.C. | Fraser Valley | 22.05.2015 | 49°13'28.73"N | 122° 0'41.42"W | 18 |
| Garden Bay | GARD | B.C. | Sechelt P. | 26.04.2015 | 49°39'3.62"N | 124° 0'59.67"W | 10 |
| Hoggan | HOGG | B.C. | Gabriola I. | 14.05.2015 | 49° 9'12.09"N | 123°49'41.94"W | 18 |
| Hotel | HOTE | B.C. | Sechelt P. | 25.04.2015 | 49°38'19.46"N | 124° 3'9.01"W | 18 |
| Kennedy | KENN | B.C. | Vancouver I. | 15.05.2015 | 49° 7'35.72"N | 125°25'35.84"W | 18 |
| Kirk | KIRK | B.C. | Texada I. | 01.05.2015 | 49°44'24.21"N | 124°34'59.82"W | 19 |
| Klein | KLEN | B.C. | Sechelt P. | 26.04.2015 | 49°44'20.34"N | 123°58'0.93"W | 18 |
| Lily | LILY | B.C. | Sechelt P. | 25.04.2015 | 49°36'44.02"N | 124° 1'16.73"W | 19 |
| North | NORT | B.C. | Sechelt P. | 26.04.2015 | 49°44'52.60"N | 123°58'31.91"W | 14 |
| Sproat | SPRO | B.C. | Vancouver I. | 15.05.2015 | 49°17'04.3"N | 124°58'25.5"W | 18 |
| Stowell | STOW | B.C. | Saltspring I. | 10.05.2015 | 48°46'55.12"N | 123°26'41.46"W | 17 |
| Trout | TROUT | B.C. | Sechelt P. | 25.04.2015 | 49°30'25.02"N | 123°52'27.28"W | 18 |
| Arness | ARNE | Alaska | Kenai | 15.06.2015 | 60°38'45.73"N | 151°18'7.36"W | 17 |
| Arrow | ARRO | Alaska | Kenai | 24.06.2015 | 60°45'1.29"N | 150°29'21.07"W | 15 |
| Barley | BARL | Alaska | Mat-Su | 08.06.2015 | 61°21'40.09"N | 150° 5'1.20"W | 18 |
| Bear Paw | BEPA | Alaska | Mat-Su | 03.06.2015 | 61°36'50.38"N | 149°45'11.91"W | 17 |
| Big | BIGL | Alaska | Mat-Su | 08.06.2015 | 61°31'58.86"N | 149°50'3.79"W | 15 |
| Bruce | BRUC | Alaska | Mat-Su | 12.06.2015 | 61°36'31.97"N | 149°33'4.77"W | 16 |
| Community | COMM | Alaska | Kenai | 15.06.2015 | 60°42'8.72"N | 151°23'1.32"W | 17 |
| Corcoran | CORC | Alaska | Mat-Su | 05.06.2015 | 61°34'23.07"N | 149°41'30.40"W | 16 |
| Daniel | DANI | Alaska | Kenai | 15.06.2015 | 60°43'28.79"N | 151°10'49.99"W | 18 |
| Duck | DUCK | Alaska | Kenai | 15.06.2015 | 60°41'7.33"N | 151°13'25.41"W | 16 |
| Jade | JADE | Alaska | Mat-Su | 08.06.2015 | 61°31'29.25"N | 149°52'1.50"W | 18 |
| Long | LONG | Alaska | Mat-Su | 11.06.2015 | 61°34'33.76"N | 149°46'29.47"W | 18 |
| Luci | LUCI | Alaska | Mat-Su | 03.06.2015 | 61°34'11.93"N | 149°28'53.26"W | 18 |
| Lynda | LYND | Alaska | Mat-Su | 08.06.2015 | 61°34'13.44"N | 149°50'24.75"W | 17 |
| Seymour | SEYM | Alaska | Mat-Su | 03.06.2015 | 61°36'52.7"N | 149°39'51.3"W | 17 |
| Tern | TERN | Alaska | Kenai | 24.06.2015 | 60°32'2.70"N | 149°32'49.59"W | 18 |
| Toad | TOAD | Alaska | Mat-Su | 03.06.2015 | 61°37'12.12"N | 149°41'56.11"W | 19 |
| Walby | WALB | Alaska | Mat-Su | 22.06.2015 | 61°37'13.2"N | 149°12'48.5"W | 18 |
| Y | YLAK | Alaska | Mat-Su | 22.06.2015 | 62°18'21.42"N | 150° 3'52.34"W | 19 |
| *Marine* |  |  |  |  |  |  |  |
| Ob nan Stearnain | OBSM | Scotland | S | 17.05.2013 | 57°36'4.82"N | 7°10'24.50"W | 19 |
| Nypslon | NYPS | Iceland | NE | 17.06.2014 | 65°46'18.85"N | 14°50'6.10"W | 19 |
| Little Campbell River | LICA | B.C. | Surrey | 27.04.2015 | 49° 0'50.89"N | 122°45'38.50"W | 20 |
| Mud | MUD | Alaska | Mat-Su | 01.06.2015 | 61°35'50.39"N | 149°20'36.69"W | 18 |

**Supplementary Table 2.** Measured abiotic and biotic characteristics of 73 freshwater lakes and average phenotypic traits for populations from those lakes. From left to right: Calcium (Ca), sodium (Na) and zinc (Zn) concentrations, pH, prevalence of *Gyrodactylus* spp. (Gyro) and *Schistochephalus solidus* (Schisto), Pricipal Components 1, 2 and 3 of body shape (Shape PC1,2 ,3), length of dorsal spines 1 and 2 (DS1, DS2), length of pelvic spine (PS), lenght of pelvis (LP), height of pelvis (HP), length of biggest armour plate (BAP), number of armour plates (plate.N), standard length (SL), gill raker number (Gill_Raker_N) and length (Gill_Raker_L).

| Radiation | Population | Ca  (mg/L) | Na  (mg/L) | Zn  (µg/L) | pH | Gyro | Schisto | Shape  PC1 | Shape  PC2 | Shape  PC3 | DS1 | DS2 | PS | LP | HP | BAP | Plate  N | SL | GRN | GRL |
| --- | --- | --- | --- | --- | --- | --- | --- | --- | --- | --- | --- | --- | --- | --- | --- | --- | --- | --- | --- | --- |
| Alaska | ARNE | 8.106 | 4.192 | 48.26 | 7.41 | 0.926 | 0 | -0.012 | 0.002 | -0.015 | -0.065 | -0.066 | -0.060 | 0.016 | 0.043 | 0.049 | 6.059 | 58.529 | 18.059 | -0.015 |
| Alaska | ARRO | 0.736 | 1.8 | 31.582 | 6.92 | 0.9 | 0 | -0.011 | 0.016 | -0.003 | 0.012 | 0.026 | -0.492 | -0.133 | -0.502 | 0.004 | 4.800 | 57.333 | 20.200 | -0.068 |
| Alaska | BARL | 42.206 | 2.75 | 8.339 | 8.37 | 0.967 | 0.1 | -0.014 | 0.006 | -0.011 | -0.013 | -0.032 | -0.073 | 0.001 | 0.011 | -0.024 | 4.833 | 60.111 | 21.444 | -0.005 |
| Alaska | BEPA | 0.21 | 0.585 | 9.244 | 5.7 | 0.167 | 0.167 | -0.010 | 0.004 | 0.003 | -0.019 | -0.018 | -0.433 | -0.084 | -0.424 | -0.038 | 3.824 | 42.000 | 20.118 | -0.005 |
| Alaska | BIGL | 21.962 | 3.552 | 5.083 | 8.07 | 0.9 | 0.067 | 0.007 | -0.002 | -0.009 | 0.073 | 0.070 | 0.098 | 0.019 | -0.010 | -0.016 | 6.333 | 47.000 | 20.067 | 0.129 |
| Alaska | BRUC | 1.111 | 0.661 | 6.779 | 6.61 | 0.862 | 0 | -0.011 | -0.002 | -0.012 | -0.021 | -0.046 | -0.428 | -0.012 | -0.125 | 0.018 | 4.563 | 47.563 | 20.000 | 0.048 |
| Alaska | COMM | 0.325 | 1.836 | 26.339 | 6.8 | 0.769 | 0.067 | -0.006 | 0.012 | -0.015 | -0.043 | -0.032 | -0.279 | -0.023 | -0.191 | 0.011 | 6.389 | 44.556 | 19.833 | -0.019 |
| Alaska | CORC | 29.2 | 4.643 | 5.147 | 8.18 | 0.7 | 0.233 | -0.018 | 0.007 | -0.018 | 0.020 | 0.036 | 0.019 | 0.020 | 0.036 | 0.012 | 5.824 | 44.000 | 20.824 | -0.191 |
| Alaska | DANI | 13.515 | 4.53 | 14.593 | 7.51 | 0.533 | 0.333 | 0.009 | -0.004 | -0.016 | 0.042 | 0.045 | 0.041 | 0.038 | 0.010 | 0.045 | 6.667 | 48.889 | 20.278 | 0.135 |
| Alaska | DUCK | 12.677 | 4.842 | 13.619 | 9.68 | 1 | 0 | 0.001 | 0.001 | -0.019 | -0.015 | -0.020 | -0.046 | -0.004 | 0.054 | -0.016 | 7.000 | 50.438 | 18.563 | -0.060 |
| Alaska | JADE | 0.826 | 0.649 | 15.431 | 6.57 | 0.423 | 0.167 | -0.009 | 0.004 | -0.006 | -0.023 | -0.012 | -0.019 | -0.004 | 0.022 | -0.094 | 5.333 | 58.833 | 21.056 | 0.001 |
| Alaska | LONG | 12.837 | 2.426 | 4.968 | 7.8 | 0.833 | 0.033 | -0.002 | -0.002 | -0.013 | 0.074 | 0.076 | 0.079 | 0.020 | 0.044 | -0.027 | 6.278 | 52.611 | 21.444 | 0.112 |
| Alaska | LUCI | 29.942 | 11.982 | 7.562 | 8.17 | 0.885 | 0.167 | -0.021 | 0.005 | -0.009 | 0.033 | 0.031 | 0.026 | 0.023 | 0.073 | 0.054 | 6.000 | 47.222 | 20.722 | -0.044 |
| Alaska | LYND | 11.641 | 2.007 | 7.692 | 7.42 | 0.467 | 0 | -0.003 | -0.005 | -0.019 | -0.018 | -0.037 | 0.003 | 0.022 | -0.003 | -0.020 | 6.056 | 50.222 | 21.500 | 0.137 |
| Alaska | SEYM | 25.906 | 3.526 | 11.674 | 8.28 | 0.857 | 0.067 | -0.010 | 0.011 | -0.014 | -0.054 | -0.014 | -0.068 | -0.002 | 0.038 | 0.009 | 5.706 | 50.765 | 22.118 | 0.002 |
| Alaska | TERN | 32.131 | 3.036 | 7.936 | 7.6 | 0.767 | 0.3 | -0.024 | -0.009 | 0.004 | 0.035 | 0.033 | 0.030 | 0.033 | 0.070 | 0.024 | 6.895 | 43.368 | 18.632 | -0.129 |
| Alaska | TOAD | 0.604 | 0.688 | 6.639 | 5.99 | 0.567 | 0.267 | -0.011 | 0.007 | -0.008 | -0.069 | -0.045 | -0.496 | -0.093 | -0.483 | -0.011 | 4.350 | 58.500 | 20.850 | -0.051 |
| Alaska | WALB | 25.468 | 7.307 | 10.209 | 8.53 | 0.462 | 0.367 | -0.019 | 0.005 | -0.012 | 0.049 | 0.054 | 0.038 | 0.027 | 0.046 | 0.016 | 6.684 | 40.895 | 20.105 | -0.080 |
| Alaska | YLAK | 2.822 | 1.295 | 16.374 | 6.98 | 0.833 | 0.133 | -0.014 | 0.012 | -0.001 | 0.004 | -0.022 | -0.482 | -0.189 | -0.453 | 0.028 | 5.789 | 54.842 | 20.842 | 0.106 |
| BC | AMBR | 2.231 | 1.906 | 10.188 | 6.88 | 0.448 | 0 | -0.007 | 0.030 | 0.011 | 0.025 | -0.001 | -0.012 | -0.026 | -0.025 | -0.010 | 6.294 | 50.235 | 24.941 | 0.288 |
| BC | BEAV | 3.87 | 1.211 | 31.199 | 6.82 | 1 | 0.04 | -0.001 | 0.005 | 0.001 | 0.098 | 0.096 | 0.164 | 0.030 | 0.073 | 0.007 | 6.950 | 51.600 | 19.278 | -0.142 |
| BC | BRAN | 7.434 | 3.162 | 8.894 | 7.82 | 0.96 | 0 | 0.006 | 0.006 | -0.003 | 0.187 | 0.182 | 0.220 | 0.028 | 0.159 | 0.035 | 5.737 | 53.737 | 17.789 | -0.096 |
| BC | BULL | 10.456 | 8.551 | 7.802 | 7.58 | 0.739 | 0.059 | -0.007 | 0.015 | -0.021 | -0.200 | -0.133 | -0.217 | -0.065 | -0.093 | -0.066 | 3.867 | 48.200 | 19.667 | 0.093 |
| BC | CRAN | 15.627 | 3.86 | 13.049 | 7.23 | 1 | 0 | -0.004 | 0.010 | 0.000 | -0.104 | -0.108 | -0.096 | -0.029 | -0.057 | -0.023 | 5.450 | 45.100 | 19.850 | -0.009 |
| BC | DOUG | 20.007 | 6.98 | 15.88 | 7.52 | 0.917 | 0 | -0.021 | -0.006 | -0.006 | -0.146 | -0.148 | -0.165 | -0.018 | -0.079 | 0.031 | 5.063 | 51.750 | 20.188 | 0.033 |
| BC | ERRO | 3.03 | 1.997 | 45.606 | 7 | 0.833 | 0 | -0.008 | -0.005 | 0.009 | 0.127 | 0.106 | 0.183 | 0.016 | 0.132 | 0.057 | 32.500 | 55.444 | 19.889 | 0.198 |
| BC | GARD | 4.699 | 4.768 | 12.68 | 6.91 | 0 | 0 | 0.017 | 0.012 | 0.011 | 0.056 | 0.002 | 0.035 | 0.003 | 0.048 | 0.009 | 7.222 | 49.778 | 22.667 | 0.060 |
| BC | HOGG | 6.771 | 6.542 | 10.6 | 6.09 | 0.88 | 0 | -0.014 | 0.011 | -0.003 | -0.085 | -0.079 | -0.138 | -0.009 | -0.083 | -0.008 | 4.800 | 45.200 | 19.500 | 0.005 |
| BC | HOTE | 2.861 | 4.992 | 9.132 | 7.36 | 0 | 0 | -0.016 | 0.005 | 0.013 | 0.142 | 0.106 | 0.197 | 0.046 | 0.108 | 0.003 | 7.000 | 46.647 | 21.875 | 0.124 |
| BC | KENN | 5.891 | 1.079 | 11.865 | 7 | 1 | 0 | -0.010 | 0.008 | -0.003 | 0.156 | 0.131 | 0.239 | 0.036 | 0.105 | -0.025 | 30.895 | 54.263 | 22.316 | 0.165 |
| BC | KIRK | 19.732 | 2.959 | 8.884 | 7.6 | 0.778 | 0 | 0.002 | 0.019 | 0.000 | -0.112 | -0.102 | -0.195 | -0.004 | -0.168 | -0.041 | 4.000 | 54.333 | 20.667 | -0.271 |
| BC | KLEN | 5.101 | 1.491 | 7.852 | 6.65 | 0 | 0 | 0.001 | 0.016 | -0.012 | -0.287 | -0.100 | -0.305 | -0.027 | -0.150 | 0.012 | 5.529 | 46.000 | 21.133 | -0.175 |
| BC | LILY | 5.665 | 6.498 | 11.854 | 6.97 | 0.029 | 0 | 0.000 | 0.010 | 0.003 | 0.055 | 0.038 | 0.087 | 0.022 | 0.047 | 0.037 | 6.053 | 42.526 | 20.842 | 0.010 |
| BC | NORT | 4.362 | 2.191 | 3.418 | 7.11 | 0 | 0 | 0.005 | 0.020 | 0.005 | 0.104 | 0.077 | 0.090 | 0.021 | 0.005 | 0.017 | 33.643 | 47.357 | 22.286 | 0.014 |
| BC | SPRO | 8.646 | 1.149 | 15.463 | 7.04 | 0.88 | 0 | 0.006 | 0.012 | -0.004 | 0.071 | 0.080 | 0.126 | 0.009 | 0.098 | -0.019 | 7.947 | 61.316 | 20.105 | -0.059 |
| BC | STOW | 8.987 | 5.941 | 8.91 | 7.61 | 0.478 | 0 | -0.017 | -0.003 | -0.002 | -0.144 | -0.158 | -0.242 | -0.032 | -0.063 | 0.005 | 4.765 | 46.176 | 21.875 | -0.001 |
| BC | TROUT | 6.263 | 3.675 | 8.692 | 7.12 | 0.64 | 0 | 0.011 | 0.020 | -0.007 | 0.032 | 0.004 | -0.008 | -0.039 | -0.036 | -0.015 | 4.389 | 46.056 | 20.882 | -0.220 |
| Iceland | BAKK | 34.329 | 58.667 | 204.131 | 9.17 | 0.886 | 0 | -0.008 | 0.000 | 0.017 | -0.016 | -0.024 | 0.041 | 0.020 | 0.123 | 0.050 | 28.421 | 47.384 | 18.158 | 0.235 |
| Iceland | EIDA | 5.128 | 5.038 | 229.685 | 7.73 | 0.714 | 0.286 | 0.001 | -0.001 | -0.011 | -0.023 | -0.019 | -0.009 | -0.008 | -0.024 | -0.011 | 4.632 | 37.026 | 17.444 | -0.026 |
| Iceland | FLOD | 5.577 | 6.074 | 194.573 | 8.05 | 0.771 | 0.686 | -0.005 | -0.015 | 0.005 | 0.044 | 0.041 | 0.075 | 0.003 | 0.046 | 0.028 | 5.684 | 48.621 | 18.278 | -0.062 |
| Iceland | GALT | 6.022 | 5.013 | 203.798 | 7.7 | 0.686 | 0.886 | 0.006 | -0.018 | -0.008 | 0.041 | 0.050 | 0.059 | 0.009 | -0.002 | -0.031 | 4.833 | 41.161 | 17.375 | -0.050 |
| Iceland | GRET | 2.486 | 49.938 | 227.627 | 8.59 | 0.143 | 0 | -0.019 | -0.008 | 0.011 | 0.033 | 0.022 | 0.041 | -0.010 | -0.007 | -0.017 | 9.789 | 42.668 | 17.368 | -0.034 |
| Iceland | GRJO | 3.925 | 4.34 | 213.518 | 7.45 | 0.286 | 0.657 | -0.016 | -0.008 | 0.009 | 0.001 | 0.013 | 0.013 | -0.004 | 0.010 | 0.001 | 5.278 | 42.144 | 18.111 | 0.002 |
| Iceland | HOLS | 3.569 | 10.519 | 191.938 | 7.53 | 0.657 | 0.571 | -0.013 | -0.016 | 0.012 | -0.004 | 0.000 | 0.016 | -0.005 | -0.006 | 0.021 | 4.353 | 37.641 | 16.941 | -0.138 |
| Iceland | HRED | 3.896 | 6.933 | 206.899 | 7.5 | 0.971 | 0.229 | 0.016 | -0.010 | -0.005 | -0.027 | -0.026 | -0.025 | -0.002 | -0.052 | 0.022 | 4.722 | 42.756 | 17.938 | -0.069 |
| Iceland | KLEI | 9.882 | 10.756 | 186.013 | 7.44 | 0.462 | 0.654 | -0.020 | -0.005 | 0.007 | 0.014 | -0.001 | 0.037 | 0.006 | 0.020 | 0.024 | 6.563 | 39.981 | 19.563 | -0.075 |
| Iceland | MJOA | 5.075 | 4.612 | 213.412 | 7.38 | 1 | 0.857 | 0.012 | -0.009 | 0.005 | 0.021 | 0.034 | 0.045 | 0.000 | 0.034 | -0.011 | 6.053 | 53.216 | 17.526 | -0.026 |
| Iceland | MYVL | 22.461 | 42.501 | 211.865 | 8.47 | 0.914 | 0.371 | -0.012 | 0.001 | 0.011 | -0.004 | 0.000 | 0.015 | 0.011 | 0.060 | -0.003 | 6.111 | 41.317 | 20.000 | 0.039 |
| Iceland | MYVM | 8.348 | 14.686 | 210.338 | 9.6 | 1 | 0.286 | -0.011 | -0.011 | 0.008 | -0.006 | -0.029 | 0.012 | 0.020 | 0.022 | 0.004 | 5.737 | 43.132 | 18.842 | -0.009 |
| Iceland | SAUR | 4.91 | 13.399 | 181.474 | 7.08 | 0.143 | 0.657 | -0.017 | -0.014 | 0.015 | -0.041 | -0.036 | -0.057 | -0.028 | -0.006 | -0.021 | 3.813 | 41.131 | 18.267 | -0.207 |
| Iceland | SKOR | 4.022 | 6.803 | 209.832 | 7.11 | 0.65 | 0.525 | 0.007 | -0.010 | 0.002 | 0.051 | 0.066 | 0.091 | 0.004 | 0.009 | 0.002 | 5.526 | 42.921 | 18.118 | 0.148 |
| Iceland | THIN | 3.735 | 7.124 | 163.347 | 8.72 | 0.914 | 0 | -0.007 | -0.014 | 0.005 | -0.027 | -0.039 | -0.002 | 0.025 | 0.036 | 0.112 | 5.750 | 41.960 | 18.350 | 0.032 |
| Iceland | URR2 | 8.4 | 7.084 | 232.63 | 7.84 | 1 | 0.171 | 0.004 | -0.005 | -0.008 | -0.022 | -0.007 | -0.027 | 0.004 | -0.042 | -0.023 | 4.474 | 38.863 | 17.579 | -0.039 |
| Iceland | URRI | 5.686 | 18.726 | 223.849 | 8.05 | 1 | 0.029 | -0.006 | -0.012 | 0.007 | -0.046 | -0.036 | -0.012 | -0.010 | 0.020 | -0.023 | 4.105 | 37.984 | 17.211 | -0.107 |
| Iceland | VIFI | 9.791 | 19.503 | 201.478 | 9.24 | 0.983 | 0.207 | 0.012 | -0.009 | 0.020 | -0.014 | -0.002 | -0.317 | -0.003 | -0.252 | -0.108 | 2.125 | 37.088 | 18.154 | 0.034 |
| Scotland | AONG | 4.54 | 19.17 | 103.4 | 6.97 | 0.2 | 0.167 | 0.018 | -0.010 | 0.002 | -0.025 | -0.019 | -0.027 | 0.008 | 0.036 | 0.009 | 3.056 | 28.206 | 18.500 | -0.100 |
| Scotland | AROI | 2.897 | 24.32 | 56.18 | 6.48 | 0.657 | 0.029 | 0.038 | 0.001 | 0.000 | 0.032 | 0.024 | 0.067 | 0.001 | 0.028 | 0.005 | 4.000 | 44.716 | 19.000 | 0.146 |
| Scotland | BHAR | 1.472 | 18.63 | 82.63 | 6.03 | 0.429 | 0.514 | 0.028 | -0.006 | 0.000 | -0.025 | -0.175 | -0.286 | -0.033 | -0.291 | -0.246 | 0.333 | 29.217 | 19.250 | -0.058 |
| Scotland | BUAI | 2.549 | 36.14 | 58.44 | 6.73 | 0.029 | 0 | -0.018 | -0.003 | 0.018 | 0.018 | 0.021 | -0.142 | -0.001 | -0.049 | 0.035 | 3.294 | 33.624 | 19.000 | -0.056 |
| Scotland | CHRU | 2.203 | 18.77 | 62.7 | 6.58 | 0 | 0.167 | -0.008 | -0.002 | 0.016 | 0.002 | 0.000 | -0.014 | 0.004 | -0.033 | 0.023 | 3.688 | 33.935 | 19.175 | -0.102 |
| Scotland | DAIM | 2.305 | 24.56 | 56.3 | 6.5 | 0 | 0 | 0.026 | -0.005 | 0.016 | -0.028 | 0.001 | -0.014 | 0.010 | -0.030 | -0.017 | 3.350 | 32.765 | 19.143 | -0.027 |
| Scotland | EISI | 2.813 | 22.54 | 79.14 | 6.82 | 0.543 | 0.143 | 0.011 | -0.005 | 0.004 | 0.001 | 0.000 | -0.026 | 0.005 | -0.007 | 0.021 | 5.375 | 34.665 | 19.667 | -0.082 |
| Scotland | FADA | 1.809 | 17.29 | 90.41 | 6.71 | 0.171 | 0 | 0.025 | -0.003 | -0.004 | -0.142 | -0.149 | -0.277 | -0.181 | -0.295 | -0.263 | 0.067 | 28.640 | 19.175 | 0.091 |
| Scotland | GEIR | 1.762 | 17.17 | 47.29 | 6.7 | 0.743 | 0.057 | 0.020 | -0.009 | 0.001 | -0.011 | -0.011 | -0.028 | 0.021 | 0.028 | -0.082 | 1.824 | 30.747 | 17.250 | -0.059 |
| Scotland | GILL | 2.706 | 21.85 | 46.04 | 6.8 | 0.875 | 0.156 | 0.023 | -0.007 | 0.003 | 0.072 | 0.078 | 0.088 | 0.045 | 0.059 | 0.068 | 4.176 | 38.612 | 19.667 | 0.071 |
| Scotland | HOST | 30.56 | 27.47 | 66.52 | 8.34 | 0.886 | 0.771 | 0.020 | -0.014 | 0.003 | 0.019 | 0.020 | 0.021 | 0.009 | 0.021 | -0.008 | 4.526 | 39.847 | 18.800 | 0.065 |
| Scotland | IALA | 2.728 | 21.4 | 65.9 | 6.36 | 0.143 | 0 | -0.004 | -0.004 | 0.016 | -0.011 | -0.006 | -0.031 | 0.027 | 0.026 | 0.062 | 4.444 | 35.306 | 18.000 | 0.027 |
| Scotland | MORA | 2.505 | 24.03 | 58.85 | 6.35 | 0.265 | 0 | 0.042 | 0.002 | -0.012 | -0.176 | -0.199 | -0.328 | -0.055 | -0.345 | -0.318 | 0.053 | 31.890 | 18.000 | 0.057 |
| Scotland | REIV | 28.6 | 40.12 | 58.52 | 8.95 | 0.514 | 0.057 | -0.005 | 0.010 | 0.011 | 0.005 | 0.006 | 0.026 | 0.030 | 0.061 | 0.053 | 5.474 | 40.995 | 19.600 | -0.022 |
| Scotland | SCAD | 1.418 | 19.75 | 78.89 | 6.14 | 0.571 | 0.086 | 0.038 | 0.002 | 0.001 | -0.135 | -0.157 | -0.354 | -0.067 | -0.367 | -0.345 | 0.176 | 33.571 | 17.000 | -0.017 |
| Scotland | STRU | 3.724 | 25.05 | 93.42 | 7.06 | 0.686 | 0.419 | 0.008 | -0.016 | 0.009 | -0.029 | -0.033 | -0.033 | 0.011 | -0.041 | -0.324 | 0.250 | 32.944 | 20.000 | -0.008 |
| Scotland | TORM | 4.005 | 25.43 | 86.88 | 6.84 | 0.286 | 0.029 | 0.020 | -0.005 | 0.010 | -0.088 | -0.120 | -0.077 | 0.000 | -0.156 | -0.284 | 0.235 | 30.974 | 17.333 | 0.032 |
| Scotland | TROS | 2.845 | 26.12 | 93.32 | 6.71 | 0.686 | 0.03 | 0.026 | -0.004 | 0.004 | 0.061 | 0.065 | 0.053 | 0.024 | 0.066 | 0.042 | 5.350 | 40.400 | 16.400 | 0.082 |

Supplementary Table 3. Loadings of environmental variables (a), shape markers (b) and armour variables (c) for PCAs used in downstream analyses.

|  | PC1 | PC2 | PC3 | PC4 | Total % variance explained |
| --- | --- | --- | --- | --- | --- |
| *a) Environment* |  |  |  |  |  |
| pH | **-0.59018** | -0.18742 | -0.12857 | 0.03533 |  |
| Zn | -0.39596 | **0.58456** | 0.09895 | 0.29584 |  |
| Na | -0.23467 | 0.34844 | **-0.73147** | 0.11473 |  |
| Ca | **-0.43975** | -0.44505 | -0.21176 | -0.514 |  |
| Gyro | -0.39064 | -0.35093 | 0.38023 | 0.58228 |  |
| Schisto | -0.30636 | 0.4249 | 0.49922 | -0.54297 |  |
| variance explained (%) | 35.3 | 26.1 | 18.6 | 11.4 | 91.4 |
| *b) Body shape* |  |  |  |  |  |
| RegResid1 | 0.1145 | -0.2174 | 0.0605 |  |  |
| RegResid2 | 0.0578 | -0.0948 | -0.0382 |  |  |
| RegResid3 | 0.1541 | **-0.4167** | -0.0233 |  |  |
| RegResid4 | -0.0730 | 0.1527 | -0.0933 |  |  |
| RegResid5 | **-0.4788** | 0.0242 | -0.0762 |  |  |
| RegResid6 | 0.1943 | -0.0471 | **0.3102** |  |  |
| RegResid7 | 0.0050 | 0.0072 | **0.6150** |  |  |
| RegResid8 | -0.0283 | 0.0683 | -0.0785 |  |  |
| RegResid9 | **0.2621** | -0.0474 | -0.2446 |  |  |
| RegResid10 | -0.0783 | 0.1055 | -0.0276 |  |  |
| RegResid11 | **0.2685** | 0.0520 | **-0.3039** |  |  |
| RegResid12 | -0.0809 | 0.0793 | -0.0557 |  |  |
| RegResid13 | 0.2529 | -0.0381 | -0.2569 |  |  |
| RegResid14 | -0.0145 | -0.0138 | -0.0399 |  |  |
| RegResid15 | -0.0388 | -0.0017 | **0.4633** |  |  |
| RegResid16 | 0.0177 | -0.1039 | 0.0803 |  |  |
| RegResid17 | **-0.6321** | **-0.3687** | -0.1694 |  |  |
| RegResid18 | 0.0986 | -0.1290 | -0.0687 |  |  |
| RegResid19 | -0.0012 | **0.5694** | -0.0576 |  |  |
| RegResid20 | 0.1164 | -0.2510 | -0.0245 |  |  |
| RegResid21 | 0.0778 | -0.0256 | 0.0001 |  |  |
| RegResid22 | -0.0540 | 0.0180 | -0.0010 |  |  |
| RegResid23 | 0.1145 | 0.1205 | 0.0591 |  |  |
| RegResid24 | -0.0929 | 0.0414 | -0.0470 |  |  |
| RegResid25 | -0.0984 | **0.3423** | -0.0663 |  |  |
| RegResid26 | -0.0630 | 0.1743 | 0.0840 |  |  |
| variance explained (%) | 29.0 | 20.6 | 16.5 |  | 66.1 |
| *c) Armour* |  |  |  |  |  |
| DS1 | **0.4026** | -0.3162 | **0.4398** | 0.20104 |  |
| DS2 | **0.4073** | -0.3276 | **0.4280** | 0.0326 |  |
| PS | **0.4584** | 0.2378 | 0.0388 | 0.15235 |  |
| LP | 0.3490 | **0.5548** | -0.0935 | -0.01418 |  |
| HP | **0.4317** | 0.3787 | -0.1735 | -0.07635 |  |
| BAP | 0.2935 | -0.3236 | -0.2532 | **-0.84338** |  |
| Plate_N | 0.2601 | **-0.4244** | **-0.7204** | **0.4669** |  |
| variance explained (%) | 53.7 | 14.5 | 11.0 | 10.3 | 89.5 |

Supplementary Table 4. Values of θ (angle between vectors) and ΔL (difference in length between vectors) for all pairwise radiation comparisons for environmental, shape, armour and trophic variation

| Multivariate Vector | Radiation pairs | θ (°) | *p*(θ) | Δ*L* |
| --- | --- | --- | --- | --- |
| Environment | Alaska & B.C | 8.709 | 0.077 | 65.28 |
| Environment | Alaska & Iceland | 26.814 | 0.828 | 18.215 |
| Environment | Alaska & Scotland | 12.261 | 0.388 | 36.553 |
| Environment | B.C & Iceland | 18.392 | 0.245 | 47.064 |
| Environment | B.C & Scotland | 9.167 | 0.066 | 28.727 |
| Environment | Iceland & Scotland | 18.335 | 0.826 | 18.337 |
| Shape | Alaska & B.C | 3.495 | 0.19 | 0.239 |
| Shape | Alaska & Iceland | 9.454 | 0.707 | 0.232 |
| Shape | Alaska & Scotland | 15.871 | 0.925 | 0.627 |
| Shape | B.C & Iceland | 12.892 | 0.834 | 0.008 |
| Shape | B.C & Scotland | 18.85 | 0.947 | 0.388 |
| Shape | Iceland & Scotland | 8.48 | 0.627 | 0.396 |
| Armour | Alaska & B.C | 6.36 | **0.004** | 99.143 |
| Armour | Alaska & Iceland | 8.365 | **0.033** | 155.091 |
| Armour | Alaska & Scotland | 6.818 | **0.021** | 54.474 |
| Armour | B.C & Iceland | 4.526 | **0.013** | 254.233 |
| Armour | B.C & Scotland | 4.125 | **0.004** | 44.669 |
| Armour | Iceland & Scotland | 8.308 | **0.053** | 209.565 |
| Gill Raker | Alaska & B.C | 1.49 | 0.112 | 1.917 |
| Gill Raker | Alaska & Iceland | 8.537 | 0.462 | 0.237 |
| Gill Raker | Alaska & Scotland | 5.787 | 0.365 | 0.436 |
| Gill Raker | B.C & Iceland | 7.047 | 0.391 | 2.155 |
| Gill Raker | B.C & Scotland | 7.334 | 0.459 | 2.353 |
| Gill Raker | Iceland & Scotland | 14.381 | 0.462 | 0.198 |

|

Supplementary Table 5. Results of ANOVAs for two first PCs of armour and three PCs of body shape for fish from different lakes and radiations. Signiﬁcant P-values in bold

|  | Df | Sum Sq | Mean Sq | F-value | Pr(>F) |
| --- | --- | --- | --- | --- | --- |
| *Armour_PC1* | |  |  |  |  |
| Radiation | 3 | 331 | 110.36 | 180.36 | **<0.001** |
| Lake | 69 | 3805 | 55.14 | 90.12 | **<0.001** |
| Residuals | 1226 | 750 | 0.61 |  |  |
| *Armour_PC2* | |  |  |  |  |
| Radiation | 3 | 146.8 | 48.94 | 166.12 | **<0.001** |
| Lake | 69 | 807.5 | 11.7 | 39.72 | **<0.001** |
| Residuals | 1226 | 361.2 | 0.29 |  |  |
| *Shape_PC1* | |  |  |  |  |
| Radiation | 3 | 0.13962 | 0.04654 | 686.77 | **<0.001** |
| Lake | 69 | 0.17873 | 0.00259 | 38.22 | **<0.001** |
| Residuals | 1223 | 0.08288 | 0.00007 |  |  |
| *Shape_PC2* | |  |  |  |  |
| Radiation | 3 | 0.06767 | 0.022555 | 173.817 | **<0.001** |
| Lake | 69 | 0.05938 | 0.000861 | 6.632 | **<0.001** |
| Residuals | 1223 | 0.1587 | 0.00013 |  |  |
| *Shape_PC3* | |  |  |  |  |
| Radiation | 3 | 0.0542 | 0.018067 | 232.41 | **<0.001** |
| Lake | 69 | 0.07891 | 0.001144 | 14.71 | **<0.001** |
| Residuals | 1223 | 0.09507 | 0.000078 |  |  |

Supplementary Table 6. Results of ANCOVAs for PC1 and PC2 of body shape and armour, and for gill raker number and length. Signiﬁcant P-values in bold.

|  | Df | Sum Sq | Mean Sq | F-value | Pr(>F) |
| --- | --- | --- | --- | --- | --- |
| ***a) shapePC1*** |  |  |  |  |  |
| EnvPC1 | 1 | 0.00076 | 0.00076 | 4.852 | **0.031** |
| EnvPC2 | 1 | 0.001183 | 0.001184 | 7.56 | **0.008** |
| Radiation | 3 | 0.006058 | 0.002019 | 12.9 | **0.000** |
| EnvPC1:Radiation | 3 | 0.000177 | 5.91E-05 | 0.378 | 0.769 |
| EnvPC2:Radiation | 3 | 0.000037 | 1.22E-05 | 0.078 | 0.972 |
| Residuals | 61 | 0.009549 | 0.000157 |  |  |
| ***b) shapePC2*** |  |  |  |  |  |
| EnvPC1 | 1 | 0.001238 | 0.001238 | 28.197 | **0.000** |
| EnvPC2 | 1 | 0.002108 | 0.002108 | 48.014 | **0.000** |
| Radiation | 3 | 0.000622 | 0.000207 | 4.724 | **0.005** |
| EnvPC1:Radiation | 3 | 0.000358 | 0.000119 | 2.718 | 0.052 |
| EnvPC2:Radiation | 3 | 0.000306 | 0.000102 | 2.326 | 0.084 |
| Residuals | 61 | 0.002679 | 4.39E-05 |  |  |
| ***c) armourPC1*** |  |  |  |  |  |
| EnvPC1 | 1 | 17.16 | 17.157 | 6.13 | **0.016** |
| EnvPC2 | 1 | 3.15 | 3.151 | 1.126 | 0.293 |
| Radiation | 3 | 17.71 | 5.905 | 2.11 | 0.108 |
| EnvPC1:Radiation | 3 | 17.46 | 5.82 | 2.079 | 0.112 |
| EnvPC2:Radiation | 3 | 6.27 | 2.089 | 0.746 | 0.529 |
| Residuals | 61 | 170.72 | 2.799 |  |  |
| ***d) armourPC2*** |  |  |  |  |  |
| EnvPC1 | 1 | 2.49 | 2.4857 | 4.108 | **0.047** |
| EnvPC2 | 1 | 0.67 | 0.666 | 1.101 | 0.298 |
| Radiation | 3 | 9.24 | 3.0795 | 5.09 | **0.003** |
| EnvPC1:Radiation | 3 | 5.01 | 1.6689 | 2.758 | **0.050** |
| EnvPC2:Radiation | 3 | 0.24 | 0.0805 | 0.133 | 0.940 |
| Residuals | 61 | 36.91 | 0.6051 |  |  |
| ***e) Gill.Raker.N*** |  |  |  |  |  |
| EnvPC1 | 1 | 22.16 | 22.16 | 18.167 | **0.000** |
| EnvPC2 | 1 | 53.37 | 53.37 | 43.744 | **0.000** |
| Radiation | 3 | 24.71 | 8.24 | 6.752 | **0.001** |
| EnvPC1:Radiation | 3 | 14.7 | 4.9 | 4.015 | **0.011** |
| EnvPC2:Radiation | 3 | 3.42 | 1.14 | 0.935 | 0.430 |
| Residuals | 61 | 74.42 | 1.22 |  |  |
| ***f) Resid.Raker.L*** |  |  |  |  |  |
| EnvPC1 | 1 | 0.0009 | 0.000911 | 0.088 | 0.768 |
| EnvPC2 | 1 | 0.004 | 0.003961 | 0.381 | 0.540 |
| Radiation | 3 | 0.0026 | 0.000853 | 0.082 | 0.970 |
| EnvPC1:Radiation | 3 | 0.0923 | 0.030779 | 2.958 | **0.039** |
| EnvPC2:Radiation | 3 | 0.045 | 0.014999 | 1.442 | 0.239 |
| Residuals | 61 | 0.6347 | 0.010405 |  |  |

Supplementary Table 7. F_ST_ values among the four countries based on 8,395 SNPs with no Chr XIX. Within radiations, individuals from different populations were pooled in order to estimate the F_ST_ values between radiations.

|  | Iceland | B.C. | Alaska |
| --- | --- | --- | --- |
| Scotland | 0.194 | 0.338 | 0.329 |
| Iceland |  | 0.323 | 0.314 |
| B.C. |  |  | 0.198 |

Supplementary Table 8. Results of AMOVA over all populations. Individuals were nested within populations (Pop), within radiations (Rad), within continents.

|  | Sigma | % |
| --- | --- | --- |
| Variations Between Continent | 889.741 | 34.748 |
| Variations Between Rad Within Continent | 142.320 | 5.558 |
| Variations Between Pop Within Rad | 366.516 | 14.314 |
| Variations Between samples Within Pop | 228.970 | 8.942 |
| Variations Within samples | 932.972 | 36.437 |
| Total variations | 2560.519 | 100 |

Supplementary Table 9. Numbers of SNPs and windows associated with each of 12 phenotypic traits and 6 environmental variables in the four independent adaptive radiations.

| Location | Variable | Associated SNP N | Associated Window N | | | |  |
| --- | --- | --- | --- | --- | --- | --- | --- |
|  |  |  | 50k | 75k | 100k | 200k | 0.1 cM |
| Alaska | Ca | 715 | 84 | 80 | 74 | 51 | 56 |
| Alaska | Gyro | 166 | 27 | 27 | 24 | 23 | 23 |
| Alaska | Na | 205 | 40 | 34 | 35 | 31 | 29 |
| Alaska | pH | 305 | 42 | 42 | 40 | 35 | 31 |
| Alaska | Schisto | 226 | 43 | 43 | 37 | 33 | 33 |
| Alaska | Zn | 285 | 43 | 42 | 41 | 34 | 35 |
| Alaska | Shape_PC1 | 152 | 36 | 36 | 33 | 32 | 23 |
| Alaska | Shape_PC2 | 242 | 52 | 45 | 39 | 30 | 34 |
| Alaska | Shape_PC3 | 205 | 43 | 42 | 37 | 29 | 31 |
| Alaska | DS1 | 259 | 50 | 43 | 43 | 37 | 31 |
| Alaska | DS2 | 179 | 33 | 30 | 29 | 28 | 31 |
| Alaska | PS | 470 | 63 | 58 | 59 | 41 | 43 |
| Alaska | LP | 682 | 81 | 70 | 63 | 47 | 59 |
| Alaska | HP | 544 | 65 | 60 | 48 | 36 | 47 |
| Alaska | BAP | 453 | 62 | 56 | 56 | 41 | 41 |
| Alaska | Plate_N | 424 | 69 | 65 | 59 | 40 | 42 |
| Alaska | Gill_Raker_L | 344 | 57 | 54 | 49 | 39 | 41 |
| Alaska | Gill_Raker_N | 104 | 28 | 23 | 20 | 20 | 19 |
| BC | Ca | 750 | 75 | 68 | 60 | 41 | 46 |
| BC | Gyro | 457 | 55 | 61 | 53 | 44 | 36 |
| BC | Na | 921 | 136 | 123 | 109 | 96 | 77 |
| BC | pH | 620 | 61 | 61 | 53 | 33 | 40 |
| BC | Schisto | 243 | 38 | 38 | 33 | 28 | 34 |
| BC | Zn | 96 | 30 | 27 | 23 | 19 | 20 |
| BC | Shape_PC1 | 116 | 21 | 16 | 12 | 10 | 15 |
| BC | Shape_PC2 | 433 | 43 | 45 | 40 | 30 | 31 |
| BC | Shape_PC3 | 365 | 45 | 45 | 45 | 36 | 36 |
| BC | DS1 | 966 | 130 | 120 | 113 | 91 | 85 |
| BC | DS2 | 875 | 134 | 122 | 115 | 94 | 83 |
| BC | PS | 991 | 146 | 136 | 126 | 98 | 90 |
| BC | LP | 942 | 145 | 130 | 119 | 101 | 84 |
| BC | HP | 641 | 93 | 89 | 77 | 64 | 61 |
| BC | BAP | 386 | 54 | 52 | 47 | 31 | 39 |
| BC | Plate_N | 709 | 129 | 114 | 101 | 79 | 72 |
| BC | Gill_Raker_L | 617 | 64 | 69 | 62 | 52 | 46 |
| BC | Gill_Raker_N | 936 | 138 | 118 | 113 | 86 | 74 |
| Iceland | Ca | 142 | 32 | 30 | 29 | 21 | 20 |
| Iceland | Gyro | 380 | 45 | 44 | 34 | 26 | 36 |
| Iceland | Na | 180 | 38 | 34 | 29 | 21 | 27 |
| Iceland | pH | 391 | 59 | 57 | 53 | 40 | 43 |
| Iceland | Schisto | 432 | 56 | 57 | 53 | 40 | 46 |
| Iceland | Zn | 395 | 48 | 48 | 40 | 24 | 37 |
| Iceland | Shape_PC1 | 569 | 59 | 51 | 48 | 32 | 46 |
| Iceland | Shape_PC2 | 200 | 39 | 34 | 30 | 22 | 31 |
| Iceland | Shape_PC3 | 215 | 37 | 33 | 27 | 21 | 28 |
| Iceland | DS1 | 158 | 29 | 28 | 25 | 23 | 24 |
| Iceland | DS2 | 148 | 26 | 24 | 20 | 18 | 21 |
| Iceland | PS | 511 | 53 | 51 | 43 | 27 | 38 |
| Iceland | LP | 306 | 38 | 40 | 40 | 29 | 32 |
| Iceland | HP | 252 | 49 | 39 | 43 | 28 | 32 |
| Iceland | BAP | 332 | 36 | 37 | 31 | 21 | 30 |
| Iceland | Plate_N | 194 | 41 | 37 | 34 | 28 | 26 |
| Iceland | Gill_Raker_L | 209 | 38 | 34 | 34 | 27 | 27 |
| Iceland | Gill_Raker_N | 296 | 45 | 46 | 46 | 42 | 35 |
| Scotland | Ca | 324 | 36 | 37 | 37 | 30 | 30 |
| Scotland | Gyro | 301 | 23 | 23 | 20 | 16 | 17 |
| Scotland | Na | 495 | 43 | 38 | 43 | 31 | 34 |
| Scotland | pH | 401 | 42 | 40 | 38 | 30 | 29 |
| Scotland | Schisto | 541 | 46 | 35 | 39 | 28 | 28 |
| Scotland | Zn | 87 | 26 | 19 | 17 | 10 | 20 |
| Scotland | Shape_PC1 | 127 | 33 | 28 | 20 | 15 | 16 |
| Scotland | Shape_PC2 | 697 | 51 | 54 | 48 | 40 | 50 |
| Scotland | Shape_PC3 | 47 | 23 | 20 | 12 | 6 | 20 |
| Scotland | DS1 | 154 | 24 | 23 | 19 | 17 | 22 |
| Scotland | DS2 | 197 | 34 | 34 | 25 | 19 | 33 |
| Scotland | PS | 385 | 37 | 33 | 31 | 33 | 28 |
| Scotland | LP | 204 | 35 | 30 | 26 | 25 | 20 |
| Scotland | HP | 394 | 49 | 43 | 44 | 34 | 25 |
| Scotland | BAP | 275 | 34 | 34 | 37 | 28 | 40 |
| Scotland | Plate_N | 334 | 41 | 37 | 43 | 36 | 33 |
| Scotland | Gill_Raker_L | 44 | 25 | 15 | 16 | 8 | 15 |
| Scotland | Gill_Raker_N | 351 | 44 | 40 | 40 | 32 | 27 |

Supplementary Table 10. Results for permuted (10,000 runs) parallelism across all radiations. Results are shown for each group of radiations compared, with the permuted expected, observed parallel, and FDR-corrected p-value. Significant parallelism is assumed when FDR < 0.05.

| Variable | Comparison | Expected | Observed | FDR |
| --- | --- | --- | --- | --- |
| Ca | All 4 | 0 | 0 | 1 |
| Ca | Alaska & BC & Iceland | 0.004 | 0 | 1 |
| Ca | Alaska & BC & Scotland | 0.0042 | 0 | 1 |
| Ca | Alaska & Iceland & Scotland | 0.0023 | 0 | 1 |
| Ca | BC & Iceland & Scotland | 0.0023 | 0 | 1 |
| Ca | Alaska & BC | 0.9326 | 6 | ***<0.001*** |
| Ca | Alaska & Iceland | 0.3696 | 3 | ***0.0062*** |
| Ca | Alaska & Scotland | 0.4136 | 1 | 0.3376 |
| Ca | BC & Iceland | 0.3289 | 0 | 1 |
| Ca | BC & Scotland | 0.3736 | 1 | 0.3131 |
| Ca | Iceland & Scotland | 0.1753 | 3 | ***<0.001*** |
| Gyro | All 4 | 0 | 0 | 1 |
| Gyro | Alaska & BC & Iceland | 0.001 | 0 | 1 |
| Gyro | Alaska & BC & Scotland | 0.0011 | 0 | 1 |
| Gyro | Alaska & Iceland & Scotland | 2.00E-04 | 0 | 1 |
| Gyro | BC & Iceland & Scotland | 0.0013 | 0 | 1 |
| Gyro | Alaska & BC | 0.2043 | 2 | ***0.0178*** |
| Gyro | Alaska & Iceland | 0.1718 | 0 | 1 |
| Gyro | Alaska & Scotland | 0.0837 | 0 | 1 |
| Gyro | BC & Iceland | 0.3415 | 1 | 0.2907 |
| Gyro | BC & Scotland | 0.1788 | 1 | 0.1632 |
| Gyro | Iceland & Scotland | 0.1579 | 0 | 1 |
| Na | All 4 | 0 | 0 | 1 |
| Na | Alaska & BC & Iceland | 0.0049 | 0 | 1 |
| Na | Alaska & BC & Scotland | 0.004 | 0 | 1 |
| Na | Alaska & Iceland & Scotland | 0.0016 | 0 | 1 |
| Na | BC & Iceland & Scotland | 0.0045 | 0 | 1 |
| Na | Alaska & BC | 0.7949 | 1 | 0.553 |
| Na | Alaska & Iceland | 0.2127 | 0 | 1 |
| Na | Alaska & Scotland | 0.241 | 0 | 1 |
| Na | BC & Iceland | 0.7198 | 1 | 0.5143 |
| Na | BC & Scotland | 0.815 | 1 | 0.5653 |
| Na | Iceland & Scotland | 0.2369 | 1 | 0.2111 |
| pH | All 4 | 1.00E-04 | 0 | 1 |
| pH | Alaska & BC & Iceland | 0.0029 | 0 | 1 |
| pH | Alaska & BC & Scotland | 0.0014 | 0 | 1 |
| pH | Alaska & Iceland & Scotland | 0.002 | 0 | 1 |
| pH | BC & Iceland & Scotland | 0.0032 | 0 | 1 |
| pH | Alaska & BC | 0.3811 | 1 | 0.3201 |
| pH | Alaska & Iceland | 0.3486 | 0 | 1 |
| pH | Alaska & Scotland | 0.2438 | 0 | 1 |
| pH | BC & Iceland | 0.4938 | 2 | 0.0882 |
| pH | BC & Scotland | 0.3557 | 1 | 0.3053 |
| pH | Iceland & Scotland | 0.3679 | 5 | ***<0.001*** |
| Schisto | All 4 | 1.00E-04 | 0 | 1 |
| Schisto | Alaska & BC & Iceland | 0.0015 | 0 | 1 |
| Schisto | Alaska & BC & Scotland | 0.0015 | 0 | 1 |
| Schisto | Alaska & Iceland & Scotland | 0.0022 | 0 | 1 |
| Schisto | BC & Iceland & Scotland | 0.0019 | 0 | 1 |
| Schisto | Alaska & BC | 0.233 | 1 | 0.2092 |
| Schisto | Alaska & Iceland | 0.3285 | 1 | 0.285 |
| Schisto | Alaska & Scotland | 0.2816 | 0 | 1 |
| Schisto | BC & Iceland | 0.2961 | 2 | ***0.0337*** |
| Schisto | BC & Scotland | 0.2386 | 1 | 0.2114 |
| Schisto | Iceland & Scotland | 0.3978 | 0 | 1 |
| Zn | All 4 | 0 | 0 | 1 |
| Zn | Alaska & BC & Iceland | 0.0013 | 0 | 1 |
| Zn | Alaska & BC & Scotland | 7.00E-04 | 0 | 1 |
| Zn | Alaska & Iceland & Scotland | 0.0011 | 0 | 1 |
| Zn | BC & Iceland & Scotland | 0.0012 | 0 | 1 |
| Zn | Alaska & BC | 0.1864 | 1 | 0.1699 |
| Zn | Alaska & Iceland | 0.2873 | 1 | 0.2493 |
| Zn | Alaska & Scotland | 0.1539 | 0 | 1 |
| Zn | BC & Iceland | 0.2042 | 0 | 1 |
| Zn | BC & Scotland | 0.1026 | 1 | 0.0963 |
| Zn | Iceland & Scotland | 0.183 | 0 | 1 |
| Shape_PC1 | All 4 | 0 | 0 | 1 |
| Shape_PC1 | Alaska & BC & Iceland | 9.00E-04 | 0 | 1 |
| Shape_PC1 | Alaska & BC & Scotland | 4.00E-04 | 0 | 1 |
| Shape_PC1 | Alaska & Iceland & Scotland | 0.0012 | 0 | 1 |
| Shape_PC1 | BC & Iceland & Scotland | 4.00E-04 | 0 | 1 |
| Shape_PC1 | Alaska & BC | 0.1128 | 0 | 1 |
| Shape_PC1 | Alaska & Iceland | 0.2974 | 0 | 1 |
| Shape_PC1 | Alaska & Scotland | 0.1724 | 1 | 0.1584 |
| Shape_PC1 | BC & Iceland | 0.1827 | 1 | 0.1674 |
| Shape_PC1 | BC & Scotland | 0.0957 | 0 | 1 |
| Shape_PC1 | Iceland & Scotland | 0.2974 | 0 | 1 |
| Shape_PC2 | All 4 | 0 | 0 | 1 |
| Shape_PC2 | Alaska & BC & Iceland | 0.0014 | 0 | 1 |
| Shape_PC2 | Alaska & BC & Scotland | 0.0026 | 0 | 1 |
| Shape_PC2 | Alaska & Iceland & Scotland | 0.0034 | 0 | 1 |
| Shape_PC2 | BC & Iceland & Scotland | 0.0016 | 0 | 1 |
| Shape_PC2 | Alaska & BC | 0.3278 | 1 | 0.2836 |
| Shape_PC2 | Alaska & Iceland | 0.2918 | 0 | 1 |
| Shape_PC2 | Alaska & Scotland | 0.3743 | 0 | 1 |
| Shape_PC2 | BC & Iceland | 0.227 | 1 | 0.2044 |
| Shape_PC2 | BC & Scotland | 0.298 | 0 | 1 |
| Shape_PC2 | Iceland & Scotland | 0.3017 | 1 | 0.2645 |
| Shape_PC3 | All 4 | 0 | 0 | 1 |
| Shape_PC3 | Alaska & BC & Iceland | 0.0012 | 0 | 1 |
| Shape_PC3 | Alaska & BC & Scotland | 0.0011 | 0 | 1 |
| Shape_PC3 | Alaska & Iceland & Scotland | 4.00E-04 | 0 | 1 |
| Shape_PC3 | BC & Iceland & Scotland | 6.00E-04 | 0 | 1 |
| Shape_PC3 | Alaska & BC | 0.2838 | 1 | 0.245 |
| Shape_PC3 | Alaska & Iceland | 0.2304 | 0 | 1 |
| Shape_PC3 | Alaska & Scotland | 0.1395 | 0 | 1 |
| Shape_PC3 | BC & Iceland | 0.2388 | 1 | 0.2134 |
| Shape_PC3 | BC & Scotland | 0.1417 | 0 | 1 |
| Shape_PC3 | Iceland & Scotland | 0.1279 | 0 | 1 |
| Shape_PC3 | All 4 | 0 | 0 | 1 |
| DS1 | Alaska & BC & Iceland | 0.0044 | 0 | 1 |
| DS1 | Alaska & BC & Scotland | 0.0036 | 0 | 1 |
| DS1 | Alaska & Iceland & Scotland | 3.00E-04 | 0 | 1 |
| DS1 | BC & Iceland & Scotland | 0.0017 | 0 | 1 |
| DS1 | Alaska & BC | 0.9636 | 0 | 1 |
| DS1 | Alaska & Iceland | 0.1978 | 0 | 1 |
| DS1 | Alaska & Scotland | 0.1594 | 1 | 0.1489 |
| DS1 | BC & Iceland | 0.5224 | 1 | 0.4124 |
| DS1 | BC & Scotland | 0.4347 | 1 | 0.3561 |
| DS1 | Iceland & Scotland | 0.1044 | 1 | 0.1002 |
| DS2 | All 4 | 0 | 0 | 1 |
| DS2 | Alaska & BC & Iceland | 0.0023 | 0 | 1 |
| DS2 | Alaska & BC & Scotland | 0.0031 | 0 | 1 |
| DS2 | Alaska & Iceland & Scotland | 0.0011 | 0 | 1 |
| DS2 | BC & Iceland & Scotland | 0.0023 | 0 | 1 |
| DS2 | Alaska & BC | 0.6511 | 1 | 0.4818 |
| DS2 | Alaska & Iceland | 0.119 | 0 | 1 |
| DS2 | Alaska & Scotland | 0.1507 | 1 | 0.1414 |
| DS2 | BC & Iceland | 0.4835 | 2 | 0.0797 |
| DS2 | BC & Scotland | 0.6224 | 0 | 1 |
| DS2 | Iceland & Scotland | 0.1308 | 3 | ***<0.001*** |
| PS | All 4 | 1.00E-04 | 0 | 1 |
| PS | Alaska & BC & Iceland | 0.0102 | 0 | 1 |
| PS | Alaska & BC & Scotland | 0.0089 | 0 | 1 |
| PS | Alaska & Iceland & Scotland | 0.0023 | 0 | 1 |
| PS | BC & Iceland & Scotland | 0.0069 | 0 | 1 |
| PS | Alaska & BC | 1.3527 | 4 | ***0.0426*** |
| PS | Alaska & Iceland | 0.4547 | 1 | 0.3668 |
| PS | Alaska & Scotland | 0.3244 | 1 | 0.2805 |
| PS | BC & Iceland | 1.0713 | 2 | 0.2857 |
| PS | BC & Scotland | 0.7441 | 3 | ***0.0385*** |
| PS | Iceland & Scotland | 0.292 | 0 | 1 |
| LP | All 4 | 0 | 0 | 1 |
| LP | Alaska & BC & Iceland | 0.0108 | 0 | 1 |
| LP | Alaska & BC & Scotland | 0.0086 | 0 | 1 |
| LP | Alaska & Iceland & Scotland | 0.0025 | 0 | 1 |
| LP | BC & Iceland & Scotland | 0.0037 | 1 | ***0.0037*** |
| LP | Alaska & BC | 1.7366 | 2 | 0.5181 |
| LP | Alaska & Iceland | 0.4221 | 4 | ***<0.001*** |
| LP | Alaska & Scotland | 0.3913 | 2 | 0.06 |
| LP | BC & Iceland | 0.768 | 0 | 1 |
| LP | BC & Scotland | 0.7015 | 1 | 0.5095 |
| LP | Iceland & Scotland | 0.1927 | 0 | 1 |
| HP | All 4 | 0 | 0 | 1 |
| HP | Alaska & BC & Iceland | 0.0059 | 0 | 1 |
| HP | Alaska & BC & Scotland | 0.0066 | 0 | 1 |
| HP | Alaska & Iceland & Scotland | 0.003 | 0 | 1 |
| HP | BC & Iceland & Scotland | 0.0043 | 0 | 1 |
| HP | Alaska & BC | 0.8742 | 1 | 0.5904 |
| HP | Alaska & Iceland | 0.4447 | 0 | 1 |
| HP | Alaska & Scotland | 0.44 | 0 | 1 |
| HP | BC & Iceland | 0.6348 | 1 | 0.4715 |
| HP | BC & Scotland | 0.6269 | 0 | 1 |
| HP | Iceland & Scotland | 0.3663 | 1 | 0.3061 |
| BAP | All 4 | 0 | 0 | 1 |
| BAP | Alaska & BC & Iceland | 0.0025 | 0 | 1 |
| BAP | Alaska & BC & Scotland | 0.0025 | 0 | 1 |
| BAP | Alaska & Iceland & Scotland | 0.0018 | 0 | 1 |
| BAP | BC & Iceland & Scotland | 0.0018 | 0 | 1 |
| BAP | Alaska & BC | 0.4878 | 0 | 1 |
| BAP | Alaska & Iceland | 0.3112 | 0 | 1 |
| BAP | Alaska & Scotland | 0.2994 | 0 | 1 |
| BAP | BC & Iceland | 0.281 | 0 | 1 |
| BAP | BC & Scotland | 0.2529 | 0 | 1 |
| BAP | Iceland & Scotland | 0.182 | 0 | 1 |
| Plate_N | All 4 | 0 | 0 | 1 |
| Plate_N | Alaska & BC & Iceland | 0.0066 | 0 | 1 |
| Plate_N | Alaska & BC & Scotland | 0.0076 | 0 | 1 |
| Plate_N | Alaska & Iceland & Scotland | 0.003 | 0 | 1 |
| Plate_N | BC & Iceland & Scotland | 0.005 | 0 | 1 |
| Plate_N | Alaska & BC | 1.2927 | 8 | ***0*** |
| Plate_N | Alaska & Iceland | 0.4013 | 0 | 1 |
| Plate_N | Alaska & Scotland | 0.3859 | 0 | 1 |
| Plate_N | BC & Iceland | 0.7269 | 2 | 0.1613 |
| Plate_N | BC & Scotland | 0.7217 | 0 | 1 |
| Plate_N | Iceland & Scotland | 0.2549 | 1 | 0.2278 |
| Gill_Raker_L | All 4 | 0 | 0 | 1 |
| Gill_Raker_L | Alaska & BC & Iceland | 0.0032 | 0 | 1 |
| Gill_Raker_L | Alaska & BC & Scotland | 0.0016 | 0 | 1 |
| Gill_Raker_L | Alaska & Iceland & Scotland | 0.0013 | 0 | 1 |
| Gill_Raker_L | BC & Iceland & Scotland | 0.0019 | 0 | 1 |
| Gill_Raker_L | Alaska & BC | 0.5381 | 1 | 0.4175 |
| Gill_Raker_L | Alaska & Iceland | 0.3034 | 1 | 0.2626 |
| Gill_Raker_L | Alaska & Scotland | 0.1997 | 0 | 1 |
| Gill_Raker_L | BC & Iceland | 0.3398 | 0 | 1 |
| Gill_Raker_L | BC & Scotland | 0.2202 | 1 | 0.1997 |
| Gill_Raker_L | Iceland & Scotland | 0.1408 | 0 | 1 |
| Gill_Raker_N | All 4 | 0 | 0 | 1 |
| Gill_Raker_N | Alaska & BC & Iceland | 0.0044 | 0 | 1 |
| Gill_Raker_N | Alaska & BC & Scotland | 0.0029 | 0 | 1 |
| Gill_Raker_N | Alaska & Iceland & Scotland | 0.0011 | 0 | 1 |
| Gill_Raker_N | BC & Iceland & Scotland | 0.0056 | 0 | 1 |
| Gill_Raker_N | Alaska & BC | 0.5578 | 7 | ***<0.001*** |
| Gill_Raker_N | Alaska & Iceland | 0.1854 | 0 | 1 |
| Gill_Raker_N | Alaska & Scotland | 0.1688 | 0 | 1 |
| Gill_Raker_N | BC & Iceland | 0.8826 | 0 | 1 |
| Gill_Raker_N | BC & Scotland | 0.8335 | 1 | 0.5665 |
| Gill_Raker_N | Iceland & Scotland | 0.2933 | 1 | 0.2564 |

Supplementary Table 11. 50kb Marine - Freshwater (MxF) outlier regions. Parallel windows are grouped and windows are ordered by extent of parallelism. Previous evidence of marine x freshwater association is also highlighted.

| window_id | Radiations | Previous Marine Freshwater evidence? |
| --- | --- | --- |
| groupXX:8950000-9000000 | Alaska & BC & Iceland & Scotland | No |
| groupI:21600000-21650000 | Alaska & Iceland & Scotland | Jones et al 2012 Nature/Hohenlohe et al 2010 PLoS Genet |
| groupI:21750000-21800000 | Alaska & Iceland & Scotland | Jones et al 2012 Nature/Hohenlohe et al 2010 PLoS Genet |
| groupIV:12800000-12850000 | Alaska & Iceland & Scotland | Jones et al 2012 Nature/Hohenlohe et al 2010 PLoS Genet |
| groupI:8600000-8650000 | Alaska & Iceland | No |
| groupI:21850000-21900000 | Alaska & Scotland | Jones et al 2012 Nature/Hohenlohe et al 2010 PLoS Genet |
| groupIV:19900000-19950000 | Alaska & Iceland | Jones et al 2012 Nature/Hohenlohe et al 2010 PLoS Genet |
| groupVIII:9350000-9400000 | Alaska & BC | No |
| groupXI:5450000-5500000 | Alaska & BC | Hohenlohe et al 2010 PLoS Genet |
| groupXI:5550000-5600000 | Alaska & BC | Hohenlohe et al 2010 PLoS Genet |
| groupXI:5750000-5800000 | Alaska & BC | Hohenlohe et al 2010 PLoS Genet |
| scaffold-37:0-50000 | Alaska & BC | No |
| scaffold-37:100000-150000 | Alaska & BC | No |
| scaffold-37:150000-200000 | Alaska & BC | No |
| scaffold-37:200000-250000 | Alaska & BC | No |
| groupIV:19850000-19900000 | BC & Iceland | Jones et al 2012 Nature/Hohenlohe et al 2010 PLoS Genet |
| groupI:21550000-21600000 | Iceland & Scotland | Jones et al 2012 Nature/Hohenlohe et al 2010 PLoS Genet |
| groupIV:23950000-24000000 | Iceland & Scotland | Jones et al 2012 Nature/Hohenlohe et al 2010 PLoS Genet |
| groupXV:7150000-7200000 | Iceland & Scotland | No |
| groupXX:250000-300000 | Iceland & Scotland | No |
| scaffold-47:1600000-1650000 | Iceland & Scotland | No |
| groupI:9350000-9400000 | Alaska | No |
| groupI:21800000-21850000 | Alaska | Jones et al 2012 Nature |
| groupII:4550000-4600000 | Alaska | No |
| groupIV:10450000-10500000 | Alaska | No |
| groupIV:10550000-10600000 | Alaska | No |
| groupIV:11000000-11050000 | Alaska | No |
| groupIV:11200000-11250000 | Alaska | No |
| groupIV:12750000-12800000 | Alaska | Hohenlohe et al 2010 PLoS Genet |
| groupIV:12900000-12950000 | Alaska | Hohenlohe et al 2010 PLoS Genet |
| groupIV:13800000-13850000 | Alaska | No |
| groupIV:16750000-16800000 | Alaska | No |
| groupIV:18100000-18150000 | Alaska | No |
| groupIV:19250000-19300000 | Alaska | No |
| groupIV:19800000-19850000 | Alaska | Jones et al 2012 Nature/Hohenlohe et al 2010 PLoS Genet |
| groupIV:19950000-20000000 | Alaska | Hohenlohe et al 2010 PLoS Genet |
| groupIV:20000000-20050000 | Alaska | Hohenlohe et al 2010 PLoS Genet |
| groupIV:20050000-20100000 | Alaska | Hohenlohe et al 2010 PLoS Genet |
| groupIV:23800000-23850000 | Alaska | Hohenlohe et al 2010 PLoS Genet |
| groupIV:25100000-25150000 | Alaska | Hohenlohe et al 2010 PLoS Genet |
| groupIV:25750000-25800000 | Alaska | No |
| groupIV:25800000-25850000 | Alaska | No |
| groupIV:26150000-26200000 | Alaska | No |
| groupIV:26200000-26250000 | Alaska | No |
| groupIV:26300000-26350000 | Alaska | No |
| groupIV:26400000-26450000 | Alaska | No |
| groupIX:8550000-8600000 | Alaska | No |
| groupIX:12450000-12500000 | Alaska | No |
| groupIX:12500000-12550000 | Alaska | No |
| groupIX:12800000-12850000 | Alaska | No |
| groupV:4350000-4400000 | Alaska | No |
| groupV:6200000-6250000 | Alaska | No |
| groupVII:7350000-7400000 | Alaska | No |
| groupVII:9300000-9350000 | Alaska | No |
| groupVII:9400000-9450000 | Alaska | No |
| groupVII:10100000-10150000 | Alaska | No |
| groupVII:10150000-10200000 | Alaska | No |
| groupVII:12500000-12550000 | Alaska | No |
| groupVII:13450000-13500000 | Alaska | No |
| groupVII:14350000-14400000 | Alaska | Hohenlohe et al 2010 PLoS Genet |
| groupVII:14400000-14450000 | Alaska | Hohenlohe et al 2010 PLoS Genet |
| groupVII:14650000-14700000 | Alaska | Hohenlohe et al 2010 PLoS Genet |
| groupVII:14700000-14750000 | Alaska | Hohenlohe et al 2010 PLoS Genet |
| groupVII:16800000-16850000 | Alaska | Hohenlohe et al 2010 PLoS Genet |
| groupVII:17050000-17100000 | Alaska | Hohenlohe et al 2010 PLoS Genet |
| groupVII:17150000-17200000 | Alaska | Hohenlohe et al 2010 PLoS Genet |
| groupVII:17700000-17750000 | Alaska | Hohenlohe et al 2010 PLoS Genet |
| groupVII:17900000-17950000 | Alaska | Hohenlohe et al 2010 PLoS Genet |
| groupVII:18950000-19000000 | Alaska | No |
| groupVII:19200000-19250000 | Alaska | No |
| groupVII:19850000-19900000 | Alaska | No |
| groupVII:20050000-20100000 | Alaska | No |
| groupVIII:8350000-8400000 | Alaska | Hohenlohe et al 2010 PLoS Genet |
| groupVIII:8400000-8450000 | Alaska | Hohenlohe et al 2010 PLoS Genet |
| groupVIII:9600000-9650000 | Alaska | No |
| groupXI:5600000-5650000 | Alaska | Hohenlohe et al 2010 PLoS Genet |
| groupXI:6300000-6350000 | Alaska | No |
| groupXI:6550000-6600000 | Alaska | No |
| groupXI:8900000-8950000 | Alaska | No |
| groupXI:9100000-9150000 | Alaska | No |
| groupXI:9550000-9600000 | Alaska | No |
| groupXII:12300000-12350000 | Alaska | No |
| groupXII:12400000-12450000 | Alaska | No |
| groupXII:14800000-14850000 | Alaska | No |
| groupXIII:11700000-11750000 | Alaska | No |
| groupXIII:11800000-11850000 | Alaska | No |
| groupXIII:12950000-13000000 | Alaska | No |
| groupXX:6200000-6250000 | Alaska | No |
| groupXX:8100000-8150000 | Alaska | No |
| groupXX:9100000-9150000 | Alaska | No |
| groupXX:12350000-12400000 | Alaska | No |
| groupXX:12500000-12550000 | Alaska | No |
| groupXX:12550000-12600000 | Alaska | No |
| groupXXI:2850000-2900000 | Alaska | No |
| groupXXI:5550000-5600000 | Alaska | Hohenlohe et al 2010 PLoS Genet |
| scaffold-173:100000-150000 | Alaska | No |
| scaffold-47:150000-200000 | Alaska | No |
| groupII:6150000-6200000 | BC | No |
| groupIX:9950000-10000000 | BC | No |
| groupV:3850000-3900000 | BC | No |
| groupV:4300000-4350000 | BC | No |
| groupVII:21300000-21350000 | BC | No |
| groupVII:21800000-21850000 | BC | No |
| groupVIII:11750000-11800000 | BC | No |
| groupXI:5350000-5400000 | BC | No |
| groupXI:5850000-5900000 | BC | Hohenlohe et al 2010 PLoS Genet |
| groupXI:7750000-7800000 | BC | No |
| groupXII:950000-1000000 | BC | No |
| groupXIII:1950000-2000000 | BC | No |
| groupXIII:8450000-8500000 | BC | No |
| groupXIII:10650000-10700000 | BC | No |
| groupXIII:13250000-13300000 | BC | No |
| groupXV:9850000-9900000 | BC | No |
| groupXVI:16500000-16550000 | BC | No |
| groupXVI:16900000-16950000 | BC | No |
| groupXX:7000000-7050000 | BC | No |
| groupXXI:2650000-2700000 | BC | No |
| groupXXI:2800000-2850000 | BC | No |
| scaffold-37:350000-400000 | BC | No |
| scaffold-37:1550000-1600000 | BC | No |
| scaffold-56:850000-900000 | BC | No |
| groupI:21650000-21700000 | Iceland | Jones et al 2012 Nature/Hohenlohe et al 2010 PLoS Genet |
| groupI:24400000-24450000 | Iceland | No |
| groupIV:22600000-22650000 | Iceland | No |
| groupIV:26000000-26050000 | Iceland | No |
| groupIX:19850000-19900000 | Iceland | No |
| groupXIV:11250000-11300000 | Iceland | No |
| groupXVI:16450000-16500000 | Iceland | No |
| groupXVIII:11700000-11750000 | Iceland | No |
| groupXXI:5750000-5800000 | Iceland | Hohenlohe et al 2010 PLoS Genet |
| groupXXI:6200000-6250000 | Iceland | Hohenlohe et al 2010 PLoS Genet |
| groupXXI:6400000-6450000 | Iceland | Hohenlohe et al 2010 PLoS Genet |
| groupXXI:6700000-6750000 | Iceland | Hohenlohe et al 2010 PLoS Genet |
| groupXXI:6850000-6900000 | Iceland | Hohenlohe et al 2010 PLoS Genet |
| groupXXI:7000000-7050000 | Iceland | Hohenlohe et al 2010 PLoS Genet |
| groupXXI:7050000-7100000 | Iceland | Hohenlohe et al 2010 PLoS Genet |
| groupXXI:7250000-7300000 | Iceland | Hohenlohe et al 2010 PLoS Genet |
| scaffold-27:850000-900000 | Iceland | No |
| scaffold-47:1400000-1450000 | Iceland | No |
| groupI:3550000-3600000 | Scotland | No |
| groupII:12400000-12450000 | Scotland | No |
| groupIII:100000-150000 | Scotland | No |
| groupIV:10900000-10950000 | Scotland | No |
| groupIV:16950000-17000000 | Scotland | No |
| groupIV:22700000-22750000 | Scotland | No |
| groupIV:23900000-23950000 | Scotland | Hohenlohe et al 2010 PLoS Genet |
| groupIX:15700000-15750000 | Scotland | No |
| groupVII:10550000-10600000 | Scotland | No |
| groupVII:19250000-19300000 | Scotland | No |
| groupX:10250000-10300000 | Scotland | No |
| groupXIV:5400000-5450000 | Scotland | No |
| groupXIV:8850000-8900000 | Scotland | No |
| groupXV:4750000-4800000 | Scotland | No |
| groupXX:150000-200000 | Scotland | No |
| groupXX:3600000-3650000 | Scotland | No |
| groupXXI:7950000-8000000 | Scotland | No |
| scaffold-108:0-50000 | Scotland | No |
| scaffold-37:2500000-2550000 | Scotland | No |

|  |  |  | Environmental Variables | | | | |  | Body shape | |  | Armour traits | | | |  |  |  | Trophic traits | |
| --- | --- | --- | --- | --- | --- | --- | --- | --- | --- | --- | --- | --- | --- | --- | --- | --- | --- | --- | --- | --- |
| Traits/ variables |  | MxF | Ca | Gyro | Na | pH | Schisto | Zn | ShPC1 | ShPC2 | ShPC3 | DS1 | DS2 | PS | LP | HP | BAP | Plate_N | Gill L | Gill N |
|  | Marine - Fresh | 21 | 2 | 0 | 1 | 2 | 0 | 0 | 0 | 0 | 0 | 0 | 0 | 1 | 1 | 1 | 0 | 1 | 0 | 1 |
| Environment | Ca | 0 | 14 | 0 | 1 | 3 | 0 | 0 | 0 | 0 | 0 | 1 | 1 | 0 | 1 | 0 | 0 | 2 | 0 | 0 |
|  | Gyro | 0 | 0 | 4 | 0 | 0 | 0 | 0 | 0 | 0 | 0 | 0 | 0 | 0 | 0 | 0 | 0 | 0 | 0 | 0 |
|  | Na | 0 | 0 | 0 | 4 | 1 | 0 | 0 | 0 | 0 | 0 | 0 | 0 | 0 | 1 | 0 | 0 | 1 | 0 | 0 |
|  | pH | 0 | 0 | 0 | 0 | 9 | 0 | 0 | 0 | 0 | 0 | 0 | 0 | 0 | 1 | 0 | 0 | 1 | 0 | 0 |
|  | Schisto | 0 | 0 | 0 | 0 | 0 | 5 | 0 | 0 | 1 | 0 | 1 | 1 | 0 | 0 | 1 | 0 | 0 | 0 | 0 |
|  | Zn | 0 | 0 | 0 | 0 | 0 | 0 | 3 | 0 | 0 | 0 | 0 | 0 | 0 | 0 | 0 | 0 | 0 | 0 | 0 |
| Body Shape | Shape_PC1 | 0 | 0 | 0 | 0 | 0 | 0 | 0 | 2 | 0 | 0 | 0 | 1 | 0 | 0 | 0 | 0 | 0 | 0 | 0 |
|  | Shape_PC2 | 0 | 0 | 0 | 0 | 0 | 0 | 0 | 0 | 3 | 0 | 1 | 1 | 0 | 0 | 1 | 0 | 0 | 0 | 0 |
|  | Shape_PC3 | 0 | 0 | 0 | 0 | 0 | 0 | 0 | 0 | 0 | 2 | 0 | 0 | 0 | 0 | 0 | 0 | 0 | 0 | 0 |
| Armour | DS1 | 0 | 0 | 0 | 0 | 0 | 0 | 0 | 0 | 0 | 0 | 4 | 2 | 0 | 0 | 1 | 0 | 0 | 0 | 0 |
|  | DS2 | 0 | 0 | 0 | 0 | 0 | 0 | 0 | 0 | 0 | 0 | 0 | 7 | 0 | 0 | 1 | 0 | 0 | 0 | 0 |
|  | PS | 0 | 0 | 0 | 0 | 0 | 0 | 0 | 0 | 0 | 0 | 0 | 0 | 11 | 1 | 2 | 0 | 0 | 0 | 1 |
|  | LP | 0 | 0 | 0 | 0 | 0 | 0 | 0 | 0 | 0 | 0 | 0 | 0 | 0 | 10 | 1 | 0 | 1 | 0 | 0 |
|  | HP | 0 | 0 | 0 | 0 | 0 | 0 | 0 | 0 | 0 | 0 | 0 | 0 | 0 | 0 | 3 | 0 | 0 | 0 | 0 |
|  | BAP | 0 | 0 | 0 | 0 | 0 | 0 | 0 | 0 | 0 | 0 | 0 | 0 | 0 | 0 | 0 | 0 | 0 | 0 | 0 |
|  | Plate_N | 0 | 0 | 0 | 0 | 0 | 0 | 0 | 0 | 0 | 0 | 0 | 0 | 0 | 0 | 0 | 0 | 11 | 0 | 0 |
| Trophic | Gill_Raker_L | 0 | 0 | 0 | 0 | 0 | 0 | 0 | 0 | 0 | 0 | 0 | 0 | 0 | 0 | 0 | 0 | 0 | 3 | 0 |
|  | Gill_Raker_N | 0 | 0 | 0 | 0 | 0 | 0 | 0 | 0 | 0 | 0 | 0 | 0 | 0 | 0 | 0 | 0 | 0 | 0 | 9 |

Supplementary Table 12. Summary table with total number of 50kb windows associated with variables in 2 or more radiations. Numbers in the diagonal (scaled green [low] to red [high]) are numbers of genes associated with the same variable (i.e. parallel windows). Table also highlights the number of windows associated with multiple variables (scaled blue [low] to red [high]).

Supplementary Table 13. Associated regions of the genome across environmental variables in more than one radiation. Regions here have been pooled if associated windows were adjacent but associated in different radiations.

| Variable | Window ID | Radiations | Previous Marine - Fresh evidence? |
| --- | --- | --- | --- |
| Ca | groupI:21550000-21800000 | Alaska & Iceland & Scotland | Jones et al 2012 Nature/Hohenlohe et al 2010 PLoS Genet |
| Ca | groupIII:15000000-15100000 | Alaska & BC | No |
| Ca | groupIX:14800000-14850000 | BC & Scotland | No |
| Ca | groupVII:14750000-14800000 | Alaska & BC | Hohenlohe et al 2010 PLoS Genet |
| Ca | groupVII:17800000-17950000 | Alaska & BC | Hohenlohe et al 2010 PLoS Genet |
| Ca | groupVIII:9400000-9450000 | Alaska & Iceland | No |
| Ca | groupX:3050000-3100000 | Alaska & BC | No |
| Ca | groupXI:9000000-9100000 | Alaska & BC | No |
| Ca | groupXII:18050000-18150000 | BC & Scotland | No |
| Ca | groupXVI:13850000-13900000 | Alaska & BC | No |
| Ca | groupXX:150000-200000 | Alaska & Iceland | No |
| Ca | groupXX:12900000-12950000 | Alaska & BC | No |
| Ca | groupXXI:7900000-7950000 | Alaska & Scotland | No |
| Gyro | groupIII:11600000-11700000 | BC & Iceland | No |
| Gyro | groupVII:7800000-7850000 | BC & Scotland | No |
| Gyro | groupVII:7950000-8000000 | Alaska & BC | No |
| Gyro | groupVII:21300000-21350000 | Alaska & BC | No |
| Gyro | groupVIII:10350000-10450000 | Alaska & Iceland | No |
| Gyro | groupVIII:19250000-19300000 | BC & Iceland | No |
| Na | groupI:21550000-21800000 | Iceland & Scotland | Jones et al 2012 Nature/Hohenlohe et al 2010 PLoS Genet |
| Na | groupIII:50000-150000 | Alaska & Iceland | No |
| Na | groupVII:17900000-17950000 | Alaska & BC | Hohenlohe et al 2010 PLoS Genet |
| Na | groupVIII:8250000-8300000 | BC & Scotland | Hohenlohe et al 2010 PLoS Genet |
| Na | groupXII:12150000-12250000 | Alaska & Scotland | No |
| Na | groupXII:13050000-13250000 | Alaska & BC | No |
| Na | groupXX:8950000-9050000 | Alaska & BC & Iceland | No |
| pH | groupI:8550000-8650000 | Alaska & BC | No |
| pH | groupI:21550000-21800000 | Iceland & Scotland | Jones et al 2012 Nature/Hohenlohe et al 2010 PLoS Genet |
| pH | groupIV:22100000-22250000 | Alaska & BC | No |
| pH | groupIX:8400000-8450000 | BC & Iceland | No |
| pH | groupIX:9300000-9350000 | BC & Iceland | No |
| pH | groupVI:7650000-7750000 | Alaska & BC | No |
| pH | groupVI:8000000-8050000 | BC & Scotland | No |
| pH | groupVI:13950000-14000000 | Iceland & Scotland | No |
| pH | groupVII:26900000-26950000 | Iceland & Scotland | No |
| pH | groupXVIII:12200000-12300000 | BC & Iceland | No |
| pH | groupXX:11700000-11800000 | BC & Iceland | No |
| pH | groupXXI:2700000-2750000 | Alaska & BC | No |
| pH | groupXXI:7900000-8000000 | Alaska & Scotland | No |
| Schisto | groupIII:11550000-11650000 | Alaska & Iceland | No |
| Schisto | groupIX:8550000-8600000 | BC & Iceland | No |
| Schisto | groupIX:14250000-14300000 | BC & Scotland | No |
| Schisto | groupIX:14650000-14750000 | Alaska & Scotland | No |
| Schisto | groupX:9250000-9300000 | Alaska & BC | No |
| Schisto | groupXVII:6550000-6650000 | Alaska & Iceland | No |
| Schisto | groupXVIII:6450000-6550000 | BC & Scotland | No |
| Schisto | groupXX:3600000-3650000 | BC & Iceland | No |
| Schisto | scaffold_27:400000-450000 | Alaska & Iceland | No |
| Zn | groupI:8550000-8600000 | Alaska & BC | No |
| Zn | groupIV:19650000-19750000 | BC & Iceland | Hohenlohe et al 2010 PLoS Genet |
| Zn | groupIV:21200000-21300000 | Alaska & Iceland | No |
| Zn | groupVII:6700000-6750000 | BC & Scotland | No |
| Zn | groupXVII:4650000-4750000 | Iceland & Scotland | No |
| Zn | groupXVIII:9750000-9850000 | Alaska & Iceland | No |
| Shape_PC1 | groupIV:13000000-13050000 | Alaska & Scotland | No |
| Shape_PC1 | groupXI:650000-750000 | Alaska & BC | No |
| Shape_PC1 | scaffold_114:300000-350000 | BC & Iceland | No |
| Shape_PC2 | groupIII:8900000-9000000 | BC & Scotland | No |
| Shape_PC2 | groupIX:8550000-8600000 | Alaska & BC | No |
| Shape_PC2 | groupV:9800000-9850000 | BC & Iceland | No |
| Shape_PC2 | groupVI:8700000-8750000 | Iceland & Scotland | No |
| Shape_PC2 | groupXII:12900000-13000000 | Alaska & Iceland | No |
| Shape_PC2 | groupXIII:3750000-3850000 | BC & Iceland | No |
| Shape_PC3 | groupIX:17200000-17300000 | Iceland & Scotland | No |
| Shape_PC3 | groupVII:6250000-6300000 | Alaska & BC | No |
| Shape_PC3 | groupVII:19600000-19650000 | BC & Iceland | No |
| DS1 | groupII:12500000-12600000 | Alaska & BC | No |
| DS1 | groupIII:15000000-15050000 | BC & Iceland | No |
| DS1 | groupIV:12150000-12300000 | Alaska & Iceland | No |
| DS1 | groupIV:17950000-18050000 | Alaska & BC | No |
| DS1 | groupIV:18450000-18550000 | Alaska & BC | No |
| DS1 | groupIX:8550000-8600000 | Alaska & Scotland | No |
| DS1 | groupVII:11200000-11250000 | Iceland & Scotland | No |
| DS1 | groupVIII:5400000-5500000 | Alaska & BC | No |
| DS1 | groupXI:6250000-6350000 | Alaska & BC | No |
| DS1 | groupXIII:11850000-11900000 | BC & Scotland | No |
| DS1 | groupXXI:6400000-6500000 | Alaska & BC | Hohenlohe et al 2010 PLoS Genet |
| DS1 | scaffold_27:1600000-1700000 | BC & Iceland | No |
| DS2 | groupI:7850000-7950000 | Alaska & BC | No |
| DS2 | groupIII:15000000-15050000 | Iceland & Scotland | No |
| DS2 | groupIV:7850000-7900000 | Iceland & Scotland | No |
| DS2 | groupIV:12250000-12300000 | Alaska & BC | No |
| DS2 | groupIV:12950000-13050000 | BC & Iceland & Scotland | Hohenlohe et al 2010 PLoS Genet |
| DS2 | groupIV:18450000-18550000 | Alaska & BC | No |
| DS2 | groupIX:8550000-8600000 | Alaska & Scotland | No |
| DS2 | groupXI:7250000-7300000 | BC & Iceland | No |
| DS2 | groupXX:12500000-12550000 | BC & Iceland | No |
| DS2 | scaffold_27:1500000-1700000 | BC & Iceland | No |
| PS | groupI:7400000-7600000 | Alaska & BC & Iceland | No |
| PS | groupI:11850000-11900000 | BC & Scotland | No |
| PS | groupI:12200000-12250000 | BC & Iceland | No |
| PS | groupII:11550000-11600000 | Alaska & BC | No |
| PS | groupII:14950000-15000000 | Alaska & BC | No |
| PS | groupIII:11600000-11700000 | Alaska & BC | No |
| PS | groupIV:12200000-12300000 | BC & Iceland | No |
| PS | groupIV:15150000-15200000 | BC & Scotland | No |
| PS | groupIV:19850000-19900000 | Alaska & BC | Jones et al 2012 Nature/Hohenlohe et al 2010 PLoS Genet |
| PS | groupVII:13500000-13600000 | Alaska & BC | No |
| PS | groupVII:16350000-16400000 | BC & Scotland | No |
| PS | groupX:14450000-14550000 | Alaska & Scotland | No |
| PS | groupXI:6000000-6100000 | Alaska & Iceland | Hohenlohe et al 2010 PLoS Genet |
| PS | groupXII:14200000-14350000 | Alaska & BC | No |
| PS | groupXVII:5400000-5450000 | BC & Iceland | No |
| PS | groupXX:12500000-12600000 | Alaska & BC | No |
| PS | groupXX:15750000-15800000 | Alaska & Iceland | No |
| PS | groupXXI:6400000-6500000 | Alaska & BC | Hohenlohe et al 2010 PLoS Genet |
| PS | scaffold_48:650000-750000 | Alaska & Iceland | No |
| LP | groupI:21550000-21800000 | Alaska & Iceland | Jones et al 2012 Nature/Hohenlohe et al 2010 PLoS Genet |
| LP | groupIII:11600000-11700000 | Alaska & BC & Iceland | No |
| LP | groupIV:12250000-12400000 | BC & Iceland & Scotland | No |
| LP | groupIV:14400000-14450000 | BC & Iceland & Scotland | No |
| LP | groupIV:19850000-19900000 | Alaska & BC | Jones et al 2012 Nature/Hohenlohe et al 2010 PLoS Genet |
| LP | groupIV:20000000-20050000 | Alaska & Scotland | Hohenlohe et al 2010 PLoS Genet |
| LP | groupIV:27250000-27300000 | Alaska & Scotland | No |
| LP | groupV:6300000-6350000 | Alaska & Iceland | No |
| LP | groupVII:13650000-13750000 | Alaska & BC | No |
| LP | groupVII:15450000-15550000 | Alaska & BC | No |
| LP | groupXI:6000000-6100000 | Alaska & BC | Hohenlohe et al 2010 PLoS Genet |
| LP | groupXII:14150000-14350000 | Alaska & BC & Iceland | No |
| LP | groupXIII:15550000-15650000 | Alaska & BC | No |
| LP | groupXXI:6400000-6500000 | Alaska & BC | Hohenlohe et al 2010 PLoS Genet |
| LP | groupXXI:6550000-6650000 | Alaska & BC | Hohenlohe et al 2010 PLoS Genet |
| HP | groupIV:12950000-13050000 | Alaska & BC | Hohenlohe et al 2010 PLoS Genet |
| HP | groupIV:19850000-19900000 | Alaska & BC | Jones et al 2012 Nature/Hohenlohe et al 2010 PLoS Genet |
| HP | groupIX:8550000-8600000 | Iceland & Scotland | No |
| HP | groupVII:18250000-18350000 | Alaska & BC | Hohenlohe et al 2010 PLoS Genet |
| HP | groupXVII:5400000-5450000 | BC & Iceland | No |
| HP | groupXVII:12500000-12600000 | Alaska & Scotland | No |
| BAP | groupII:5550000-5650000 | Alaska & Scotland | No |
| BAP | groupVIII:9750000-9850000 | BC & Scotland | No |
| Plate_N | groupI:11850000-12050000 | Alaska & BC | No |
| Plate_N | groupI:21550000-21800000 | Iceland & Scotland | Jones et al 2012 Nature/Hohenlohe et al 2010 PLoS Genet |
| Plate_N | groupIV:9750000-9800000 | Alaska & BC | No |
| Plate_N | groupIV:12750000-12850000 | Alaska & BC & Iceland | Jones et al 2012 Nature/Hohenlohe et al 2010 PLoS Genet |
| Plate_N | groupIV:13450000-13500000 | Alaska & BC | No |
| Plate_N | groupIV:19800000-19950000 | Alaska & BC | Jones et al 2012 Nature/Hohenlohe et al 2010 PLoS Genet |
| Plate_N | groupIV:23900000-24000000 | BC & Iceland | Jones et al 2012 Nature/Hohenlohe et al 2010 PLoS Genet |
| Plate_N | groupIV:25550000-25600000 | Alaska & BC | No |
| Plate_N | groupVII:14300000-14450000 | Alaska & BC | Hohenlohe et al 2010 PLoS Genet |
| Plate_N | groupVII:15450000-15550000 | Alaska & BC | No |
| Plate_N | groupVIII:9400000-9500000 | BC & Iceland | No |
| Plate_N | groupXII:12700000-12800000 | Alaska & BC | No |
| Plate_N | groupXX:150000-200000 | BC & Iceland | No |
| Plate_N | groupXX:6150000-6200000 | Alaska & BC | No |
| Gill_Raker_L | groupIII:5350000-5400000 | Alaska & Iceland | No |
| Gill_Raker_L | groupIV:11800000-11850000 | BC & Scotland | No |
| Gill_Raker_L | groupIX:14500000-14600000 | Alaska & BC | No |
| Gill_Raker_L | groupVII:15000000-15050000 | Alaska & BC | Hohenlohe et al 2010 PLoS Genet |
| Gill_Raker_N | groupIII:10400000-10450000 | Alaska & BC | No |
| Gill_Raker_N | groupIV:13350000-13400000 | Alaska & BC | Hohenlohe et al 2010 PLoS Genet |
| Gill_Raker_N | groupIV:14150000-14250000 | Alaska & BC | No |
| Gill_Raker_N | groupIV:21400000-21500000 | Alaska & BC | No |
| Gill_Raker_N | groupIX:8350000-8450000 | Iceland & Scotland | No |
| Gill_Raker_N | groupVII:14650000-14750000 | Alaska & BC | Hohenlohe et al 2010 PLoS Genet |
| Gill_Raker_N | groupX:14500000-14550000 | Alaska & BC | No |
| Gill_Raker_N | groupXV:7150000-7200000 | Iceland & Scotland | No |
| Gill_Raker_N | groupXXI:6100000-6200000 | Alaska & BC | Hohenlohe et al 2010 PLoS Genet |
| Gill_Raker_N | scaffold_47:100000-200000 | BC & Scotland | No |

Supplementary Table 14. Counts of windows (associated and non-associated with variables) included in the analysis. Columns represent Venn segments denoting windows common to all combinations of radiations.

|  | Window Size | | |  |  |
| --- | --- | --- | --- | --- | --- |
| Groupings | 50 kb | 75 kb | 100 kb | 200 kb | 0.1 cM |
| All 4 | 4868 | 4085 | 3536 | 2163 | 3711 |
| Alaska & BC & Iceland | 244 | 142 | 83 | 20 | 210 |
| Alaska & BC & Scotland | 220 | 141 | 80 | 20 | 187 |
| Alaska & Iceland & Scotland | 150 | 100 | 58 | 18 | 138 |
| BC & Iceland & Scotland | 301 | 177 | 126 | 33 | 215 |
| Alaska & BC | 494 | 274 | 141 | 32 | 498 |
| Alaska & Iceland | 23 | 19 | 6 | 4 | 25 |
| Alaska & Scotland | 24 | 15 | 4 | 2 | 17 |
| BC & Iceland | 42 | 22 | 13 | 2 | 35 |
| BC & Scotland | 59 | 33 | 18 | 5 | 29 |
| Iceland & Scotland | 383 | 216 | 135 | 52 | 361 |
| Alaska | 95 | 51 | 36 | 24 | 83 |
| BC | 158 | 82 | 57 | 12 | 111 |
| Iceland | 94 | 53 | 31 | 19 | 69 |
| Scotland | 92 | 52 | 24 | 7 | 97 |
| Total | 7247 | 5462 | 4348 | 2413 | 5786 |

Supplementary Dataset 1 (separate file)

All Bayenv2 associated windows for window sizes of a)50kb, b)75kb, c)100kb, d)200kb) and e)0.1cM. Chromosome/ scaffold where the window is located, base pair numbers at start and end of the window, Ensmebl ID, the name of the radiation(s) and the associated variable / trait. Full names of traits / variables can be found in Figure 2.

Supplementary Dataset 2 (separate file)

Genes located in Bayenv2 associated windows along with GO information from Biomart. From left to right: Ensembl ID of the gene, gene common name, GO Term, GO name, 50kb window where gene is located, radiation and variables that are associated with windows where gene is located. Full names of traits / variables can be found in Figure 2.

Supplementary Dataset 3 (separate file)

Summary table with details of each individual analysed (from left to right): sample code name, country of collection, year sample was collected in, code name for lake where sample was collected, sex, weight and standard length of each individual, RAD tag code and number of reads retained.
